# Discovery and implications of a novel, visual-attracting trap for wood-boring beetles (Curculionidae: Scolyidini): beetle response behaviors and underlying mechanisms

**DOI:** 10.64898/2026.06.22.733798

**Authors:** Russell F. Mizell

## Abstract

*Xylosandrus crassiusculus* (Motschulsky), the granulate ambrosia beetle, was one of the first highly-destructive ambrosia beetles introduced into the southern U.S in the 1970’s where it was found in South Carolina (Kovach 1986). The Redbay ambrosia beetle, *Xyleborus glabratus* Eichhoff, was first detected in the U.S. in South Georgia in 2002. This beetle and its associated fungi, the laurel wilt fungus *Raffaelea laurelensis* and others have caused substantial destruction to native redbay (*Persea borbonia*) in GA, SC, FL and elsewhere. This beetle-pathogen complex also poses a threat to commercial avocado production in the U.S., Central and South America as well as to valuable other *Persea* spp. and related plants (Laureacea) that are known hosts. As an addition here, 10 years of the spring appearances (Fig.1) of *X. crassiusculus* in North Florida is offered for future comparisons. A second unusual appearance is the finding and working with UV mulch and ethanol, as a surprising attraction of *X. crassiusculus* and other ambrosia beetles including *X. glabratus*. It was also found that the ambrosia beetles do not respond to yellow and green as expected by most. Also, adding burlap was found to be attractive (increases dead and dying appearing trees) as is silver metallic like UV mulch, while camouflage (camo) was found to work like yellow and green. These occurrences led to the invention and development of UV mulch with new traps to better monitor ambrosia beetles. New traps led to new uses for yellow, green and camo to monitor and decrease damage and losses from ambrosia beetles. The data are presented as evaluated and appear in the figures, discussion and a supplemental section.

**Figure 1:**
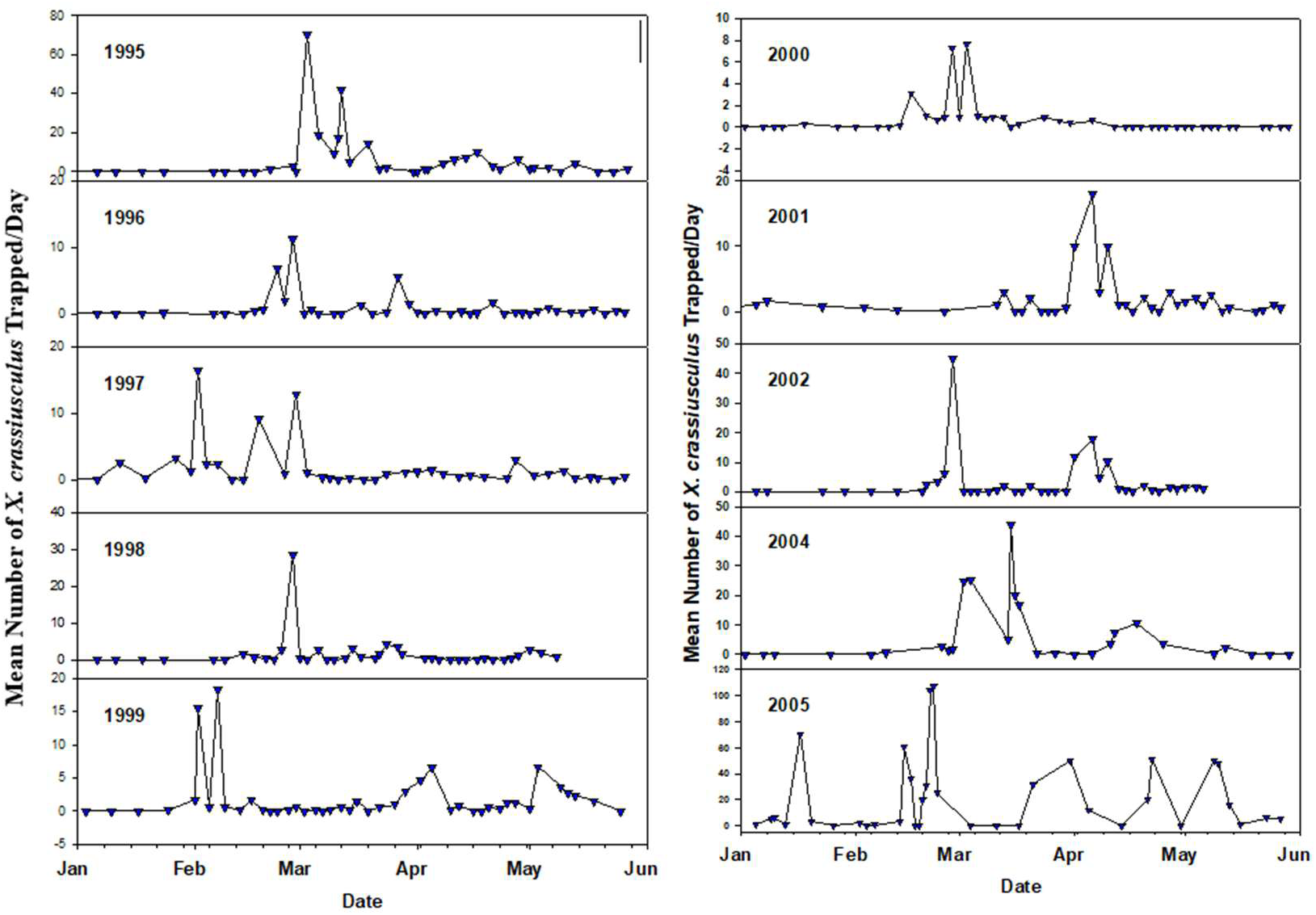
Relative timing of annual emergence of *Xylosandrus crassiusculus* in north Florida. Collected over 10 years using 5 Baker traps with a 10% ethanol/water solution. Data are from years as marked. Note: data from year 2003 was not collected.

## 1. Introduction

Ambrosia and bark beetles in the family Curculionidae are some of the most important insect pests of hardwood and conifer species in forests, nurseries and landscapes. Ambrosia beetles make up around 3400 of the ∼7500 species in the subfamily Scolytinae. Most of them construct tunnel systems in the heartwood of trees (typically in weakened or recently dead trees or, more rarely, in (ostensibly) vigorous hosts; others colonize pith, large seeds, fruits and leaf petioles (Hulcr and Skelton 2015). Ambrosia refers to the fungi cultivated by the beetles on their gallery walls, upon which they feed as an exclusive, or near exclusive food source. The beetles are obligately dependent upon the fungi from which they acquire essential vitamins, amino acids, and sterols (Paine et al. 1997).

Over 3000 species of ambrosia beetles are known and separated into a number of tribes with Xyleborini being one of the most important with ∼1,400 species (Hulcr and Skelton 2015).. In the U.S. there are at least 39 species, of which many are exotic (Rabaglia et al. 2006). The three best known and most economically important pest species are the granulate (formerly Asian) ambrosia beetle (GAB), *Xylosandrus crassiusculus* (Motschulsky) (Mizell and Riddle 2004), the redbay ambrosia beetle, *Xyleborus glabratus* Eichhoff (Mayfield et al. 2008a,b) and *Xylosandrus germanus* (Blandford) (Weber and McPherson 1983, Ranger et al. 2010).

Ambrosia beetles vary widely as a group with respect to host plant species, aggressiveness in attacking ostensibly healthy trees, and in other biology, ecology and behavior. Most exhibit a broad host range and sib mating systems where females mate with their brothers or sons. As their name indicates they transport and transmit from tree to tree a number of species of fungi that colonize host trees. The critical question of whether fungi introduced by colonizing bark beetles are important in killing host trees is open to research but is likely important in the early phases of the interaction (Hulcr and Stelinski 2017).

*Xylosandrus crassiusculus* (GAB) was introduced into South Carolina from Asia and first detected in 1974 (Kovach 1974). It overwinters in all life stages inside the galleries and emergence is dependent upon weather from mid-January to early March in north Florida. They usually attack dormant or “stressed” living trees with roots intact under nursery and landscape conditions. They also attack mimosa (*Albizia julibrissin*), the cut tops of sweet gum and mimosa and pushed-up hardwood trees with roots intact (Mizell, unpublished). During the spring emergence period on into early summer, GAB may be monitored using several types of common wood borer traps (Baker, Kovach, Lindgren, etc.) baited with 95% ethyl alcohol. Traps indicate quite well when GAB emerges in spring and the peak in trap catch accurately reflects times of attack. However, the traps do not always work effectively in mid-summer and fall (Mizell, unpublished). To date no evidence of a GAB pheromone has been demonstrated and this is not unexpected since only the mated females attack new trees. However, the attraction to ethanol suggests that other behaviorally active plant volatiles may be involved in GAB behavior. Ethanol is a known plant volatile produced by “stressed” trees which strongly suggests that tree stress affects GAB host susceptibility. We recently discovered that camphor oil by synergizing ethanol will enhance trap catch by ∼30%, but is not attractive alone (Mizell, unpublished). Miller and Rabaglia (2009) working in several southern states in the U.S. reported that the addition of ∞-pinene along with ethanol increased the attraction of a number of ambrosia beetles including GAB. GAB attack hardwood trees many of which are exotic introductions such as mimosa, crape myrtle, Kwanzan cherry, Drake elm, ‘Bradford’ pear, golden raintree, oaks, and sweet gum. We have also determined that in smaller trees, <5 cm dbh, 3-5 GAB attacks are enough to kill attacked trees (Mizell, unpublished).

The redbay ambrosia beetle, *Xyleborus* glabratus Eichhock an early target of the research presented herein, is native to India (Assam, Bengal), Japan (and the Bonin Islands), Myanmar and Taiwan (Rabaglia 2005). The current range covers most of the southeastern U.S. including most of the coastal and adjacent inland counties in Alabama, Georgia, Mississippi, South Carolina and Florida (Mayfield and Thomas (2006) and now has been detected from Virginia to Texas (Firley 2017). Fungal spores are carried in specialized structures (mycangia) found at the base of each mandible (Fraedrich et *al.* 2008). Brood development can occur in 50-60 days (Hanula et al. 2008). Males of *Xyleborus* species are generally dwarfed, haploid and flightless (Rabaglia 2005). The vascular wilt pathogen (*Raffaelea lauricola*) recovered from infested redbays is vectored by *X. glabratus*. In the U.S., *X. glabratus* has been shown to attack healthy hosts but a low number of species. Although the flight activity of *X. glabratus* tends to be highest in the summer (Hanula *et al*. 2008), adult beetles have been successfully flight trapped in low numbers in north Florida during both winter and summer months (Hudson and Mizell 1999, Mizell unpublished).

Confirmed hosts in the U.S. include redbay, sassafras and avocado. The wilt pathogen associated with *X. glabratus* (*R. lauricola*) has also been recovered from: *Persea americana* (avocado), *P. borbonia* (redbay), *P. palustris* (swampbay), *Sassafras albidum* (Sassafras), *Litsea aestivalis* (pondspice), *Lindera melissifolia* (pondberry) and *Cinnamomum camphora* (Camphor) (Mayfield *et al*. 2008a, Smith *et al*. 2009). Known host trees in Asia include Asian spicebush, *Lindera latifolia*; yellow litsea, *Litsea elongata*; and sal, *Shorea robusta* (Rabaglia 2005). Interestingly, while camphor tree is attacked by *X. glabratus,* the wilt fungus usually does not establish well, and trees are usually not killed (Mizell, unpublished).

Trees attacked by *X. glabratus* exhibit small match-stick-like strings of compacted sawdust protruding from the bark at the point of attack which disintegrate easily and are not always apparent. *X. crassiusculus* and *X. germanus* also exhibit similar behavior. Removal of bark at the point of attack reveals “shot-holes” from which a dark stain extends into the surrounding xylem. The stain is the tree’s response to infection by the *R. lauricola* fungus, which gradually spreads through much of the outer sapwood. Attacked trees eventually exhibit wilted foliage with a reddish or purplish discoloration. Foliar discoloration may occur first within a section of the crown (*e.g.,* major branch) or simultaneously throughout the entire crown. The foliage eventually turns brown and tends to remain on the branches. This wilt scenario is more extensive than the isolated branch “flagging” caused by the black twig borer (*X. compactus* (Eichoff)), which commonly kills twigs and outer portions of small-diameter branches of redbay (Dixon and Woodruff 1982). As the host is attacked, weakens and dies, it is typically colonized by several other species of ambrosia beetles, including *X. crassiusculus*, *Monarthrum mali* (Fitch), *Xyleborus affinus* (Eichhoff) (Foltz et al 2006). Similar associate species and other scolytinae also attack trees that are initially attacked by *X. crassiusculus* (Mizell and M. Thomas, pers. observ.).

Natural enemies of *X. glabratus* and other ambrosia beetles are not widely known. Checker beetles, Cleridae, are common associates with Scolytini of all types and arrive at trees during the attack phase in response to released kairomones (Mizell et al. 1982). Several species of Cleridae have been trapped using sticky traps on trees being attacked by *X. crassiusculus* in Florida (Mizell, unpublished). However, clerid larvae have not been observed feeding in the galleries by the author. During the ambrosia beetle attack phase, semiochemicals are produced that many other scolytinae may use as kairomones. This fact may come into play when attract-and-kill and other semiochemical-based strategies are used to suppress *X. glabratus* (Martini et al. 2015).

According to Hanula et al. (2008) adult *X. glabratus* in Georgia are active throughout the year with peak activity in early September. Brood development takes ∼50-60 days. Wood infested with beetles and infected with the *Raffaelea* sp. is similar in attraction to uninfested redbay wood, whereas both are more attractive than a nonhost species. Sassafras, *S. albidium* (Nutt.) Nees, also a Lauraceae, was not attractive to *X. glabratus* and had few beetle attacks in comparison to redbay. Conversely, avocado, *P. americana* Mill., is as attractive as swampbay, *P. palustris* (Raf.) Sarg., and both are more attractive than the nonhost red maple, *Acer rubrum* L. Trap catches of *X. glabratus* and numbers of entrance holes in redbay trap bolts correlate with the number of dead trees with leaves attached. *X. glabratus* populations drop dramatically after suitable host material is gone. Hanula et al. (2008) suggested that the lack of attraction of *X. glabratus* to sassafras may indicate that spread of *X. glabratus* may slow once it is outside the range of redbay. Cut sections from healthy or diseased redbay trees can be used as baits for trapping. Wounded or cut redbay are attractive to *X. glabratus*, but presence of the fungal symbionts or beetles in the wood does not enhance attraction (Hanula et al. 2008). In a search for attractive baits for *X. glabratus,* Hanula and Sullivan (2008) reported that manuka oil, the essential oil extracted from *Leptospermum scoparium* Forst. and Forst., which contains high proportions of two redbay odor components, α-copaene and calamenene, was attractive to *X. glabratus*. More recently Kendra et al. (2018) reported that the essential oils from redbay, namely cubeb oil plus the addition of (-)-∞-copaene enhanced the attraction of *X. glabratus* to traps.

Given the current pattern of redbay mortality and the rapid range expansion by *X. glabratus*, this non-native ambrosia beetle and its associated fungi have the potential to seriously affect Lauraceous hosts across the southeastern US. Kock and Smith (2008) estimated that *X. glabratus* would spread throughout the range of redbay in <40 years. At the site of the initial *X. glabratus* detection in Florida, redbay mortality increased from 10% to more than 90% in a period of only fifteen months (Fraedrich *et al*. 2008, Mizell, unpublished).

Redbay is important to wildlife as its fruit, seed and/or foliage are eaten by several species of songbirds, wild turkeys, quail, deer, and black bear (Brendemuehl 1990). Larvae of the Palamedes swallowtail (*Papilio palamedes* (Drury)) feed primarily on species of *Persea*, thus this butterfly species is likely to be negatively impacted by *X. glabratus*. There is also considerable concern about the potential impact of laurel wilt on avocado (*P. americana*), which is an economically important crop in Florida. Avocado trees have been found infected with the *R. lauricola* pathogen in the field, and laboratory experiments have shown that *X. glabratus* can successfully bore into avocado trees and introduce the pathogen (Mayfield *et al*. 2008a, 2008b). Apparently, *X. glabratus* do not prefer avocado trees and other ambrosia beetle species have become more important in vectoring the pathogens to avocado in South Florida. Moreover, Menocal et al. (2022) reported that light intensity, canopy cover and other abiotic factors affected the movement and other behaviors differently according to the species of ambrosia beetles.

*X. glabratus* differs from *X. crassiusculus* in having a narrower host range and being more aggressive in attacking larger, ostensibly healthy trees. However*, X. crassiusculus* will attack redbay following attacks by *X. glabratus,* so the two species obviously have some overlap in ecology and behavior in response to tree host plants. Hulcr and Skelton (2017) provided a summary of the relative biology of the above beetles as well as others.

Ambrosia beetles are difficult to control with insecticides for several reasons likely primarily due to lack of substrate exposure or contact (Mizell and Riddle 2004). This contention is supported by differential results from toxicity tests between lab studies in enclosed but ventilated small containers and field evaluations of insecticides (Mizell and Riddle 2004). Because beetles do not consume wood they are not exposed to toxicants through gut poisoning except during the short initiation of galleries when the exterior bark is first removed. Second, before gallery initiation when vulnerable to predation by clerid beetles, the ambrosia beetles spend only a short time period walking on the tree surface where only the extremities of their legs will contact the toxicant; again, reducing overall insecticide contact. Third, they appear to be naturally tolerant of many insecticides (chlorpyrifos, lindane, carbaryl) that are usually effective against other related wood borers and even some *Xyleborus* spp. (Mizell and Riddle 2004, Jagginavar and Naik 2005).

Mayfield et al. (2008c) reported that micro-infusion of the fungicide propiconazole protected redbay from infection by the laurel wilt pathogen when introduced through inoculation. Detection and monitoring of ambrosia beetles currently relies on sticky traps or other commonly used beetle traps (Miller and Rabaglia 2009) which are baited with lures of ethyl alcohol for species that attack trees of host species that are damaged or stressed in some manner such as *X. crassiusculus* and *X. germanus*. A lure for the redbay beetle which attacks more specialized host plants was discovered and optimized (Kendra et al. 2018).

In this publication both old and new research will be presented to improve the understanding of the biology, ecology and behavior of *X. crassiusculus* and *X. glabratus* primarily with other related species also utilized when they occur to strengthen the results. All of the reported research data was gathered in north Florida forested areas in a number of state parks, lands owned by the University of Florida or acreage owned by various private companies including that owned by the author. Moreover, the research was partially funded by a USDA-NIFA competitive grant. All of these are listed in the acknowledgements. An early, unexpected, and very significant discovery of ambrosia beetle behavior occurred and as a result the questions asked, the methodology used to address them, and the results gathered were dramatically changed and also included bark beetle behavior. Therefore, additional inclusion of work that broadened the conclusions and identified new research tools were also included (Conner et. al. 2026). Moreover, the avocado industry has been hurt enough in south Florida to the point of losing $54. mil.

Thus, the objectives are based on the new discovery that ambrosia and bark beetles are attracted to ultraviolet mulch material that is used in vegetable production such as tomatoes, squash and melons for repellency, e.g., for the opposite behavior. This discovery was identified as a very important critical tool to develop and evaluate to promote better understanding of behaviors and management of these pests so old (unpublished) and new critical experiments and outcomes toward these goals were compared and reported with focus on *X. crassiusculus* and *X. glabratus*. Also, a number of new and old tools and support material used by the author or invented are provided, described and compared in the supplementary portion of the manuscript. The latest publication from this work is also now available (Conner et al. 2026). Current damage and continual risks from these two and many related and unknown species have reached the level that warrant research toward improved management. The objectives identified as critical for effective management of these pests and critical outcomes toward these goals are reported with main focus on *X. crassiusculus* and *X. glabratus*.

## Materials, Methods and Results

### Part 1

#### 1.1: 10 years of *Xylosandrus crassiusculus* phenology during spring emergence using the Baker trap

After detection in the southern U.S., *X. crassiusculus* quickly spread around the southeastern states affecting many different host trees in the landscape and crops such as fruits and in nurseries (Coleman et al. 2023). After it became a nursery pest in Florida, research began and early on testing pesticides that would improve manage (Mizell et al. 1998). In the course of that work, using the Baker trap (Frank et al. 2023) with a lure of ethanol was initiated to monitor the early spring emergence flight times of the beetles. In the course of that and other efforts, traps were maintained around local nurseries which were checked for *X. crassiusculus,* the only species collected in meaningful numbers in the early 1990s. Six traps mounted on top of tomato stakes placed in the ground ∼30m apart were placed similarly along the edges of woods where the beetles occurred each year from 1995 until 2006 (2003 was not sampled, thus no data included).

**1.1: Results:** The total number of *X. crassiusculus* collected every 2-3 days are provided from this endeavor (Fig. 1). The data is offered as a baseline indicating the early (January and later) emergence flight behavior of the beetles early in their colonization in the area from year to year. The data is particularly of interest in lieu of the climate change in North Florida in the years since. There are wide differences in the year-to-year data on initial emergence dates, number of beetles collected, when and the relative patterns of trap collections; some with single peaks while other years showing multiple peaks. From a management perspective, this presents a difficult effort to strategically time management tactics to suppress the populations, especially in nurseries and landscapes. Nonetheless, the phenology of this invasive species found in North Florida is not unexpected. In fact, this pattern, 5-7 weeks differences from year to year in the emergence of overwintering adult insects in north Florida and South Georgia, while not a well-known scientific phenomenon, can be readily observed in stink bugs, several pecan insects and many other species (Mizell, unpublished, Grabarczyk et al. 2022).

#### 1.2: Four experiments looking at the impact of DEET and insecticides on *X*. *crassiusculus*

Insecticide efficacies were tested alone and with 40% DEET, the repellent, N,N-diethyl-meta-toluamide (Bayer Crop Science and sold by WPC Brands, Inc., Bridgeton, MO 63044) against *X. crassiusculus.* Four experiments were conducted using bolts from an “occasional” natural host, the non-native mimosa tree (*Albizia julibrissin* Durazz) bolts that were <6cm in diameter x 46cm long. All bolts used were placed with the end from closest to the tree roots in a 10% solution of ethanol and water in a small, cup size, plastic yogurt cup. Insecticides were hand sprayed with the treatments using a small hand-pump sprayer. The 40% DEET was sprayed directly from the original spray can. When DEET was used in combination with an insecticide, it was sprayed over top of the treated insecticide after it had dried. All treatments contained 6 bolts as replications and were set out within 6 randomly placed blocks. Wires were placed between trees low enough to support the bolts that were tied to the cross wire with wire. Two locations for the two treatment groups, e.g., insecticides with and without DEET, were used for the experiments in late June-July, one in Monticello and another in Quincy, FL. The experiments were conducted over the same days at the two locations. Each day after treatments, the bolts used in the treatments were checked for frass boring “sticks” commonly made by attacking *X. crassiusculus* and the evaluations were based only on the recorded number of holes with frass sticks counted/bolt/day. No other species of beetles occurred in the test bolts. The following insecticide treatments and rates were used in the experiments: Untreated control: a bolt placed in 10% ethanol and water solution and 40% DEET as mentioned above alone or with the insecticides. All rates are presented as an amount of insecticide material per 100 gals of water: Discus (2.94% ai imidacloprid and 0.7% ai cyfluthrin) at 100oz, Onyx EC (23.4% ai bifenthrin) at 32oz, Talstar (7.4% ai bifenthrin) at 32oz, Tempo (11.8% ai beta-cyfluthrin) at 1.5oz, and Proclaim (5% ai emamectin benzoate) 1.2oz. One difference in the treatments used in the 2 locations was that Proclaim was used while Talstar was not in the single insecticide group, whereas the opposite was true for the group with DEET combined with the insecticide. This was done because Talstar and Onyx are both bifenthrin but the active ingredients are ∼68% different, see above. Moreover, in a previous experiment that compared the two chemicals, Talstar did not kill the beetles as well as Onyx, and the idea was to see if DEET would help improve beetle mortality from Talstar. Additionally, and earlier, several lower rates of DEET were similarly compared (data not shown) and did not cause equivalent efficacy of 40% DEET on the target species.

**1.2 Results:** The results from the comparison of insecticides, DEET and the individual insecticides combined with DEET, are presented (Fig. 2, 3, 4, 5). The results from the 4 experiments (2/location) indicate that the insecticides and the addition of DEET with them provided different results between locations and treatments. The total *X. crassiusculus* caught was similar in the two locations with beetle collections being in the 15-20 beetles/bolt range across treatments. Moreover, use of DEET caused a reduction in attacking beetles alone or when added to the insecticides (including Talstar but at less than Onyx, the low and high rates, respectively) that displayed the least number of attacking beetles – Tempo and Onyx.

**Figure 2:**
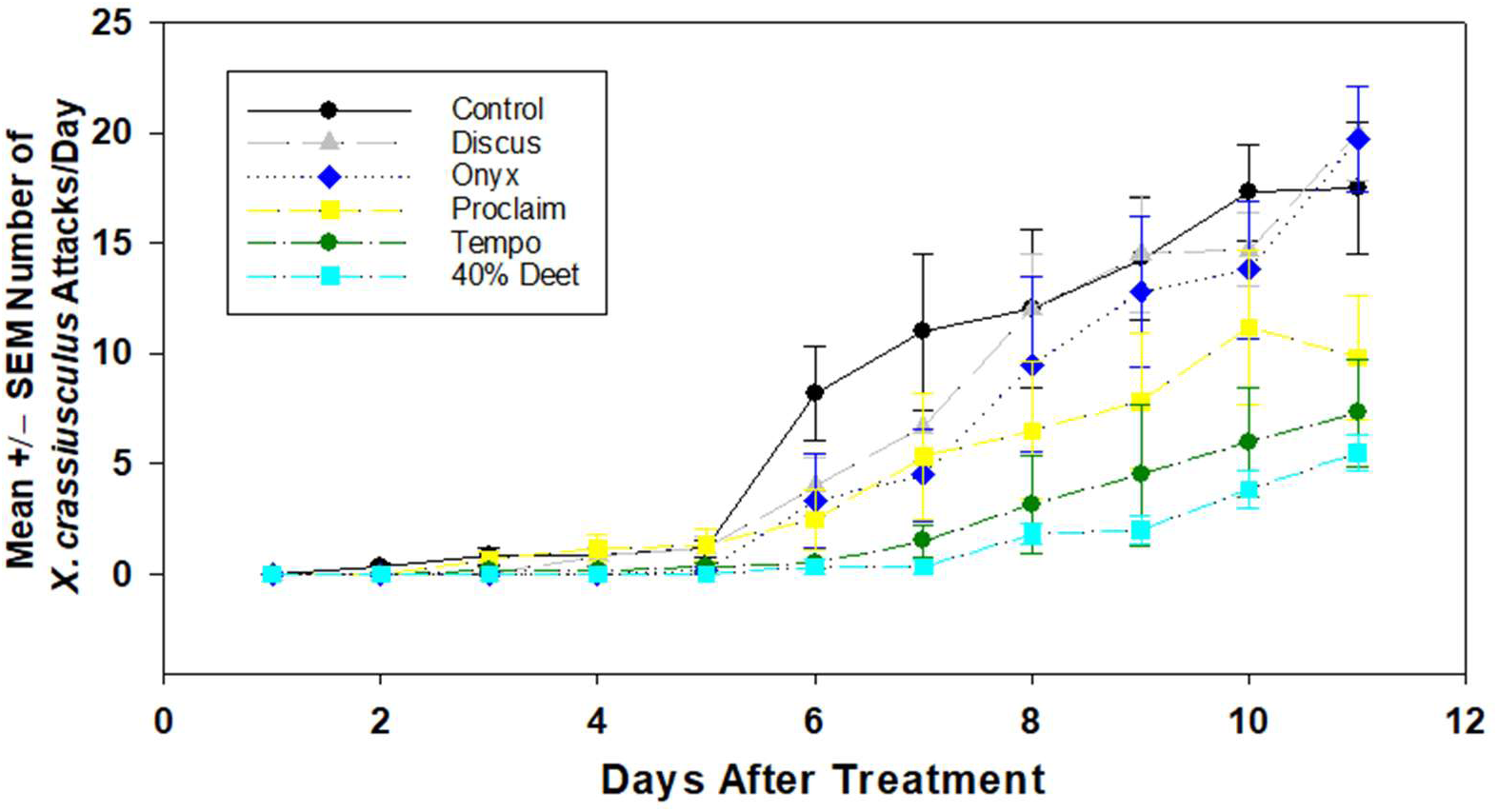
Response of *X. crassiusculus* to insecticides over an eleven day period. Six reps/treatment of mimosa bolts 45cm (18”) long and < 6.5cm (3.0”) diameter in a 10% ethanol/water solution. The low numbers that occurred in the early days were from Quincy, FL and the higher numbers were from the two experiments in Monticello, FL. All rates of the chemicals used were at the middle of the percentage recommended on the labels.

**Figure 3:**
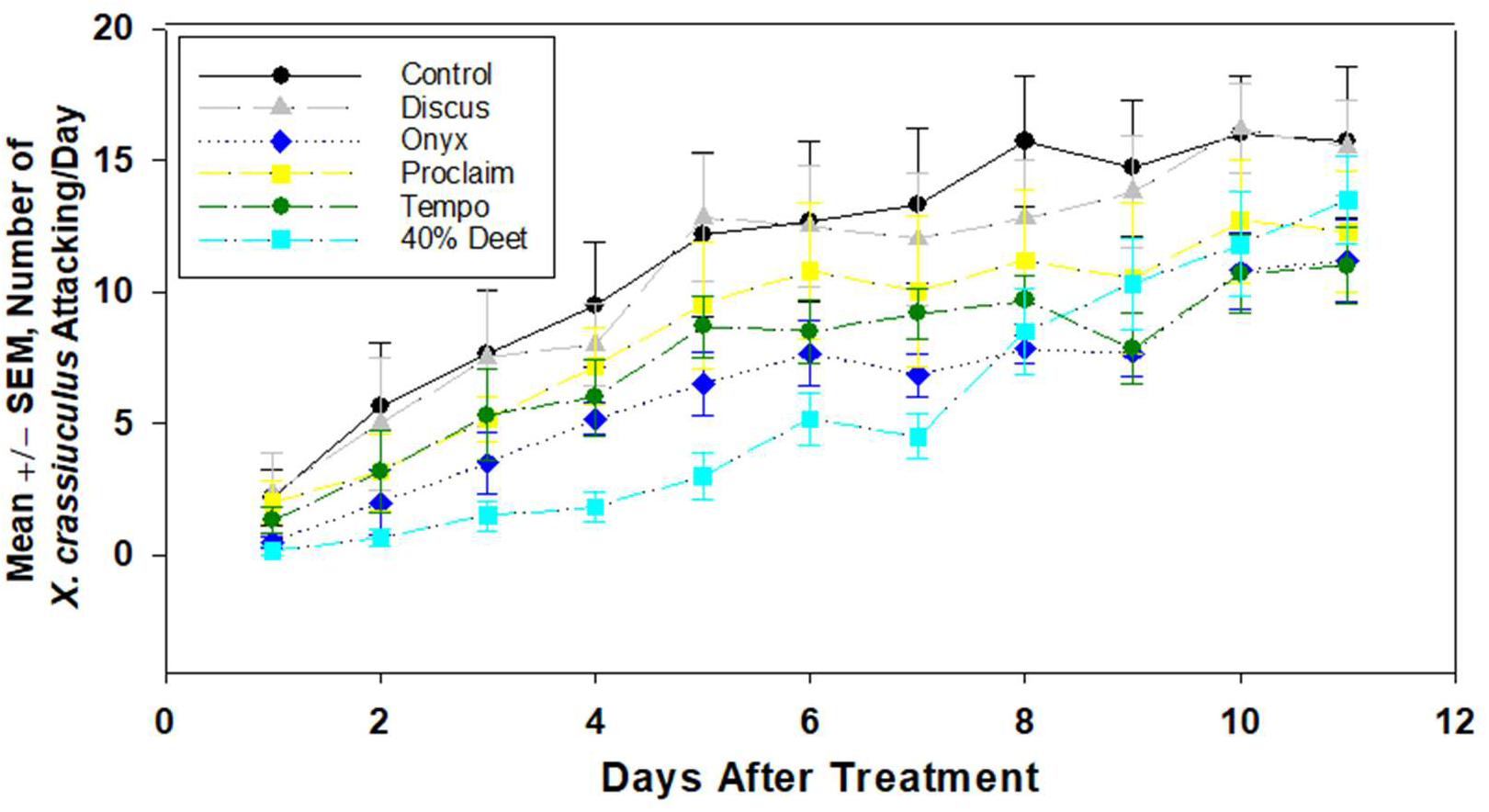
Response of *X. crassiusculus* to insecticides over an eleven day period in Monticello, FL. Six reps/treatment of mimosa bolts of 45cm (18”) long and < 6.5cm (3.0”) diameter in a 10% ethanol/water solution.

**Figure 4:**
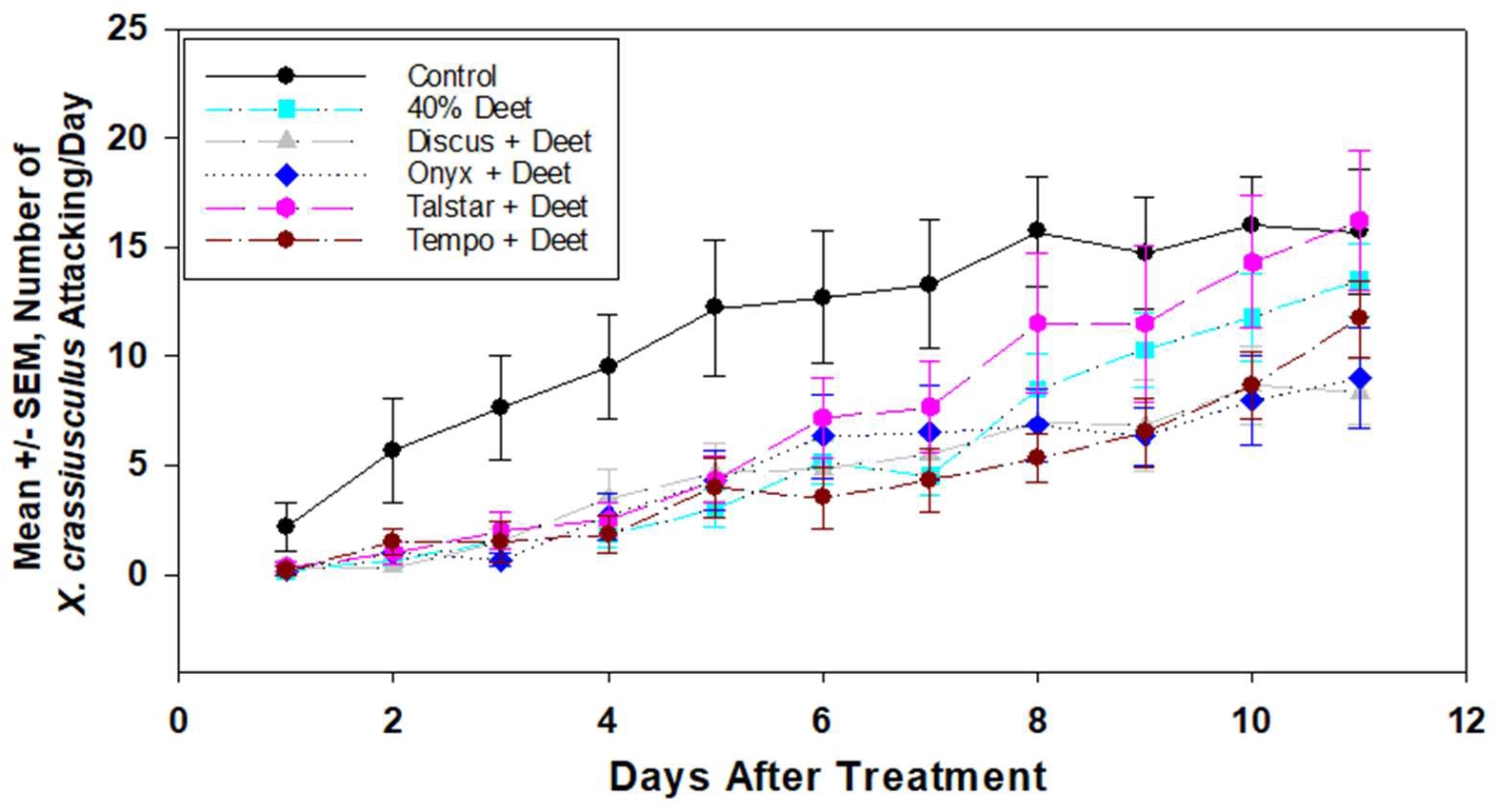
Response of *X. crassiusculus* to insecticides over an eleven day period in Monticello, FL. Six reps/treatment of mimosa bolts of 45cm (18”) long and < 6.5cm (3.0”) diameter in a 10% ethanol/water solution. Note: all 4 runs are same on these.

**Figure 5:**
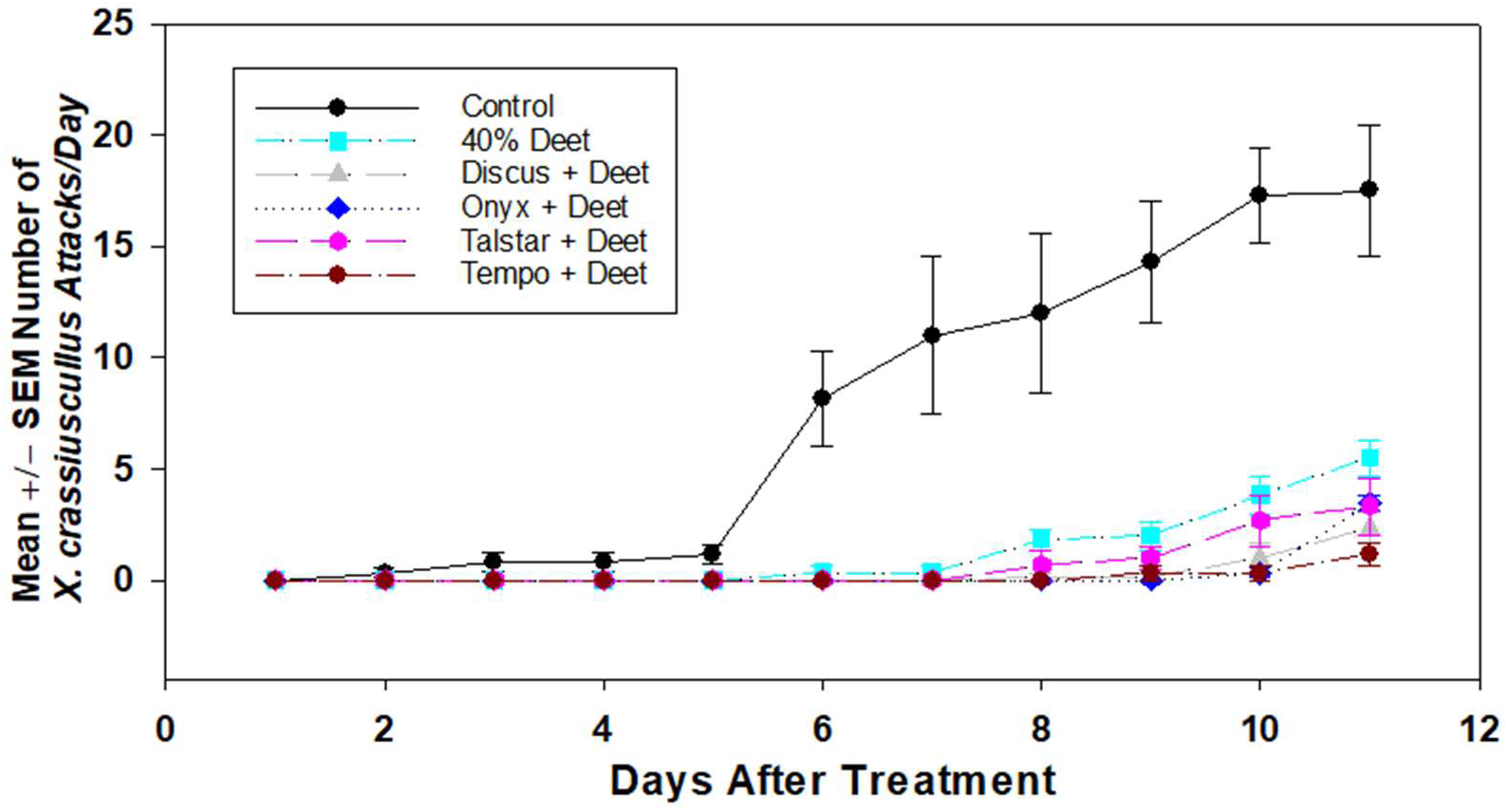
Response of *X. crassiusculus* to insecticides over an eleven day period in Quincy, FL Six reps/treatment of mimosa bolts of 45cm (18”) long and < 6.5cm (3.0”) diameter in a 10% ethanol/water solution. Note all 4 runs are same on these. The early low count days are from Quincy, FL and the higher numbers are from Monticello, FL. Previous results from DEET at lower concentrations were much less effective than a 40% concentration (data not shown).

The Monticello experiment with insecticides only, indicated that DEET lasted as a protective treatment just as long as the insecticides and at numerically lower rates than Tempo and Onyx and much lower than Proclaim, Discus and the control (Fig. 2). The Monticello experiment with insecticides and DEET together (Fig. 3) indicated an increase in efficacy from the mixed chemicals up until the sixth day when beetle numbers started to increase toward the control. Also, the same 3 chemicals, DEET, Tempo and Onyx, provided the best efficacy as without DEET early, but faded in efficacy at 8 days posttreatment (Fig. 4). In the Quincy experiment there were no beetle attacks in the first 5 days post-treatment, but the numbers changed quickly on the sixth day (not sure why the difference from Monticello?) for chemicals but DEET remained lowest with Tempo. Similar efficacious results occurred as in the Monticello experiment, but the attack numbers were much lower when DEET was combined with the insecticides for all of them, whereas without DEET, Tempo and DEET prevailed over the other 3 treatments that rose closer to the control numbers (Fig. 5).

### Part 2: Discovery of unknown behaviors by ambrosia and bark beetles

Topic background. Andersen et al. (2012) provides a summary of the definition, uses and efficacies of polyethylene mulches commonly used in agriculture. There are various types of plastic (or silver) mulches available for use and have been since the 1950’s in vegetable production (Emmert 1957, Andersen et al. 2012). The mechanisms of the different types of mulches are related to the transmission, absorbance, and reflectance of solar radiation. The high ultraviolet-reflecting material is applied over mounded soil at planting time, and then for a period, until the crop plant size has overgrown the material, it effectively disrupts the normal behavior of vegetable pests such as thrips and whiteflies. In this paper, the plastic mulch that is an ultraviolet-reflective metalized mulch (Reflectek Foils, Lake Zurich, IL, 60047) will be referred to as the use of UV mulch and then “UVM” to save space (Andersen et al. 2012). It was originally tested to see if ambrosia beetles were repelled by it as when used in management of whiteflies and thrips in vegetables. The results were the opposite of expectations and were then tested for attraction behavior and underlying mechanisms relative to ambrosia and bark beetles as described below. Before further details are discussed below, 2 spectra of the UVM and related materials are compared as well as a spectrum showing UVM and 4 other materials of interest including white paint and other factors of coroplast discussed below. The reflectance of sunlight by UVM, aluminum foil and aluminum paint are compared (Fig. 6) and that of UVM vs metallic silver paint (Fig 7). The third spectrum (Fig. 8) provides a comparison of light reflection as above to enable discussion of how light is changed dependent upon the materials, which is relevant to ambrosia beetle color behavior discussed next.

**Figure 6:**
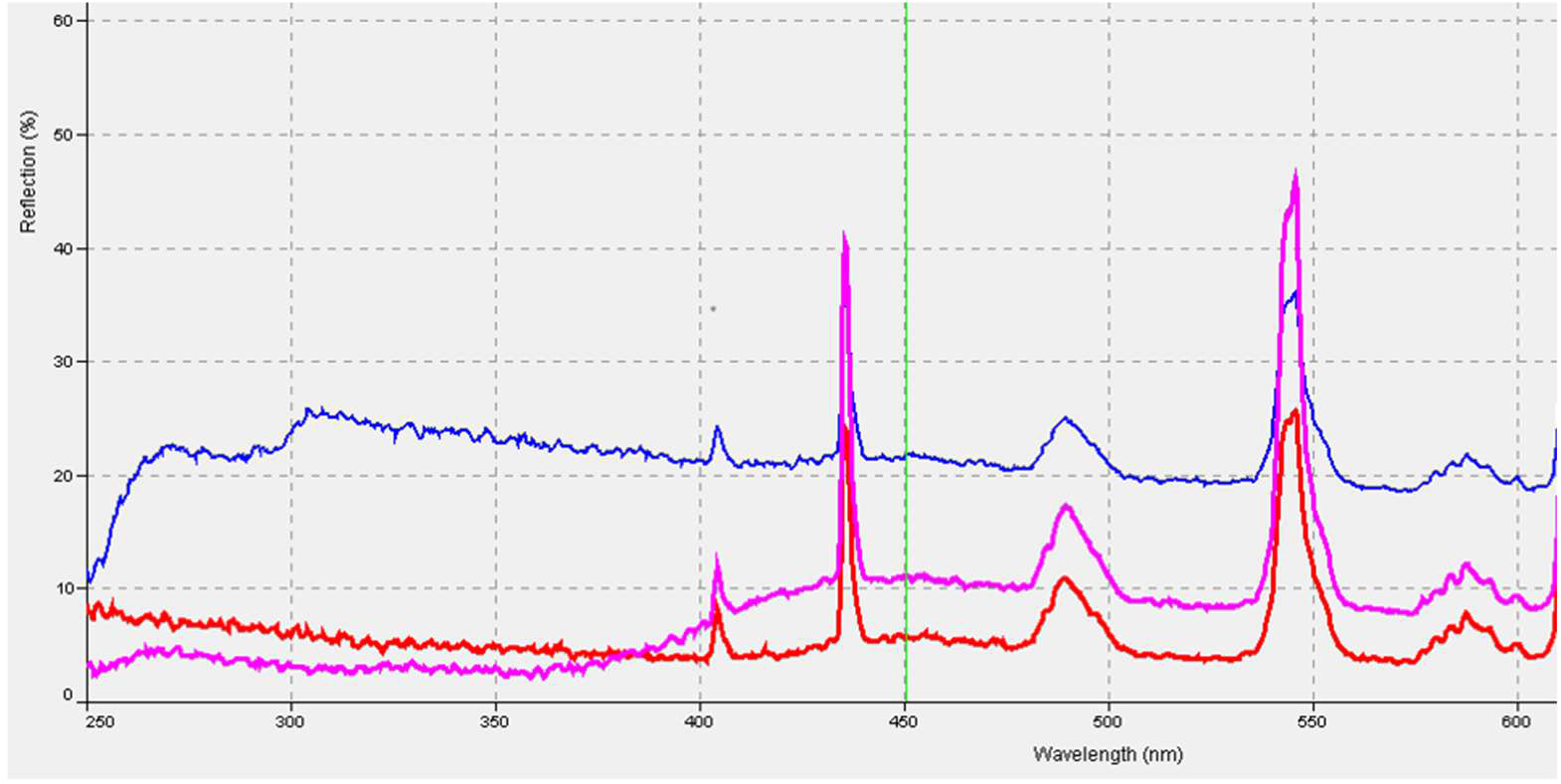
Reflection spectra of UV mulch (bottom and pink) versus aluminum foil (middle and red) and aluminum paint (top and blue). Data collection used with an Ocean 2000 spectrophotometer.

**Figure 7:**
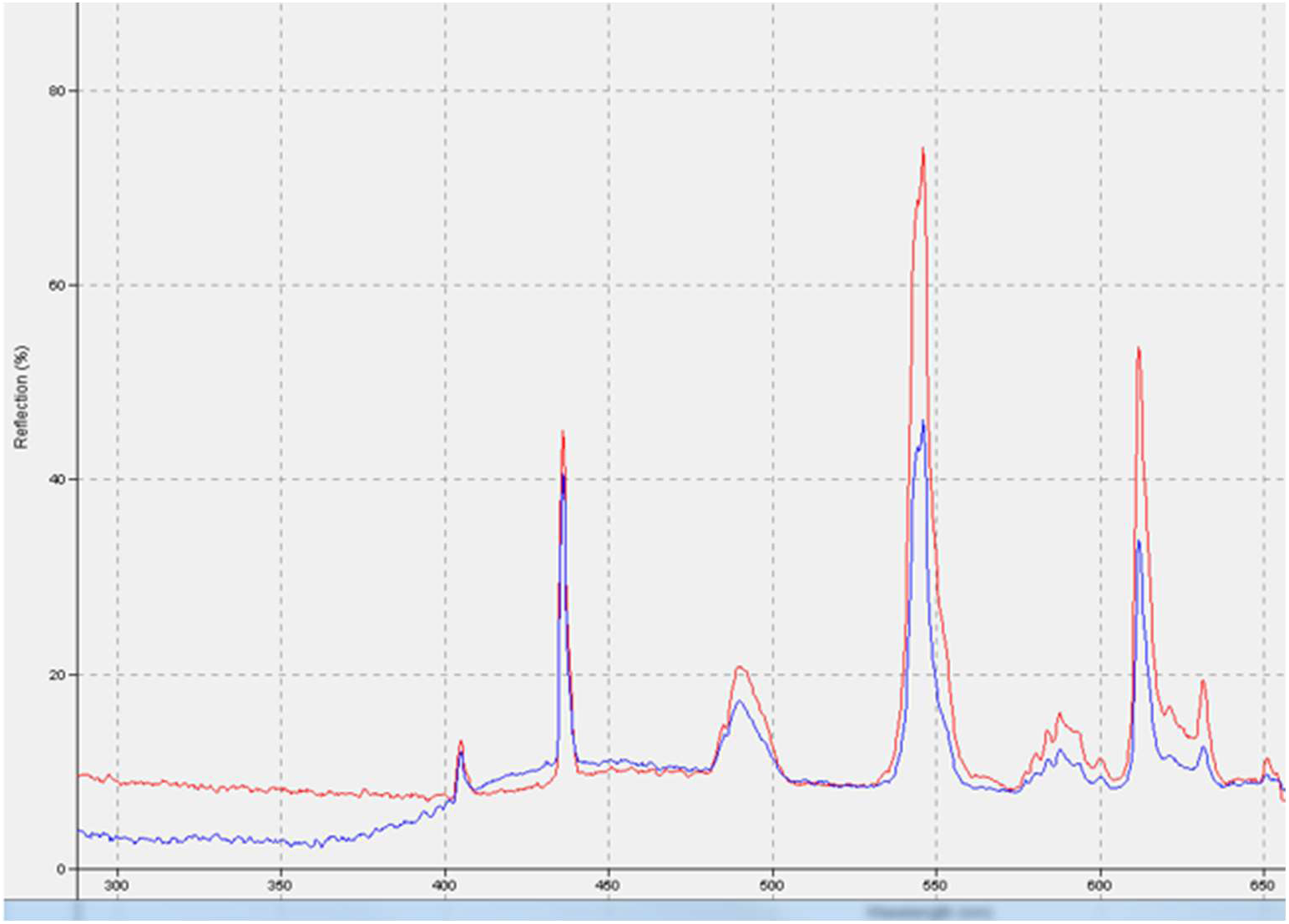
Reflection spectra of UV mulch (bottom blue) versus metallic silver paint (top red). Data collection used with an Ocean 2000 spectrophotometer.

**Figure 8:**
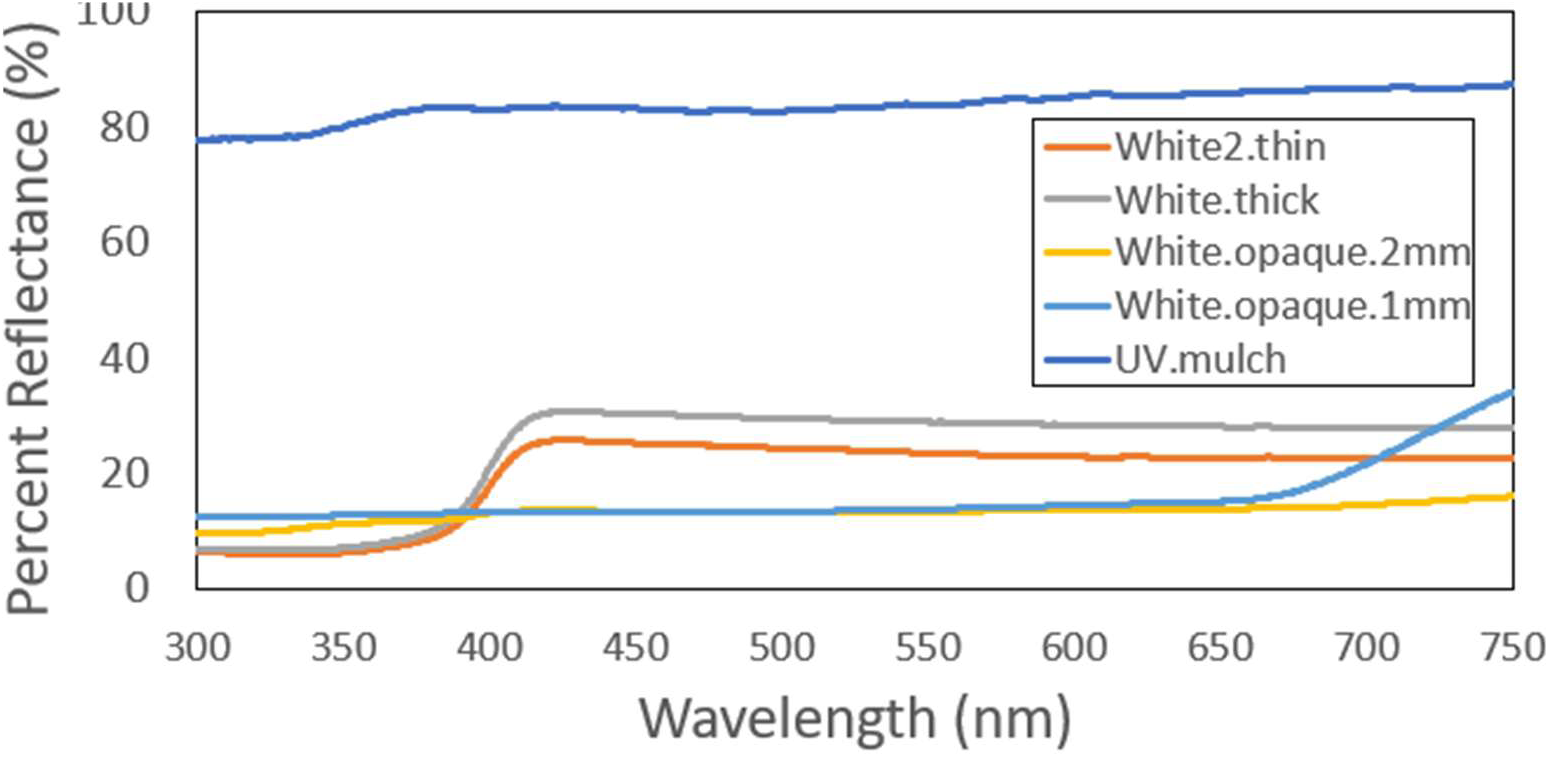
Reflection spectra of white-opaque from 2mm coroplast (bottom – yellow) versus white opaque 2mm coroplast (top blue), white – thin 2mm coroplast (3^rd^ from bottom orange), white 4mm thick coroplast (4^th^ from bottom gray) and UV mulch (bottom and blue). Data collection used a Stellar Net (UV-VIS) Spectrometer.

#### 2.1: Response of *X. glabratus* and *X. crassiusculus* to colors and UVM. This section and more also deal with the redbay beetle, *Xyleborus glabratus* Eichhoff

Color response of *X. glabratus* in a forest in north Florida with small – large host trees of redbay, magnolia, oaks, and several other species were present in the location. Pieces of 4mm coroplast 30.5 x 45.7cm in size were painted different colors or combinations of colors, covered with Tangle-Trap™ (Tangle-Trap™ sticky coating “paste formula”, The Ortho Group, P.O. Box 190, Marysville, OH, 43040) (hereafter Tangle-Trap™) on one side and attached to tree trunks at 1.5 - 2.0m above ground. In a forest with large, different size redbay trees, different sizes were selected in a size range of 10-15cm at DBH and without infestation by ambrosia beetles. Other *X. glabratus* beetle-infested trees were in the area and the traps were baited with ∞-copaene (AgBio, Inc., 9915 Raleigh St, Westminster, Colorado, 80031). Colors tested of 2mm coroplast included UV mulch (specific material discussed above), bright metallic silver (paint similar to the spectra reflected by UV mulch and equal to aluminum foil), paints of black, white, gray, and opaque coroplast. Opaque in this experiment is the grayish and more or less “clear-gray” coroplast available in sheets in 2, 4 or 10mm in thickness.

**2.1 Results:** Bright metallic silver was evaluated in this experiment to determine if *X. glabratus* (and other beetle species as well) would respond to silver paint that could be used as a replacement for the use of UV mulch which is more expensive. The SAS Proc Mix procedure was used to compare the response by *X. glabratus* to the colors. Using a contrast statement indicated that bright silver was significantly different than the other colors with df = 18, F = 12.6, Pr > 0.0023. Differences in Least Squares means indicated that bright silver and opaque collected equal numbers, while white was significantly less than both. In similar comparisons to UV mulch and gray, bright silver did not differ from UV mulch but gray was less than both (t-values of 2.66 (Pr > 0.014, t = 0.014) and Pr > 0.039, t= 2.18, respectively). Response to black as usual was much lower than to both UV mulch and bright silver. UV mulch was originally added to a previous experiment to determine if it might be repellent to ambrosia beetles as it is to such insect pests as whiteflies and thrips. Finding that it was attractive to *X. glabratus* and other ambrosia beetle species was a “revelation” about ambrosia beetle behavior, the potential use of UV mulch, and led to these follow-up experiments. Lastly, the response to the coroplast traps by *X. glabratus* demonstrated a consistent pattern that resulted in a strong “edge effect” in the area on the coroplast of capture of the beetles. This can be interpreted as a visual behavior relying on a contrast between the near and far visual stimuli for orientation to a target attractant.

#### 2.2: A Trap Color Experiment – redbay beetle, *Xyleborus glabratus*

Before this study as stated above, the response of ambrosia beetles (mainly *X. crassiusculus* and *X. glabratus*) to UVM was discovered along with the determination that bright silver paint was equivalent to UVM material for attracting ambrosia beetles. From previous work, it was determined that white and black colors were inferior in attracting ambrosia beetles to traps. That work was done with the redbay beetle and *X. crassiusculus* (if also trapped) using 30.5 x 45.6cm (12” x 18”) coroplast squares of various color combinations on redbay or trunks of other tree species placed at DBH. That methodology removed much of the redbay odor from the color testing and was not in the cylindrical shape of the natural tree trunks. Moreover, it was also determined (data not shown) that: 1. redbay beetles preferred larger trees and more beetles attacked the trees in lower areas closest to the ground, and 2. that 25 - 30.5cm diameter tubes in UVM were equally effective (and more effective than smaller tubes) and the optimum sizes for the new UVM trap. Color comparisons on trees for *X. glabratus* with 5 treatments all on each of 5 trees with same heights (< 2.0m) changed/tree so all treatments were at all 5 heights once among the 5 trees. Clear plastic sheeting was used as the clear treatment and by painting for the white, black and bright silver colors. UV mulch as described above and by Andersen et al. (2012) was used in the original state and bright silver was selected to mimic the reflectance spectrum of UV mulch (see above figures). The UV mulch pieces painted or used were cut to provide 30.5cm length and at width large enough to completely wrap around the trees. Each treatment was covered lightly with Tangle-Trap™ to capture the responding *X. glabratus*. To enhance the attraction of *X. glabratus* and other ambrosia beetles to the 5 trees, a drawknife was used to peel bark off the tree in a small section (∼2.5 x 5cm) under each of the wraps prior to their mounting by stapling. A lure containing α-copaene was placed on a wooden stake ∼1.2m above ground and 1m away from the trees to attract *X. glabratus* to and within the experimental location. All traps were immediately attacked by redbay beetles in the first week post setup.

**2.2 Results:** Proc Mixed analysis (SAS, Inc) and a Type 3 test of Fixed effects indicated height as NS, (df 4, F = 0.6 and P > F = 0.671) but color was significantly different df 4, F = 4.61, Pr > F = 0.0097. Response to all 5 colors via least squares means were significantly > 0 with a total catch of 1778 beetles. Significant difference among the colors via contrast statements was as follows: UV mulch vs the other 4 colors (25.5), F = 12,68, Pr > F=0.0026 following least squares estimates. Using contrast tests UV mulch vs silver (20.5) was not significantly different (F =2.0, Pr >F = 0.19), black vs UV mulch (11.6) was significantly lower (F =15.9, Pr > F = 0.0011. UV mulch and both clear (15) and white (15.7) were not significantly different, although both white and clear numerically were 41% lower than in the catch rate of *X. glabratus*. The results of Tukey’s Studentized Range Test indicated a significant difference between UV mulch and white, black and clear. Silver (Rust-oleum metallic, bright and reflective brand as above), which was a paint compared to the UV mulch material, were not significantly different.

### 2.3: Use of traps of different colors and shapes for comparison of trap catch of *X glabratus* and other ambrosia beetles

In preliminary experiments (after the UVM discovery and before the fake tree experiments, see below) a series of traps were tested in the field to ensure that the color spectrum of light reflected by UVM and its equivalent metallic silver paint both caught ambrosia beetles. Regular Lindgren traps (6 funnels) were also evaluated in the standard colors of white and black along with “homemade” traps that were smaller traps with vanes or tubes placed over a large white 31cm plastic funnel with a collecting jar at the bottom. Traps were placed in the forest and attached to tree limbs so that the 0.5-1.2m space above ground held the trap “entry” areas. “Entry” areas were either vanes 20cm wide x 37 cm in length covered with a lid at the top and attached inside the funnel within the top 5cm, or painted cardboard tubes 15cm in diameter and 37cm in length attached to a lid and funnel as with the vanes. The entire open areas of the vanes, tubes and lids were painted either metallic silver, white or black. The lure α-copaene was placed inside the top of the traps. Four or more reps of each trap type and color were placed in a line at 20-30m apart in blocks containing traps with one of the 3 colors and a Lindgren trap.

**2.3 Results:** In all cases the results (data not shown) indicated that metallic silver collected numerically equal or greater numbers of beetles than similar traps in either black, white or the Lindgren trap. Numbers in vane traps were consistently 25-50% higher than in tube or Lindgren traps. Only one problem was observed: a number of times during the field experiments, the trap funnel was used by birds (wrens) to build nests which resulted in the loss of the beetle count data.

#### 2.4: Evaluation of the UV mulch material for its ability to attract beetles away from hosts (3 different visual approaches) and protect redbay host trees from *X. glabratus* under forest conditions. This is the first test of UV mulch response by ambrosia beetles without lures involved. There are 3 (2.4.1, 2, 3) different experiments in this section

**2.4.1** Attraction of host and non-host trees wrapped in UV mulch: Four trees were used without lures to determine the natural landing rate of the beetles in a stand of hardwood trees. In a moderately dense forest, trees used were an oak, a beach, a southern magnolia (3 non-hosts) and a redbay tree (natural host). The trees were all approximately equal at DBH – 13-18cm (5-8”) and naturally occurring within 5m (13’) of each other. UV mulch from a roll of 1.5m in width was wrapped around tightly and stapled on to each tree to cover the tree boles from the ground to the end of the mulch width. In the middle of each of the trees, Tangle-Trap™ was painted around the UV mulch on the tree in a “belt” ∼ between 0.6 −1.2m, e.g., 0.3m wide in the middle of the wrapped area. The number of *X. glabratus* and *X. crassiusculus* were counted on the tree covered with Tangle trap once per week over 4 weeks. The data were analyzed using Proc ANOVA and Tukey’s Studentized Range test (SAS 9.4).

**2.4.1 Results:** All four trees in an absence of a chemical lure captured beetles of both species. Mean number of *X. glabratus* caught/week were significantly different by Tukey’s studentized range test (P= 0.05, df =12, Minimum significant difference =33.45 by tree type: magnolia mean = 42.5, redbay 22.5, beech 18.8 and oak at 7.75 - magnolia numbers 50% higher - while *X. crassiusculus* numbers were much lower - means of 2.5 vs 7.0 low-high - redbay. Although *X. crassiusculus* was not statistically different, the numerical response of both species was surprising given the absence of lures and the higher numbers on magnolia and redbay by *X. glabratus* and *X. crassiusculus*, respectively. High populations of both species were present at the site and the capture of higher numbers of *X. crassiusculus* on unattacked redbay without lures might suggest that ethanol isn’t the only attractant for *X. crassiusculus*? Numbers: *X. glabratus* (df=3, F value = 3.31, Prob > F = 0.057) means ± SEM/week (n=4 weeks) = oak 7.75 ± 2.53, beech 18.75 ± 1.93, redbay 22.5 ± 10.02, and magnolia 42.5 ± 11.98. *X. crassiusculus* (df=3, F value = 1.71, Prob > F = 0.02), redbay 7.0 ± 2.86, magnolia 2.75 ± 1.49, beech 2.25 ± 1.03, oak 2.0 ± 1.22.

**2.4.2:** Attraction of beetles to eight redbay host trees with 1 “fake tree” wrapped in UV mulch (see below for further explanation on its development) on each side of the centered trees made from 10cm x 1.2m tubes at 2m away, one with a lure and trap, the other without a lure but also a 15 x 30cm trap with Tangle trap (as described above). The 16 fake tree tubes were (4 each) 5, 10, 20 or 25cm in width and all were 1.2m long. The pairs/host trees were made by placing the smallest - largest (based on DBH as above) to the end giving 8 combinations: 5 and 10cm, 5 and 20cm, 5 and 25cm, 10 and 20cm, 10 and 25cm, 20 and 25cm, 5 and 20cm and 10 and 20cm. The host trees also had a UV mulch trap similar to the fake trees at 61cm above ground (see previous experiment). The lure used was α-copaene. Data were collected 6 times from 1 June-10 July at 7 day intervals except for collection 3 which due to bad storms was 13 days apart. The lures were changed to the opposite fake tree without the lure each collection time so that all of the fake trees carried or did not carry the lures 3 times and for all diameters (5 – 25cm) of fake trees. Also, 2 of the host trees’ trunks were also wrapped (under trap) with UV mulch from ground to ∼1.5m. The data were analyzed using Proc ANOVA and Tukey’s Studentized Range test (SAS 9.4).

**2.4.2 Results:** The area of the experiment contained redbay trees of all sizes which were in all stages of attack by ambrosia beetles, mostly *X. glabratus*. Approximately 17,000 total *X. glabratus* were collected on the fake trees and the host trees during the 7-week period. None of the experimental host trees were attacked until the fourth collection date followed by 1 tree being attacked during each of the last two weeks of the experiment, giving 3 of 8 replicates attacked. Mean numbers of *X. glabratus* per types of traps, e.g., each host tree, fake tree trap with and without lures were as follows: host tree 28.9 ± 4.6 (sum 1,288), fake tree with lure 274.2 ± 26.8 (sum 13,161) and fake tree without lure 55.9 ± 10.5 (sum 2,682). A linear regression (Proc Reg, SAS 9.4) of number of beetles collected relative to size of the fake trees and the size of the host trees at DBH was not significant: F value 1.1, Pr. > F 0.30). *X. crassiusculus* appeared on traps at very low numbers (less than 10/trap/count) until the fourth count when the redbay tree was attacked at which time the number of *X. crassiusculus* captured rose into the 300-500 range on both the traps on the tree and the fake tree with the lure. This did not happen at the fifth and sixth collection time when the second and fourth trees were attacked by *X. glabratus*.

**2.4.3:** To further evaluate the response to fake and natural trees for attraction of beetles to hosts and non-host trees, 2 fake UV mulch trees were provided as one on each side of the natural trees (n=6 total) at 1, 2 or 3m, one with the lure the other without a lure, same as in experiment 2 above. Three redbay, 2 beech and a wild cherry tree were used to provide 1 host and non-host combination for *X. glabratus* at each distance. The fake UV mulch trees used were placed over a bamboo pole vertically and all were 10cm wide by 1.22m long with a 15 by 30cm trap covered with Tangle trap in the 61-73cm space above ground and around them. The lure used was α-copaene for *X. glabratus* which was rotated from one to the other fake trees after each count. Counts were made once a week over 9 weeks. The data were analyzed using Proc ANOVA and Tukey’s Studentized Range test (SAS 9.4).

**2.4.3 Results:** At week eight one redbay tree with fake trees at 2m was attacked by *X. glabratus*, otherwise the experiment was completed without attacks of the redbay centered trees. Fake trees and both the nonhost and host trees (via their Tangle traps) attracted *X. glabratus* in high numbers (700+/week) but only a relative few of *X. crassiusculus*. Distance between trees and numbers of beetles collected manifested by the placement of the fake trees was not statistically different (F = 0.53, P > 0.59) and (mean ± SEM/trap/week): 1m, 4.8 ± 4.2, 2m, 5.3 ± 2.5 and 3m, 5.2. ± 4.0. Lure placement was significantly different (F = 128.7, P > 0.0001), but interactions between distance and lure placement were not (F = 1.03, P > 0.35). A Tukey’s Standard Range test indicated that the lure placement and collection of a mean/trap/week of 9.38 (A), > than host tree 3.99 (B) and > than fake tree without a lure of 1.96 (C). Fake trees, when without lures, attracted mean numbers of *X. glabratus*/week anywhere from a minimum of 22 in week 2 to 170 in week 3. The overall mean ± SEM was 24 ± 9.3 percent of the total numbers of *X. glabratus* caught/week on any fake tree or natural tree traps. This result indicates the strong visual attraction of UV mulch to this invasive ambrosia beetle. Moreover, the trap numbers on non-hosts in this experiment were similar to numbers on the main host redbay, again indicating the unexpected attraction of ambrosia beetles to UV mulch even from small size traps.

#### 2.5: The importance of host tree physical factors on ambrosia beetle attraction behavior

The importance of host plant characteristics on beetle behavior was important to further determine if a “fake tree” trap (at the time thinking about a tube shape) had potential for further development. The previous experiments were followed by determining *X. glabratus* response to UV mulch-treated trees of different sizes and at different heights. Small (<20cm) and large (>20cm DBH) non-host trees, n=6, 3 of each size, wrapped in UV mulch from 0-2m in height were deployed. Each of the six trees carried only 1 of the size/height treatments. A 15cm piece of UV mulch was also wrapped around the trees to capture the responding beetles. The UV mulch strips were placed around the trees to fit over the middle of the heights tested (1 each/test tree), e.g., centered over 30, 60 and 90cm above ground. A lure was added to each section (only 1 each) on the trees. The number of beetles present on the wrapped trap were counted after 1 week in the field on one half of the size of the trap to make the counts as presented. The experiment was conducted twice two months apart in north Florida using 6 trees in the first run and 12 trees in the second run (Fig. 9 and 10, respectively).

**Figure 9:**
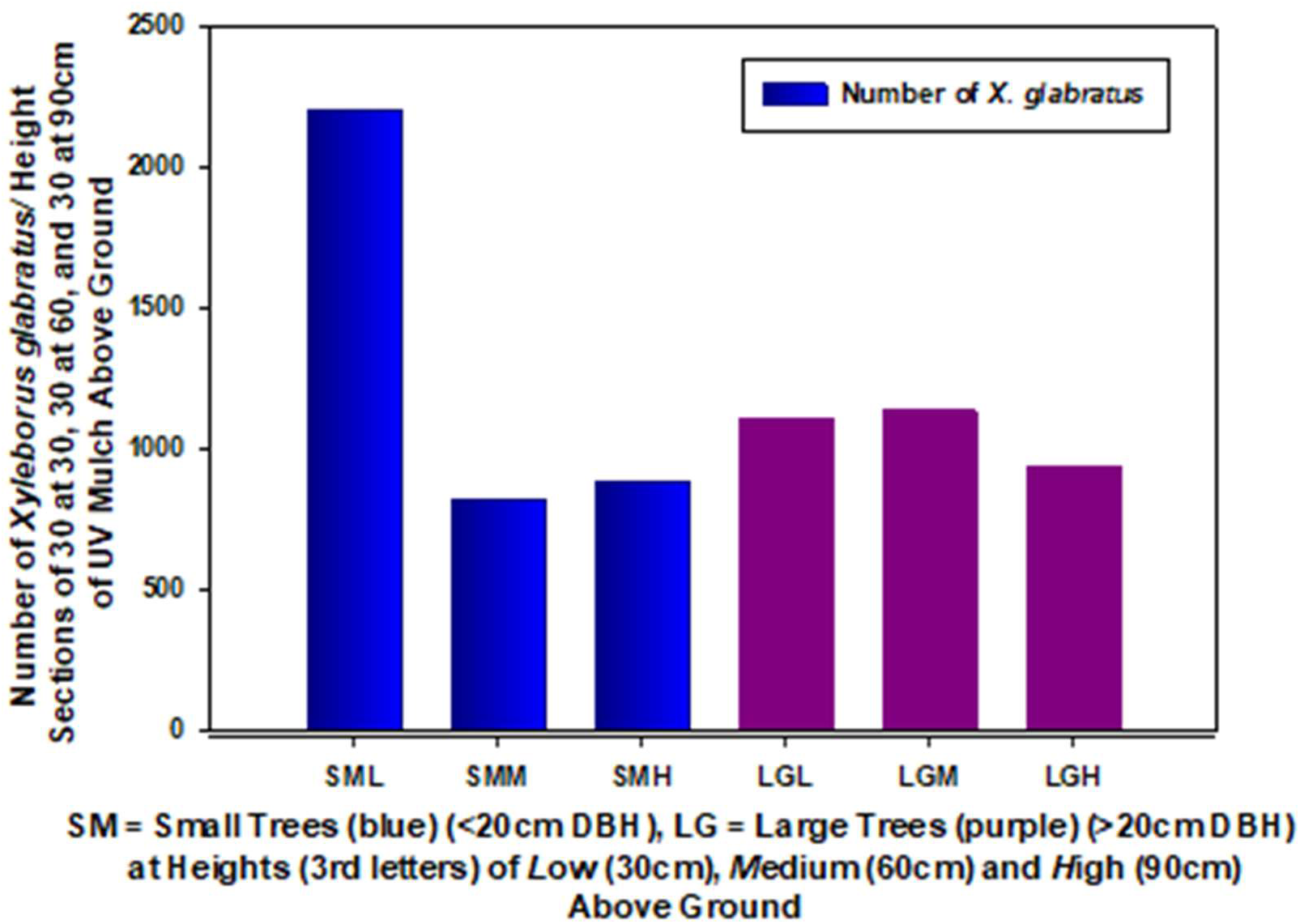
Number of *Xyleborus glabratus* captured over a 1 week period in small (<20cm) and large (>20cm DBH) non-host trees, n=6, 3 of each size, wrapped in UV mulch from 0-2m in height. The colors have to do with the size of the trees left to right. A 15cm piece of UV mulch was also wrapped around the trees to capture the responding beetles. The UV mulch strips were placed around the trees to fit over the middle of the heights tested (1 each/test tree), e.g., centered over 30, 60 and 90cm above ground. The number of beetles present on the wrapped trap were counted after 1 week in the field on one half of the size of the trap to make the counts as presented.

**Figure 10:**
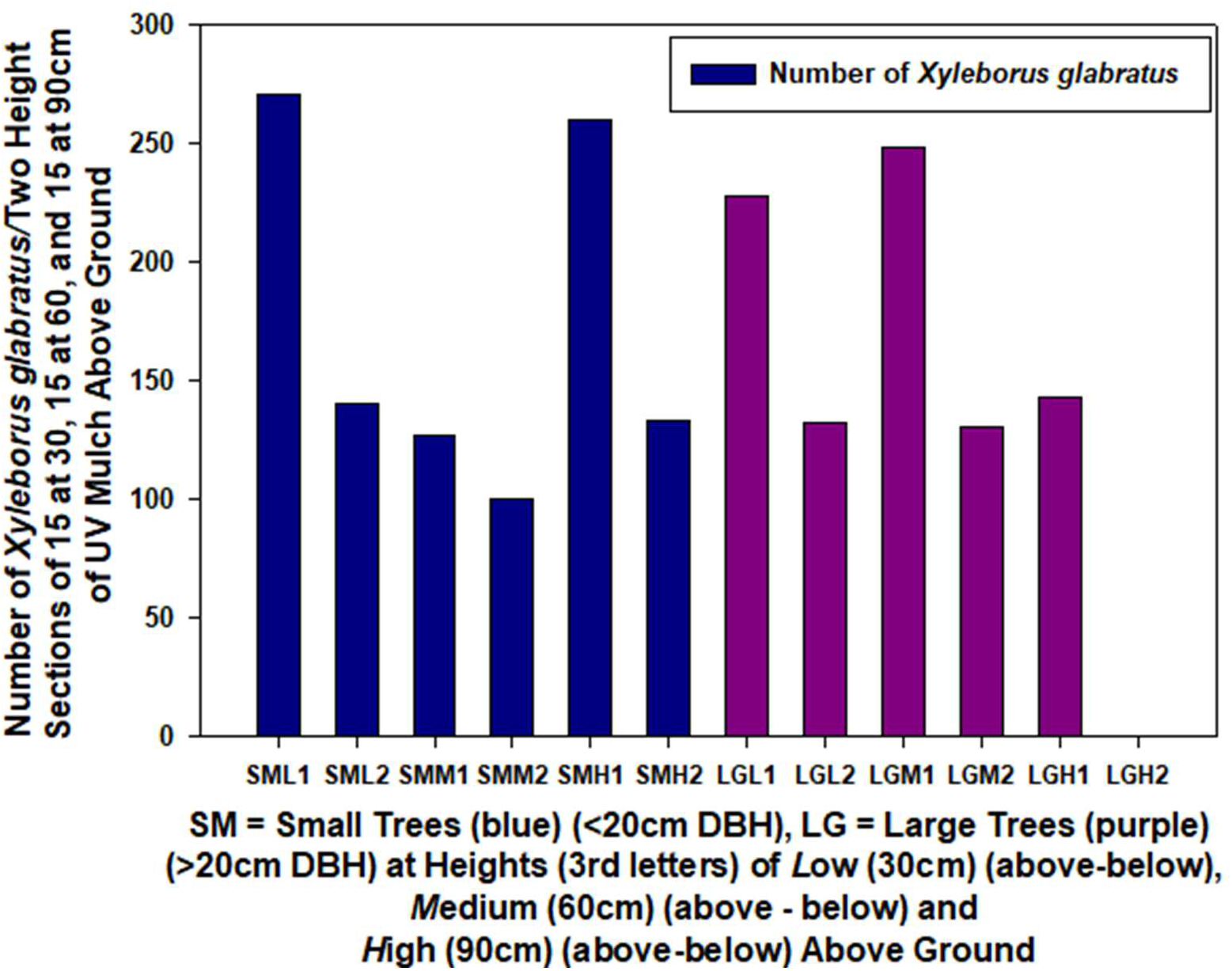
Number of *Xyleborus glabratus* captured over a week period in small (<20cm) and large (>20cm DBH) non-host trees, n=6, 3 of each size, wrapped in UV mulch from 0-2m in height. The colors have to do with the size of the trees left to right. A 15cm piece of UV mulch was also wrapped around the trees to capture the responding beetles. The UV mulch strips were placed around the trees to fit over the middle of the heights tested (1 each/test tree), e.g., centered over 30, 60 and 90cm above ground. The number of beetles present on the wrapped traps were counted after 1 week in the field on the entire size of the trap by 1 half each to make these divided counts as presented. The zero count in the LGH2 in the righthand side of the figure did not capture any beetles and was the only section without beetle response.

**2.5 Results:** The numbers of *X. glabratus* captured over a 1-week period by treatments are presented (first time Fig. 9 and second time Fig. 10). The results indicate that there was some response to both height and size of the trees but not an overwhelming behavior such that it would interfere with attempts to develop a new trap. Moreover, the more important factor was that lots of *X. glabratus* were present and relatively large numbers of beetles were present on the UV mulch-covered trees on all sizes of trees at all heights. This coated tree experiment with small and large size, along with height backs up the line of experiments to get to a full fake trap from tubes. It was used to make sure it worked to show there was *X. glabratus* response to all “trunk” sizes and heights and then likely tube size- which it did.

#### 2.6 Development and evaluation of a “fake tree” trap using UV mulch as the attractant

Due to the discovery of the surprising attraction of ambrosia beetles to UV mulch (UVM) and the visual equivalent as indicated above, bright metallic silver paint, efforts were initiated to further test UVM on a new trap as a monitoring and suppression tool, etc. The first idea was to develop a “fake tree” to use as an enhanced visual tool to accompany and deliver the UVM as a better attractant for ambrosia beetles. The logic here was the apparent tendency of *X. glabratus* to land on small and larger trees as shown above. The first experiment was aimed at further determination and optimization of the fake tree size (diameter) and shape (round or square) for several reasons: relative costs, efficacy, and potential usefulness as a scientific tool. Mailing tubes 1.22m long and 5cm or 10cm wide were obtained (Uline, Inc.) along with building form tubes (footers, trusses) (QUIK-TUBE, The Home Depot, Inc.) of 1.22m long and 20cm and 25cm wide (see supplemental material). Tubes were covered with UVM, a second piece of UVM of 15cm width was wrapped around each tube at 61cm from the end and covered lightly with Tangle trap. A wire was attached at the top of each tube for hanging from a tree or another wire. Five tubes of each size as described were placed in the forest in groups of 4 (=blocks, 1 trap of each size) and spaced 61cm apart while hanging upright from tree limbs or wire that was placed across trees to accommodate the hanging need so that the traps remained vertical. Each set of 4 fake tree “trunks” (now “tube traps”) were randomized in placement and a lure of α-copaene was placed near the top of the wire holder or tree limb supporting the tubes. Traps were checked for ambrosia beetles (primarily *X. glabratus*) once a week over seven weeks. When beetles were counted and removed, Tangle trap was cleaned and replenished if necessary, and the 4 tubes/block were rerandomized relative to their positions next to each other.

**2.6 Results:** Mean capture rates on the tubes of 4 different sizes varied per trap over 7 weeks from 160-240 and indicated that the *X. glabratus* counts on the smallest tube trap was significantly lower than the 3 larger tubes (Fig. 11). Despite a large numerical difference between the 10cm and 20 and 25cm sized tube traps ∼25% (160 vs 200) it was not statistically different using the Least Squares means and the populations present of *X. glabratus* during the experiments. However, Fig. 12 shows that for one week where the number of *X glabratus* captured was highest by a factor of 10, there was a very strong response (slope of >1 in the regression in Fig. 12) to the trap sizes. These results provide evidence for a conclusion that *X glabratus* preferred the traps with the larger diameters (20 - 25cm) and that the 20cm (8”) trap that reached the asymptote was as effective as the 25cm (10”) diameter (Fig. 11 and 12).

**Figure 11:**
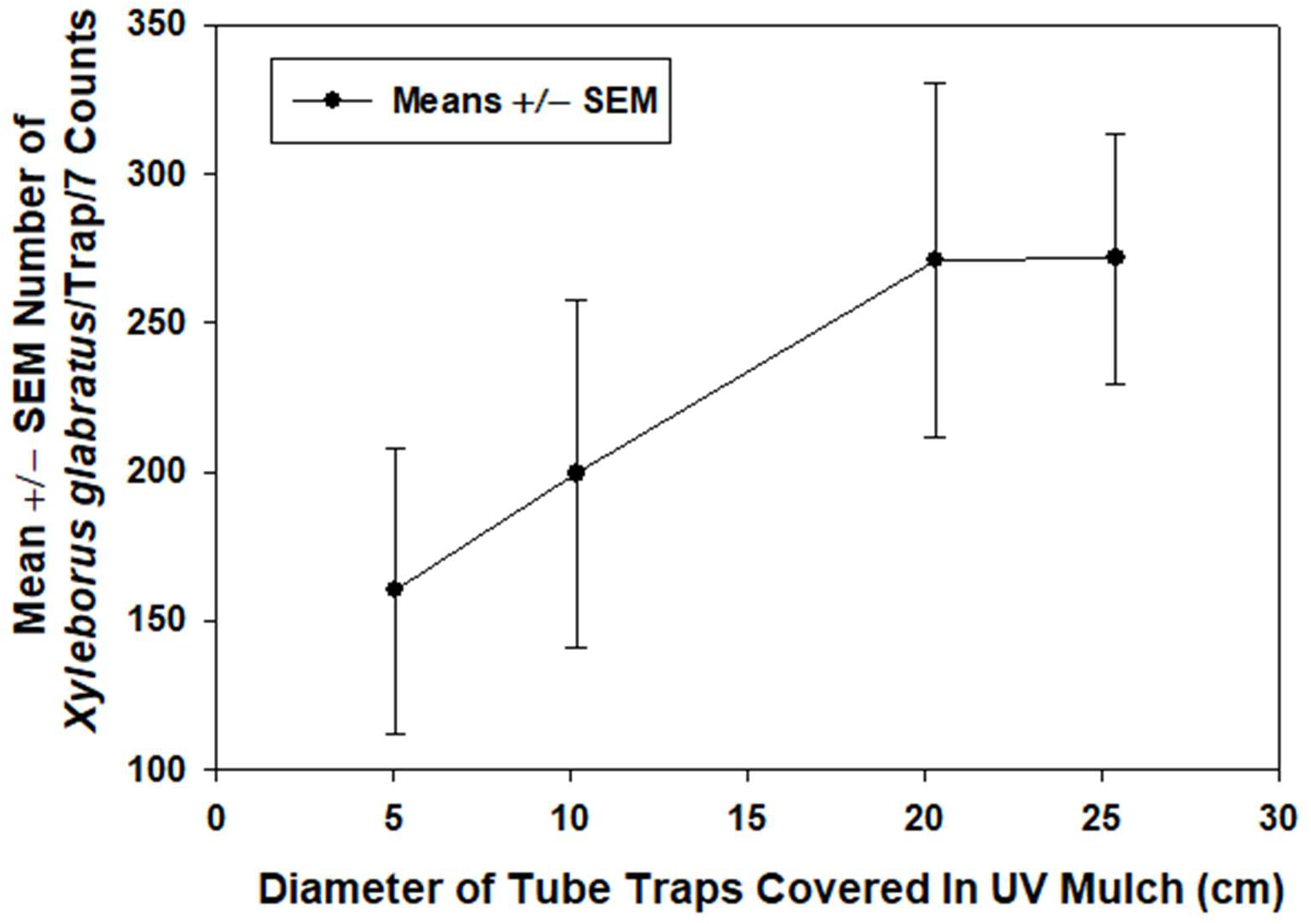
Mean ± SEM number of *Xyleborus glabratus* per diameter of cylinder-shaped UV mulch traps 1.22m in height.

**Figure 12:**
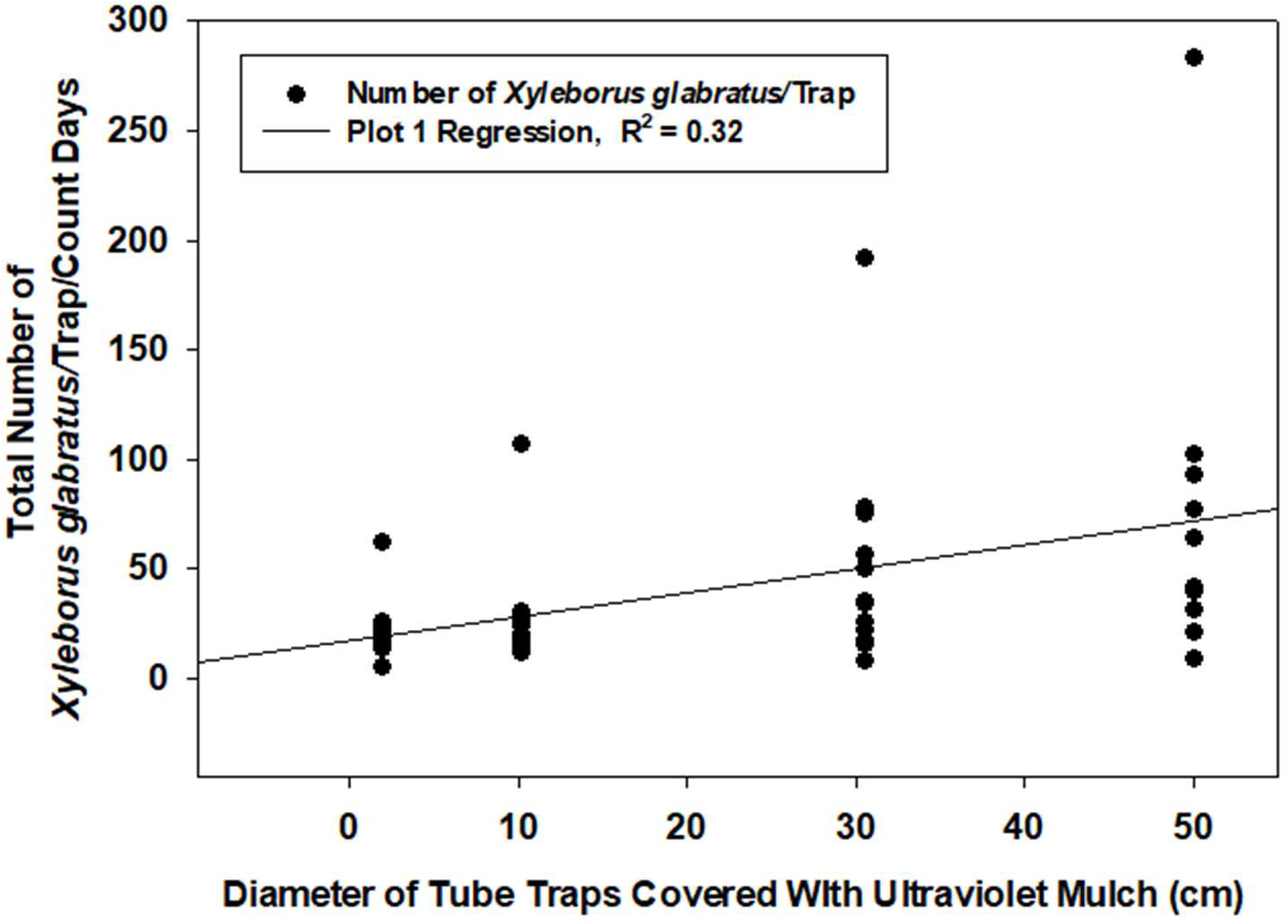
Total number of *Xyleborus glabratus* per trap size in relation to 9 weekly counts. Note that the more beetles that were collected the more linear the results that provided a better manifestation of the beetle’s preference for larger traps as with trees.

#### 2.7 Further optimizing the fake tree trap and tool using UV mulch as the visual attractant

The results above, using the Home Depot Quik Tube, indicated that a tube 20cm in diameter was as attractive as a 25cm tube. These tubes were considered expensive, at the time costing between $10-15.00 each. It was reasoned that cheaper traps could likely be substituted, and the following describes the methods used to achieve a cheaper, better and more easily used trap, e.g., lighter in weight and for enabling broader scientific uses. Toward that end, other types and sources of tubes and boxes were obtained, processed and tested in a number of ways. Heavy cardboard boxes that were lighter and square (∼20cm) in shape were obtained. The original 20cm tube traps described above were cut in half to make 20 cm (width) x 60cm (length) traps. Cardboard boxes “200lb, Test Long Corrugated Box” (Uline, Pleasant Prairie, WI 53158) of 91 x 21 x 21cm (Uline # S-18928 at $4.26/each in box of 15) and 91 x 21 x 21cm (Uline S-4361, $2.25 each per box of 25) were purchased. The Uline boxes were ∼square in shape unlike the round Quik Tube as above. The traps were mounted on a 1.22m sharpened, wooden tomato stake with 20cm of supporting length inside the trap and 132cm of length below the trap for placing vertically in the ground at ∼ 15cm depth. All 3 trap types were covered with UVM material using 3M Super 77 spray glue. Special care was used with the UVM material to overlap the sides and tops of the traps to make them fully waterproof. An additional 15 x 30cm rectangle of UVM was stapled on one side in the middle (height) of each trap and covered with Tangle trap. Each trap received a lure of ethanol (AgBio, Inc.) placed ∼10cm below the top of the trap and above the extra sampling UVM piece with Tangle-Trap™. The 4 types of traps (∼20cm in diameter and 61 or 122cm in length) were then placed in the forest at 30m intervals within a grid and randomly arranged in 4 rows of 4. Traps were checked 1-2 times per week for ambrosia beetles on the attached Tangle-Trap™- coated part of the trap. Tangle trap was cleaned and changed as needed.

**2.7 Results:** There was no significant difference between the total number of ambrosia beetles (mostly *X. crassiusculus*) caught in any of the trap types (data not shown). This result enables selection of the cheaper, money-saving, square box traps and a shorter version of the two at 61cm (2’) long if of interest. The experiment was repeated in another location and along with the above traps 122 x 20cm “QUIK-tube” traps were added with and without a 61cm plastic lid to reduce rainwater and heat from sunlight affecting the traps. As in the above experiments, no significant differences were observed between any of the tested traps in terms of total ambrosia beetles captured/trap (data not shown).

#### 2.8 Trap sizes

The change over from round “QUIK-tube” cardboard traps to 61cm and/or 91cm length cardboard boxes in 20cm (∼8.5”) squares wrapped in UV mulch found that all were similar, no significant difference in numbers of *X. crassiusculus* trapped when ethanol lures were used. However, surprisingly, numerically more *X. crassiusculus* of ∼ 45% were caught in the 61cm shorter traps (2’) vs the 91cm (3’) traps. As a result of the wide variation often observed between traps, an experiment was formulated to determine if the 20 x 20cm wide box trap could be changed in some manner to affect beetle visual response and possibly change the trap to trap variation physically. Pieces of cardboard 25 x 40cm were cut out and covered with UVM as described above. These pieces, “EARS” so to speak, were added to a trap for comparison by stapling them to a trap vertically focused on the mid portion of the trap’s height. The usual separate piece of UVM material 15 x 20cm covered with Tangle trap. was added to a trap side without an EAR. The treatments tested were a control without an EAR, a trap with one EAR and a trap with 2 EARs - attached on opposite sides. An ethanol lure was added on the top of each trap and 4 traps/treatment each attached to a wooden tomato stake as described above were tested in a randomized complete block design. Adding the EARs to the traps in effect increased the size of the traps and therefore the visual cue size in 1 or 2 of the cardinal directions by single or double the size of the normal width of 20cm.

**2.8 Results:** Adding 1 or 2 EARs to the box-shaped traps did not affect the counts of responding ambrosia beetles (mostly *X. crassiusculus)* to any treatment in any manner (data not shown). These results indicate that behavioral response to host trees using vision and odors has a hierarchy for ambrosia beetles whereby odors are more important than vision at close range. Later experiments discussed below will add further explanation of this observation. Moreover, the experiments conducted had no control over the population of beetles that were present at any time.

#### 2.9.1: Use of a 4-sided (square) cardboard structure of the UVM fake tree trap to compare treatments by exploiting the cardinal directions of the 4 sides

Assume a UVM-covered QUIK-tube trap and a square UVM trap (as above) are oriented so that the sides of the traps are aligned as direct as possible with the accepted 4 cardinal directions. A physical and visual comparison of a round QUIK-tube and a 20cm square box trap relative to the visual target, the relative potential accompanying attraction level, and behavioral action by ambrosia beetles to it, will find that the round-shaped QUIK-tube, even though it is more or less the same diameter as the square box (20cm), will present a wider but continuously changing visual target to the beholder as it/they move even slightly left or right in cardinal direction space as the trap is approached. This is due to the QUIK-tube’s smooth circular sides without corners like a square trap. The square UVM trap, on the other hand, due to its well identified shape with 4 flat and right-angle corners, will present a more constant, well defined, and narrower visual target to an approaching beetle (or any other entity approaching the traps) maintaining its pathway within the cardinal direction lines, e.g., the pathway from which it starts response. Thus, the name was changed to UVMT and the view of the UVM square trap is constant and smaller in size due to the flat shape and longer sideways movement needed to see the adjacent sides of the trap in either direction when moving toward the trap. As a result, the two types of traps offer different potential uses to better understand the behavior used by the target organism(s). Thus, the UVMT square trap offers the ability to carry 4 different treatments at once, e.g., within each of the cardinal directions and constant angles, without visual overlap at close distances from small sideway movements by the responding organism. This factor will obviously relate to the orientation pathway angle(s) changed related to the trap by the approaching organisms and distance away from the trap of the responder; at a longer distance away, one will see more of the square sides, and this will also happen with movement angles closer aligned to the angles of the trap edges. These results also relate to why and how ambrosia and bark beetles (see below) respond visually to UV mulch when in fact UV mulch is used under many situations to repel leaf-feeding insect-pest species such as whiteflies, thrips, etc.

#### 2.9.2: Use of the UVM square trap to compare and contrast the visual behavior responses of ambrosia beetles to various colors, compounds, light spectra and material patterns with their implications and potential uses

All UVM square traps (UVMT) used in the following experiments were 91cm (3’) long, ∼20cm (8”) wide (on all 4 sides), attached as above to a wooden tomato stake, and had an ethanol lure (AgBio, Inc.) attached ∼10cm below the top of the trap and relative to the treatment side(s) as indicated for each experiment. Counts, when provided, consisted of 3 factors: number of *X. crassiusculus*, number of other unknown ambrosia beetles, and total ambrosia beetles per treatment replication (= the first 2 summed) if they differ significantly in numbers trapped.

#### 2.9.3: Sunscreen and other light control chemicals as new tools to improve understanding of ambrosia beetle visual behavior under outdoor conditions

Two kinds of sunscreen agents are recognized whose composition and mechanisms of actions differ. They exert their effects on sunlight by screening through blocking, reflecting, absorbing and/or scattering sunlight (Latha et al. 2013). Organic types, e.g., with the common ingredients of escamsule, silatriazole, bemotricinol and bisoctrizole, are broad-spectrum sunscreens that absorb UV rays, whereas inorganic, physical blockers, with “metallic” compounds, e.g., zinc oxide, titanium dioxide and related, reflect or scatter light. The third related component, commercially available as “UV Blocker”, “UV Killer”, and “UV Shield” (Atsok, Inc. Orangeberg, SC), etc., are used because they block or enhance the UV wavelengths. They are often used by deer hunters to avoid the “metallic’ cleaning materials that are contained in the clothes being detected by Cervidae (deer) that are color blind.

#### 2.9.4: To determine the impact on ambrosia beetles, mainly the species most available, *X. crassiusculus*, behavior in response to various mentioned light regimes created by different light-effecting materials as above

Because the UVMT has 4 different sides, it provides a unique and excellent scientific tool to study insect behavior and to add and compare the impact of the 3 different wavelength-affecting options described above on ambrosia beetle behavior in response to the natural attractive UV mulch. Thus, experiments were set up to test these ideas concerning beetle response to UVMT treated with various sunscreens. The exact sources used were as follows: 1. ultraviolet blocking (UV-Shield by ATSKO, Inc., Orangeburg, SC, 29115,) a material used on clothes by hunters (referred here as UV blocker), 2. a light-reflecting mineral-based product with 4.3% titanium dioxide and 15% non-nano zinc oxide) by Bare Republic, (Coolla, LLC, San Diego, CA 92010), and 3. a multiple chemical-based, light absorbent, Sport Sunscreen SPF 50, Equate™, containing 3% avobenzone, 10% homosalate, 5% octasalate and 5% octocrylene (WalMart Stores, Inc., Bentonville, AK, 72716). These three sunscreen types were set up using a regular spray or paint brush with UV-Shield - the true UV blocking agent on 1 per side of the UVMT along with a standard side as the UVMT control. Each of the 4 sides were covered with a 10 x 30cm piece of UVM as described above covered lightly with Tangle trap to capture the beetles on each of the 4 available treatment sides. Lures of ethanol were added at the top middle of the trap so as not to bias the odor attraction to any of the 4 sides. Five traps were covered with these 4 randomized treatments/trap for each experiment. Beetles were counted every 2-3 days over 31 days to gather the results. The effect of the reflecting metallic material from Bare Republic did not last as long in the field as the other treatments. Also, in another previous experiment with the coroplast rectangles discussed above, UV-Shield, the UV Blocker, was also tested and the beetle numbers (*X. glabratus* and *X. crassiusculus*) collected did not differ from the controls (data not shown). This earlier result suggested that ambrosia beetles are not visually nor behaviorally affected by ultraviolet wavelengths of light (assuming that is what effect the chemical has). Moreover, previous experiments by Horvath et al. (1998) have shown that aluminum foil is very similar to UVM in terms of reflected light including the lack of any reflectance of polarized light that is responded to in several ways by Tabanidae and other aquatic-associated insects. This assumption was made here due to the results, the observed strength of the published information, and the lack of access to equipment for polarity measurement.

#### 2.10 Traps and trapping

Five cardboard-based, 91 x 21.6cm, square-shaped traps (S-18928, 36 x 8 x 8” 275# DW Box 15/180, Uline, Inc., PO Bx 88741, Chicago, IL 60680), were used for the experiment. Once the cardboard was secured with duct tape and staples, they were attached with wood screws to a 1.22m tomato stake placed into the inside ∼51cm. The cardboard traps were then covered completely with UVM as described above. On each of the 4 sides of the traps a 20 x 30cm piece of UVM was added within 10cm of the trap top by stapling. This served as the trapping surface for the responding ambrosia beetles. Each of the 4 sides were used as treatment surfaces. The treatments were added and then covered with Tangle trap. An ethanol lure was added to the side with the UVMT treatment which served as the control treatment. The patterns of the 4 treatment placements were randomized as much as possible and the traps were set up so that the control side (UVM + lure) pointed north. After each count each trap was rotated clockwise by 1 quarter turn. The 5 UVMT traps provided 5 replicated comparisons of the 4 treatments as follows and identified above: 1. a control surface of UVMT only plus the ethanol lure, 2. a UVMT surface covered with the mineral reflecting material Bare Republic Vanilla Coco, 3. a UVMT surface covered with a UV blocker (UV Shield), and 4. a UVMT surface covered with an absorbent material, Sport Sunscreen as above. The 10 x 25cm UVM pieces were covered with the sunscreen materials first then Tangle-Trap™ was lightly applied over the treatments on each trap. The three sunscreen materials were reapplied to the traps on the tenth day but due to weather had no noticeable effect. Once completed the traps were placed via the tomato stake into the ground by drilling a requisite hole 15-20cm deep. Thus, the bottom of the trap was ∼50cm above ground with the treatment portions within the 70-122cm space above ground on the traps. The finished traps were placed on the edges of wooded areas with habitats determined in previous above experiments that were sparsely filled with hardwood trees and other vegetation, e.g., typical habitats for ambrosia beetle movement pathways. The traps were baited with ethanol lures stapled to the top side of the UVM control side on each trap. Note: the lure placement varies in these experiments, observe accordingly! Traps were checked and counted every 2-3 days over 13 days in March-April, 2020 in Monticello, FL. *Xylosandrus crassiusculus* is usually the most prominent ambrosia beetle species found in North Florida during that time and are easily recognized in the field. Note that *X. glabratus* had not been found in the area of this experiment at the time it was conducted. The beetles were counted and recorded for each trap by treatment and removed along with any other observed “unknown” ambrosia beetles. Unknown beetles were counted and recorded. Traps that had leaves, other insects, or items caught in the Tangle trap, were cleaned, e.g., removal of the object each check day. The traps were then turned 1 quarter turn clockwise to ensure the trap sides (= treatments) got equal ambient light exposure from each of the cardinal directions.

**2.10 Results:** Because the UVM side of the 5 traps are the sides with the strongest odor cue, the expectation was that this experiment, to show differences between the visual treatments, would mean that the treatments are strong competitors with the standard lure-containing side. The results show the response by *X. crassiusculus* to the treatments over 13 days as expected and displayed (Fig. 13 and 14) equal response to the control and UV blocker treatments, higher numbers in response to the reflected light on some days – particularly day 1. After that day the reflecting material did not last long outdoors (especially in the first sunscreen experiment Fig. 13 above) and was lower to the absorbed light treatment. These results vary early after application by ∼12% lowest to highest day 1, are equal due to a much cooler temperature that occurred on day 2, returned back to a similarity to day 1, but with the change to lower numbers from the reflectance treatment (see Fig. 14, now with the same Fig. 13 shown without the reflectance treatment for clarity) then different within 5-8 % before being exactly equal on the last 3 cooler nights. The UVMT with lure treatment and both visual and odor factors was usually very close to the UV blocking treatment indicating that the lure and UVMT attract the beetles while ultra-violet wave links are not responded to by ambrosia beetles. Moreover, the UVMT evoked very strong visual response compared to the odor, whereas the sunscreen absorbent affected (lowered response numbers) the entire light spectrum by reduction in the *X. crassiusculus* response (Fig. 13 and 14). The overall perspective with differences in least square means shows significant difference between the absorbing treatment and both the UV mulch control (6.64, DF=16, t value 2.93 P > t = 0.10) and the UV Blocker (4.74, DF 16, t value 2.09, P > t = 0.05). Along with the captured *X. crassiusculus*, a modest number of unknown species were also captured. Adding that data indicates the control treatment is significantly higher than all three of the other treatments by P > 0.0029 absorb, P > 0.029 reflection, and P > 0.167 for UV Blocker (only numerically). Least squares means indicated that over all days the mean numbers of *X. crassiusculus* captured by the treatments were: control 14.4 ± 1.6, UV Blocker 12.4 ± 1.6, reflection 11.2 ± 1.6 and absorption 7.7 ± 1.6. The control and UV Blocker were significantly higher than the absorption treatment at Pr > t = 0.01 and Pr > t = 0.53. Along with the *X. crassiusculus* there were a number of unknown species caught in the traps at the same time. Those results combined with *X. crassiusculus* numbers (means/treatment) were as follows: control – 44.3, reflections 51.1, UV blocker 40 and absorption 28.4. The control was significantly greater than the other three treatments with Pr > t > 0.03.

**Figure 13:**
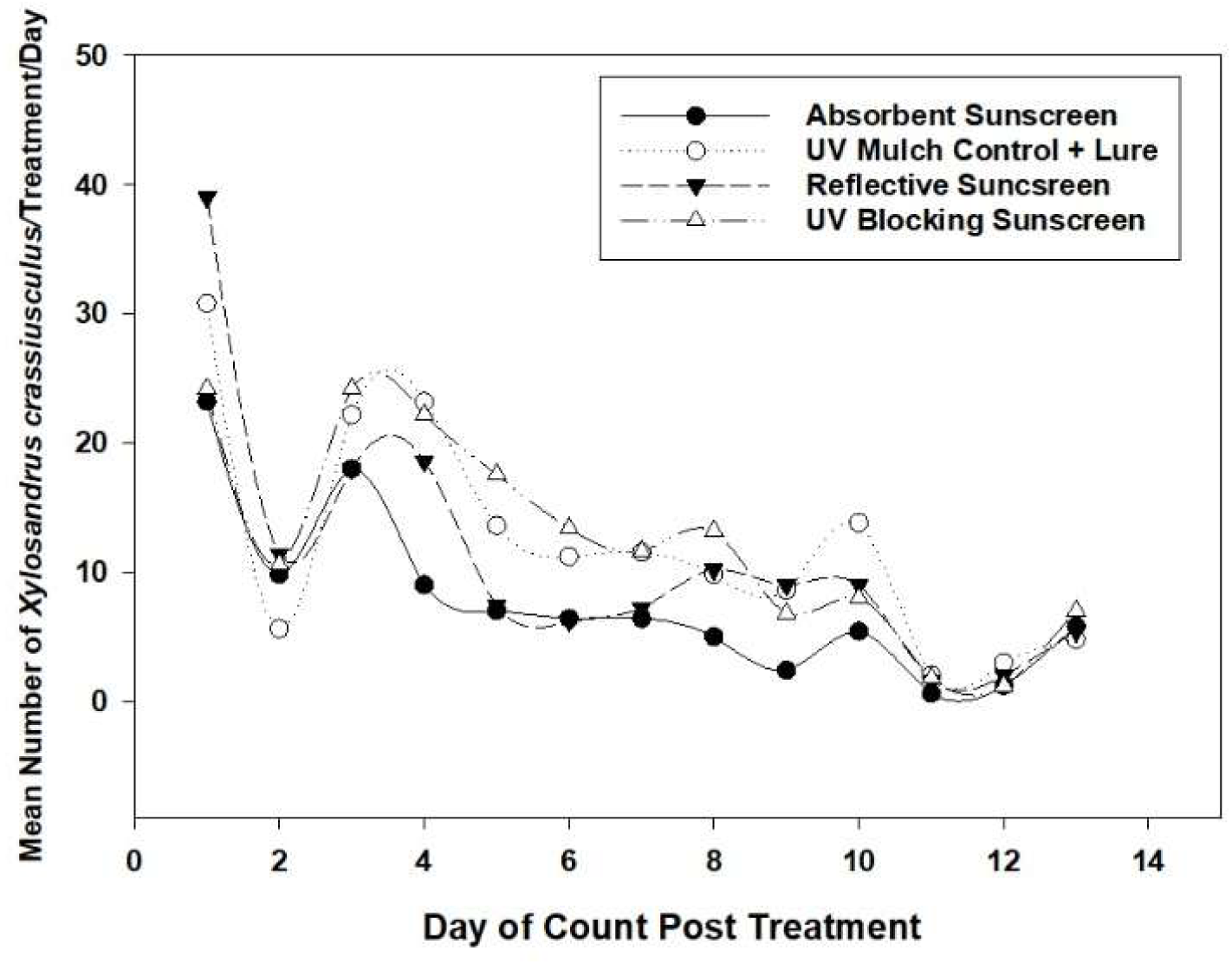
Response of *X. crassiusculus* to a square UV mulch trap containing a UV mulch side with the ethanol lure and the other 3 sides each with a sunscreen treatment - absorbent, reflectent, and a UV blocker.

**Figure 14:**
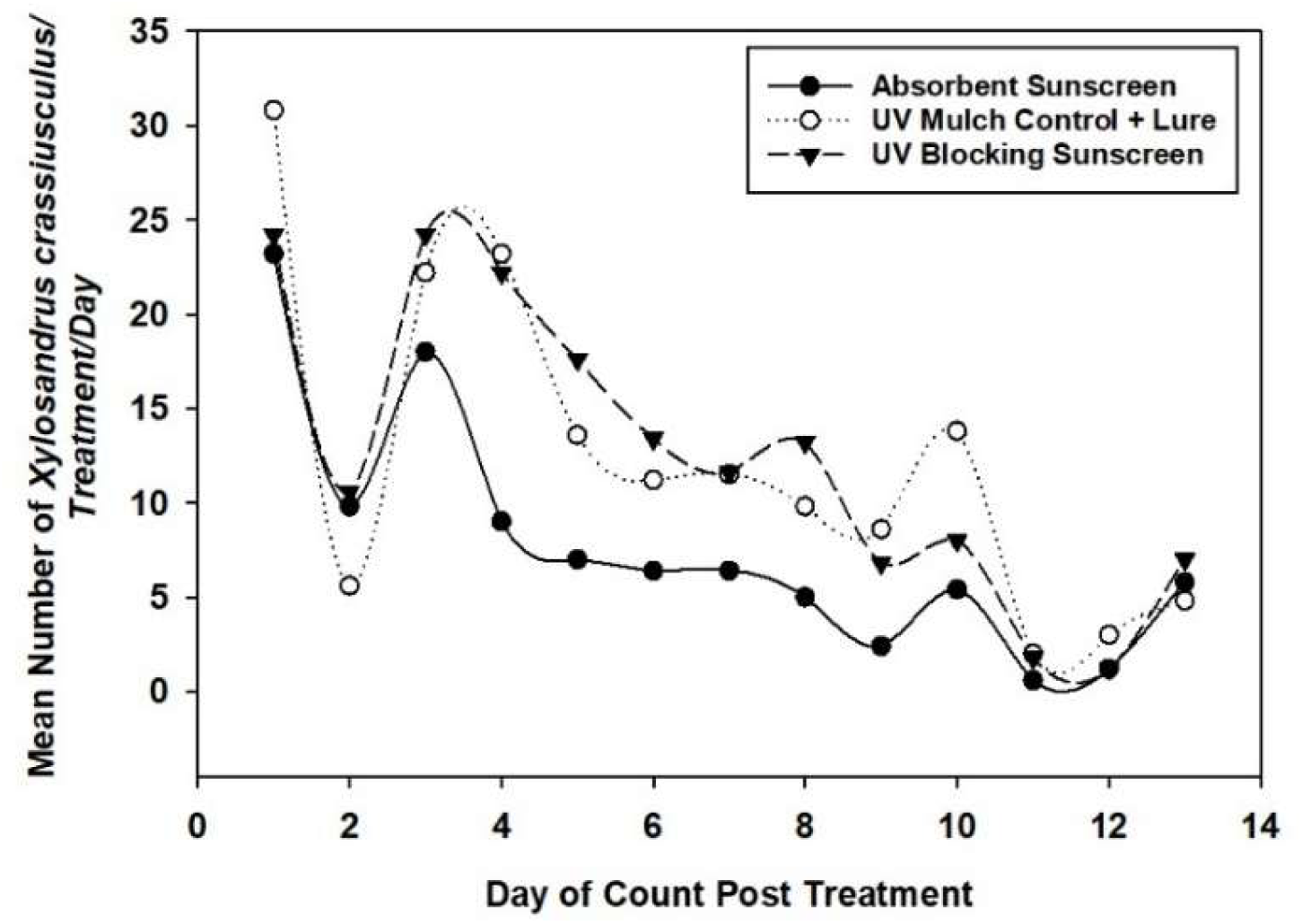
Response of *X. crassiusculus* to a square UV mulch trap containing 3 single side sunscreen treatments: the absorbent, the UV blocker (without the reflectent which was removed for visual clarity) vs a standard control, the UV mulch side with the ethanol lure.

#### 2.11: Adding solar lawn night lights during dark hours also had no positive or negative effect on UVMT trap numbers

Given the brightness of the UV light ostensibly reflected by UV mulch, one might guess that adding increased light to the UVMT might increase beetle response given that they fly in late afternoon and evening as the sun goes down. Thus, using the UVMT as the trap, a common solar-powered LED light in a low (20 Lumens) and a higher power (60 Lumens) (Mainstays™, WalMart, Inc., Bentonville, AR, 72716) were added to shine (pointed at) on one side (a cardinal direction) of each trap. The lights were naturally triggered by the lack of light in the evening part of the day. The lights (1 per trap) were placed as normally used in the soil around the traps at random cardinal directions. From their original randomly (n=4 choices) selected cardinal direction, the lights were rotated clockwise by 1 direction after each count. Most ambrosia beetles fly from mid-afternoon to dark so the lights would have increased the entire light spectrum available for the beetles’ response mainly later in the day and overnight.

**2:11 Results:** Unlike the positive results from green LED lights reported earlier (Gorzlancyk et al. 2014) with *X. crassiusculus*, *Cnestus mutilatus* and other ambrosia beetles, no increases in response numbers were observed regardless of the light’s power, actual time of day or in what cardinal direction that the lights were placed (Fig. 15). This suggests that the UVMT is naturally attractive enough to gather and reflect enough natural sunlight that is present during ambrosia beetle late afternoon-evening flights to optimize capture rates.

**Figure 15:**
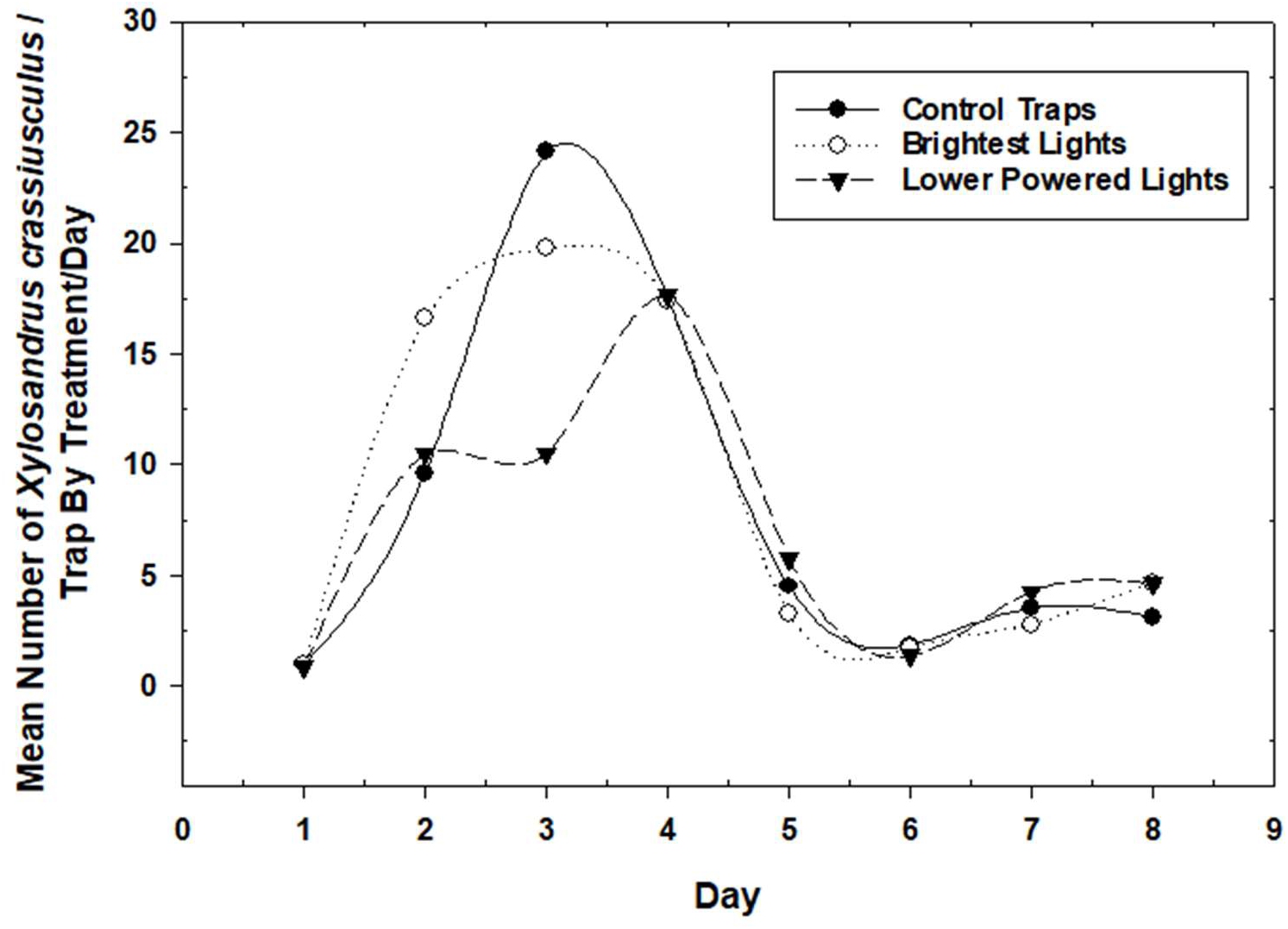
Response of *X. crassiusculus* to UV mulch-covered, square traps with portable solar-powered, outdoor, house lights in one cardinal direction each. Each trap and light were rotated clockwise 1 direction after each count.

#### 2.12: Effects of specific colors on ambrosia beetles: response to yellow when on the UVMT

Yellow hues were chosen for this experiment for three reasons: 1. Most plant feeding insects respond to and/or often prefer yellow when provided a choice of trap colors, 2. Yellow is a color associated with plants and trees specifically and mostly considered as an attractant but has not been found as so for ambrosia or bark beetles (R. Mizell, above and unpublished experiments), and 3. The forest habitat of these woodboring beetles is overwhelmed with the colors of yellow and green which are close in wavelength on the light spectrum. Because of the obvious contradictions between the response to UVM and colors, and the accepted knowledge of beetles limited response to colors, the potential behavior displayed by ambrosia beetles and how it might be exploited for potential strategies and tactics in insect pest suppression was investigated further using the new UVMT. Six box-shaped UVMTs (trap name used from now on) were set up as above and 3 hues of yellow were developed using coroplast material in 2 ways. Coroplast (2mm thickness) was purchased (Corrugated Plastics, 601 Rt. 206, Suite #26-476, Hillsborough, NJ, 08844) in a bright but darker yellow closely resembling “safety yellow” (Rust-oleum #242258 safety yellow). Two other spray paints were selected such that when compared to the natural yellow coroplast there was one in a “middle” appearing yellow hue and one in a lighter, more “whitish appearing” yellow hue relative to safety yellow. The coroplast was cut into pieces that fit the sides (∼22 x 122cm) of the UVMTs and maintained as the original brightest hue or painted one of the other two yellow hues. A UVMT trap was then set up with one of the 4 sides retaining (covered by) the basic UVM and the 3 other sides were covered with one of each of the yellow coroplast-painted sheets using staples. The juxtaposition of the 4 colors was randomly placed in the side locations on the traps to maximize the number of relative spatial side by side relationships. To maintain this pattern relative to potential light directional effects and to remove any potential position/locational bias, all traps were placed in the beginning of the experiment oriented in the forest with the UV mulch side facing north and then rotated 1 cardinal direction in a clockwise rotation after each count. Six traps were used and counts were made on each trap by treatments every other day over 5 days. For this experiment, the entire areas of the treatment sides were covered with Tangle trap. Beetles were removed from the traps, and the Tangle trap was cleaned and replaced where necessary. An ethanol lure was added on the top of each trap to reduce any bias difference in odor attraction to any single individual side of the trap as above. Counts were separated into 2 groups – *X. crassiusculus* alone and *X. crassiusculus* plus the total number of unknown species collected at the same time.

**2.12 Results:** While the total number of *X. crassiusculus* collected/trapped were low, the UVMT sides caught significantly more ambrosia beetles (83%) than any of the 3 yellow hues tested (17%) indicating that yellow, as suspected, is not a preferred host regardless of hue, and actually may be acting as a repellent (see below) (Fig. 16). The UVMT was highly attractive to the ambrosia beetles – *X crassiusculus* - and the unknown species, which differed by an order of magnitude, despite having only a quarter of the traps’ surface areas to choose from and the lure on the top equally attracting potentially all 4 sides per the box-shaped UVMT (Fig. 17). This result raises the question as to whether singularly or in combinations (push-pull) UV mulch as an attractant and/or yellow as a non-preferred, perhaps repellent host, could be used to protect host trees from attacks by ambrosia beetles (see below). Therefore, the idea was pursued as follows using several approaches.

**Figure 16:**
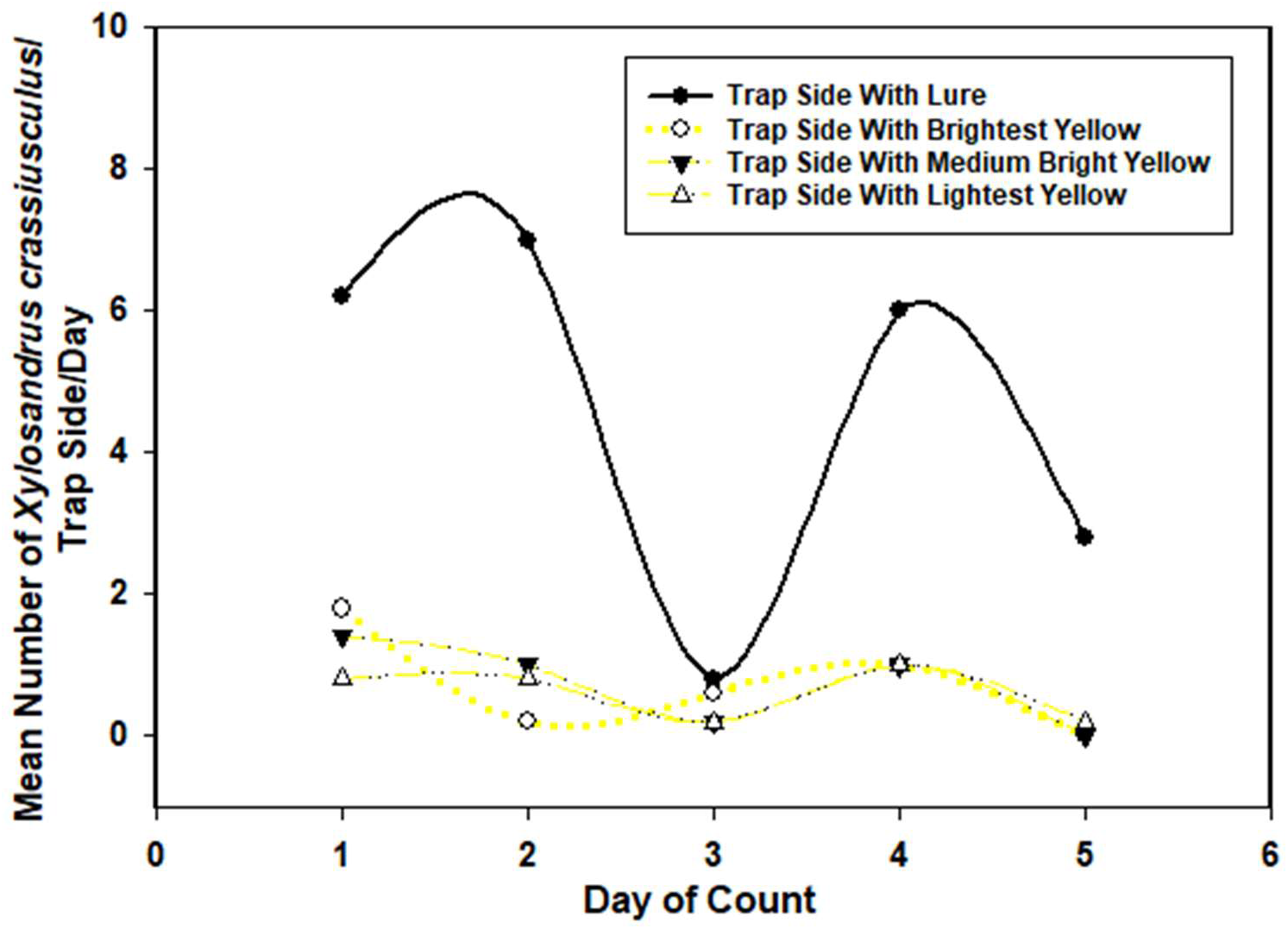
Response of *X. crassiusculus* to a square UV mulch trap containing 3 single-sides with hues of yellow – bright safety yellow, medium bright yellow and a whitish yellow - and a UV mulch side with an ethanol lure. All sides contained Tangle trap for beetle collection on a 10 x 30 cm space in the middle top of each side.

**Figure 17:**
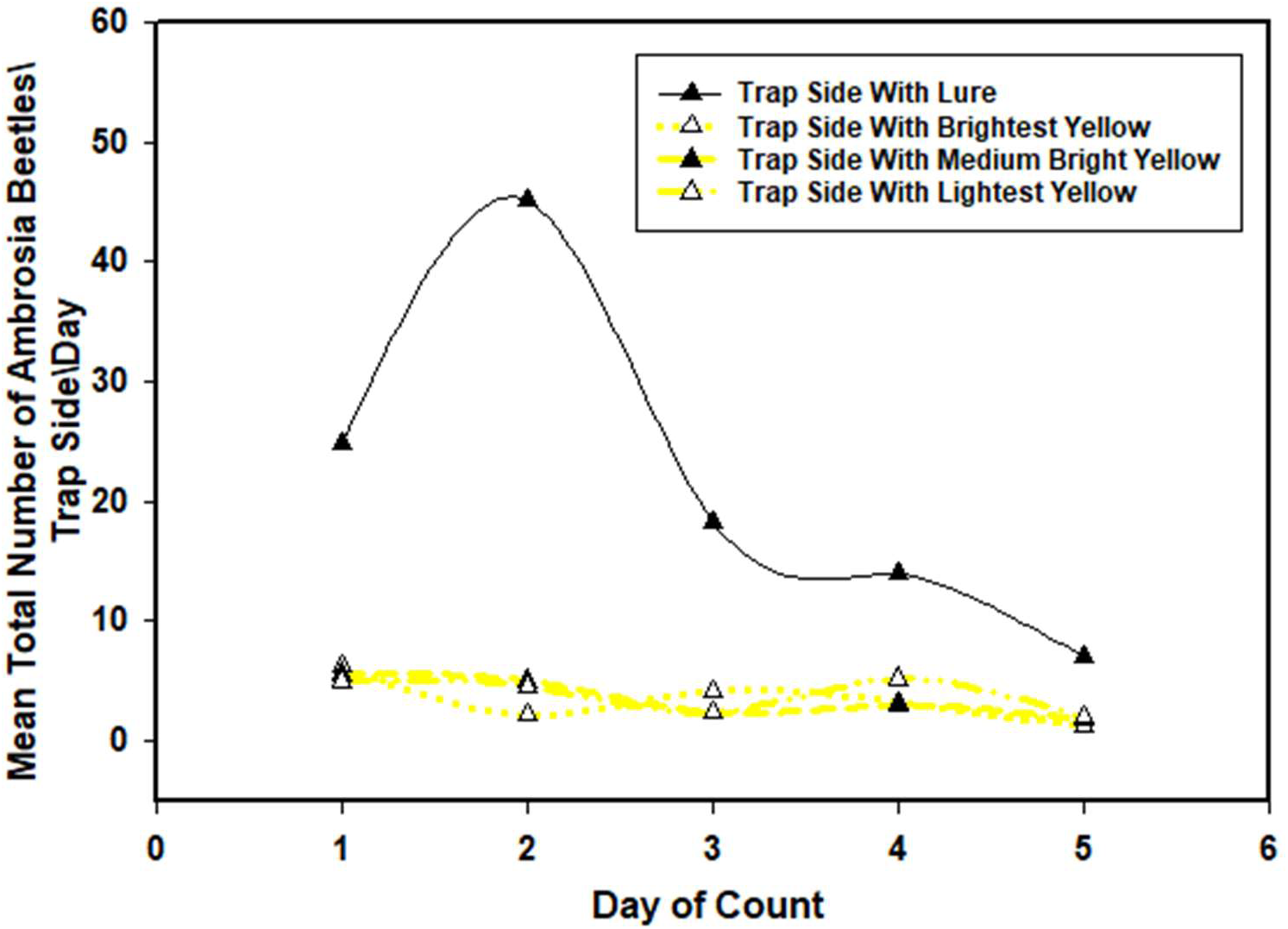
Response of the total number of ambrosia beetles (multiple species) to a square UV mulch trap containing 3 single sides with hues of yellow – bright safety yellow, medium bright yellow and a whitish yellow - and a UV mulch side with an ethanol lure. All sides contained Tangle trap for beetle collection as above.

#### 2.13: New ideas

As a follow-up to the above experiment to learn more about the response to yellow, 3 UVMTs were set up to have the opposite pattern on the 4 sides of the traps as follows: 1. Side 1 and side 2 with UVM only, 2. side 3 with UVM + the ethanol lure (control), and 3. side 4 with safety yellow. Each of the trap sides contained a small piece of UVM (10 x 31cm) in the middle covered with Tangle trap to capture the responding beetles. Beetles on the traps were counted every other day for 7 days and the traps were turned clockwise 1 cardinal direction at each count. The results are presented for *X. crassiusculus* and the unidentified other ambrosia beetle species that were also captured and counted.

**2.13 Results:** The capture rate was high for both types of ambrosia beetles captured (Fig. 18, 19). The results indicate that the 2 sides with UVM only had the highest number of both counted beetle types whereas, the UV mulch plus lure was much lower and closer to the yellow side which captured the lowest numbers of beetles. Therefore, the yellow maintained its consistent negative effect on the beetles while the UVMT captured the highest number of ambrosia beetles on the sides without lures. This is notable from previous experiments, but not unexpected, given that the UVM material always captures some beetles on sides of the trap that do not have the lure (Fig. 18 and 19).

**Figure 18:**
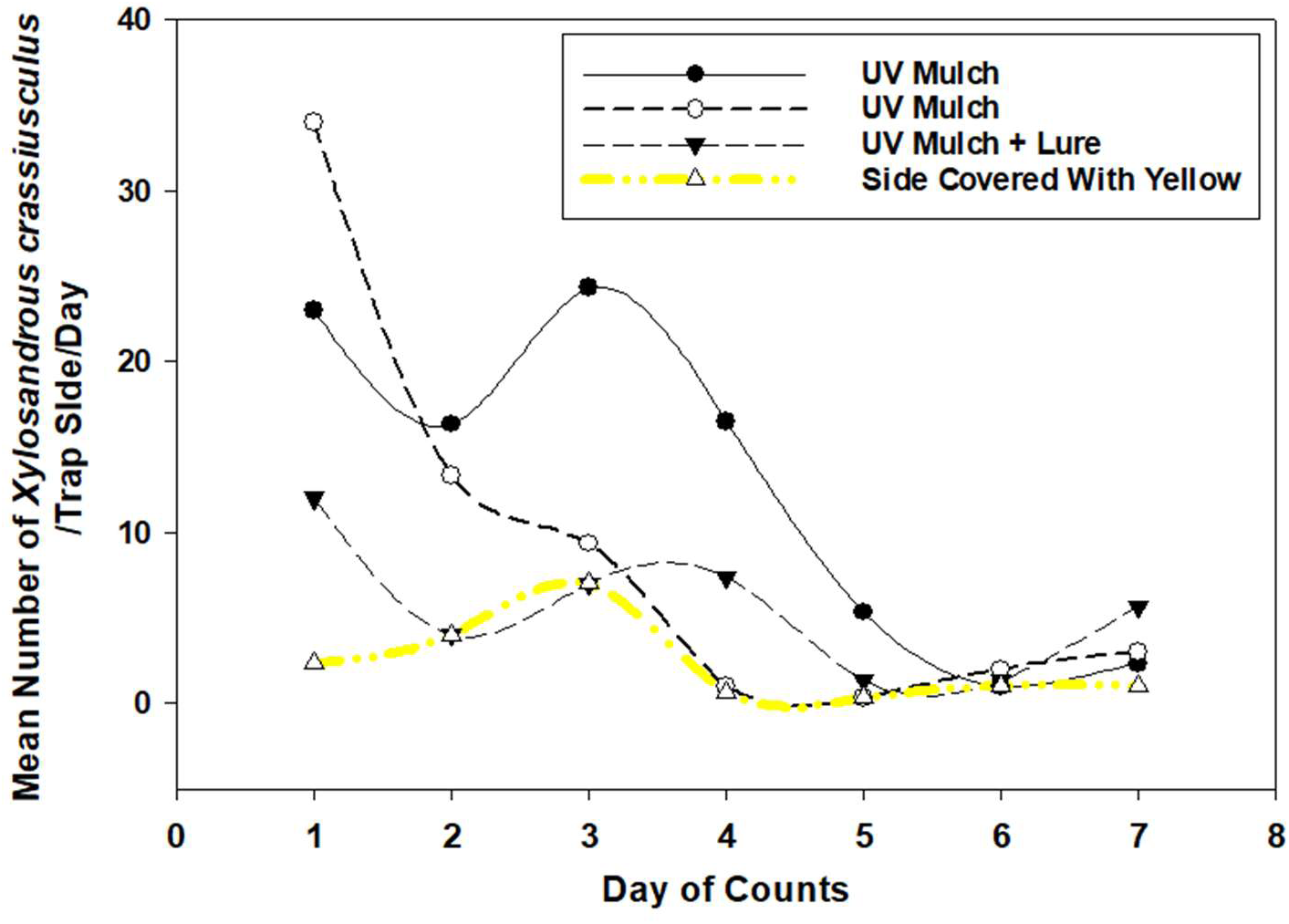
Response of mean number of *X. crassiusculus* to a square UV mulch trap containing 1 yellow – bright safety yellow – side, 2 UV mulch sides without a lure and a UV mulch side with an ethanol lure. Each side also contained a small piece of UV mulch, 10 x 31cm. with Tangle trap to collect the beetles.

**Figure 19:**
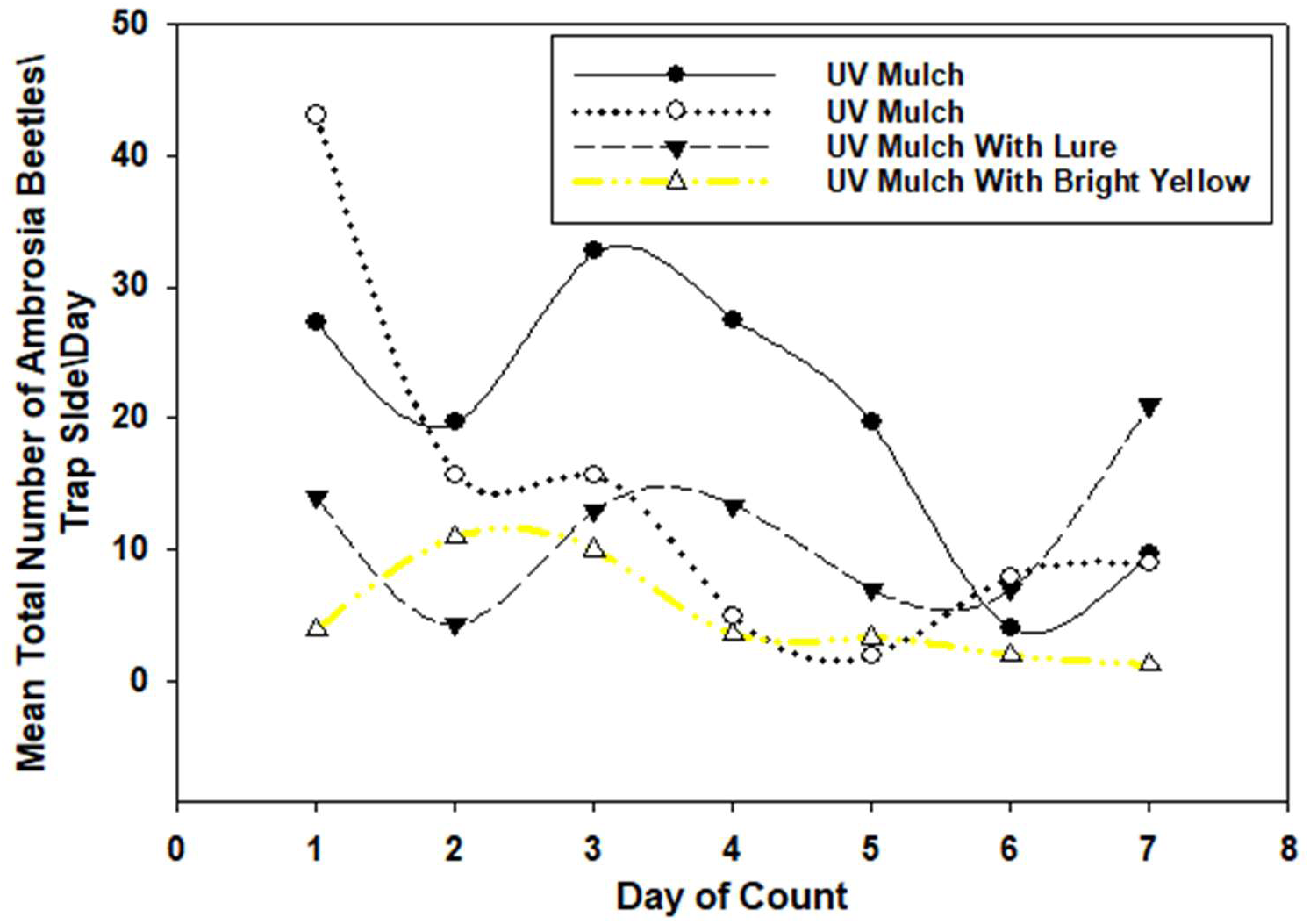
Response of mean total number of ambrosia beetles to a square UV mulch trap containing 1 yellow – bright safety yellow side, 2 UV mulch sides without a lure and a UV mulch side with an ethanol lure. Each side also contained a small piece, 10 x 31cm, of UV mulch with Tangle trap to collect the beetles.

#### 2.14: Response to yellow and metallic silver on beetle host trees using fresh tree bolts to compare beetle response to colors

Given the positive beetle response to the UVMT, and the opposite, much lower response to the hues of yellow colors tested, an obvious question arises. It’s one thing to observe selection of one color treatment over another in a non-natural test situation, e.g., UVM is much more attractive than yellow. However, it is another to observe the same effect under a natural (tree or bolt) situation, despite previous experiments with traps. For this situational question, refer back to the methodology, ethanol-induced fresh tree bolts, used in the early insecticide tests in part 1 of this manuscript. Question: What are the potential effects for use of fresh bolts and have yellow manifesting a similar result using the non-native host, mimosa or silk tree (*Albizia julibrissin* Durazz), and a native host, sweetgum (American sweetgum (*Liquidambar styraciflua* L., a native plant to North America))? Note, the previous experiments using the coroplast rectangles described above for *X. glabratus/X. crassiusculus* indicated that yellow and silver colors provided opposite behavioral responses from the beetles. Both types of trees are attacked by *X. crassiusculus* and other ambrosia beetles naturally when alive or cut down in Florida. Thus, to follow up on this result, fresh bolts from natural host trees were colored and compared. Bolts of each of the trees were prepared as indicated in Part 1 by using small tree bolts 47cm long and from ∼5-7cm in diameter. Treatments of the bolts were: 1. untreated controls, surface remains just natural bark, 2. bolts sprayed with metallic silver spray paint (equivalent to UVM as shown above), and 3. bolts painted with yellow paint (safety yellow as above). All test bolts were placed upright (end of bolt closest toward original tree roots) in a small yogurt cup containing 10-15% ethanol in water and attached with a plastic marker tape to a line of wire between trees in the forest raised above ground such that the tops of the bolts could be attached easily to the wire. Four reps were used for each treatment. Numbers of beetle galleries/bolt as indicated by appearance of frass (match like) material were counted once per week for 4 weeks of data. Two mimosa experiments were conducted along with three using sweetgum as shown (Figs. 20-24).

**Figure 20:**
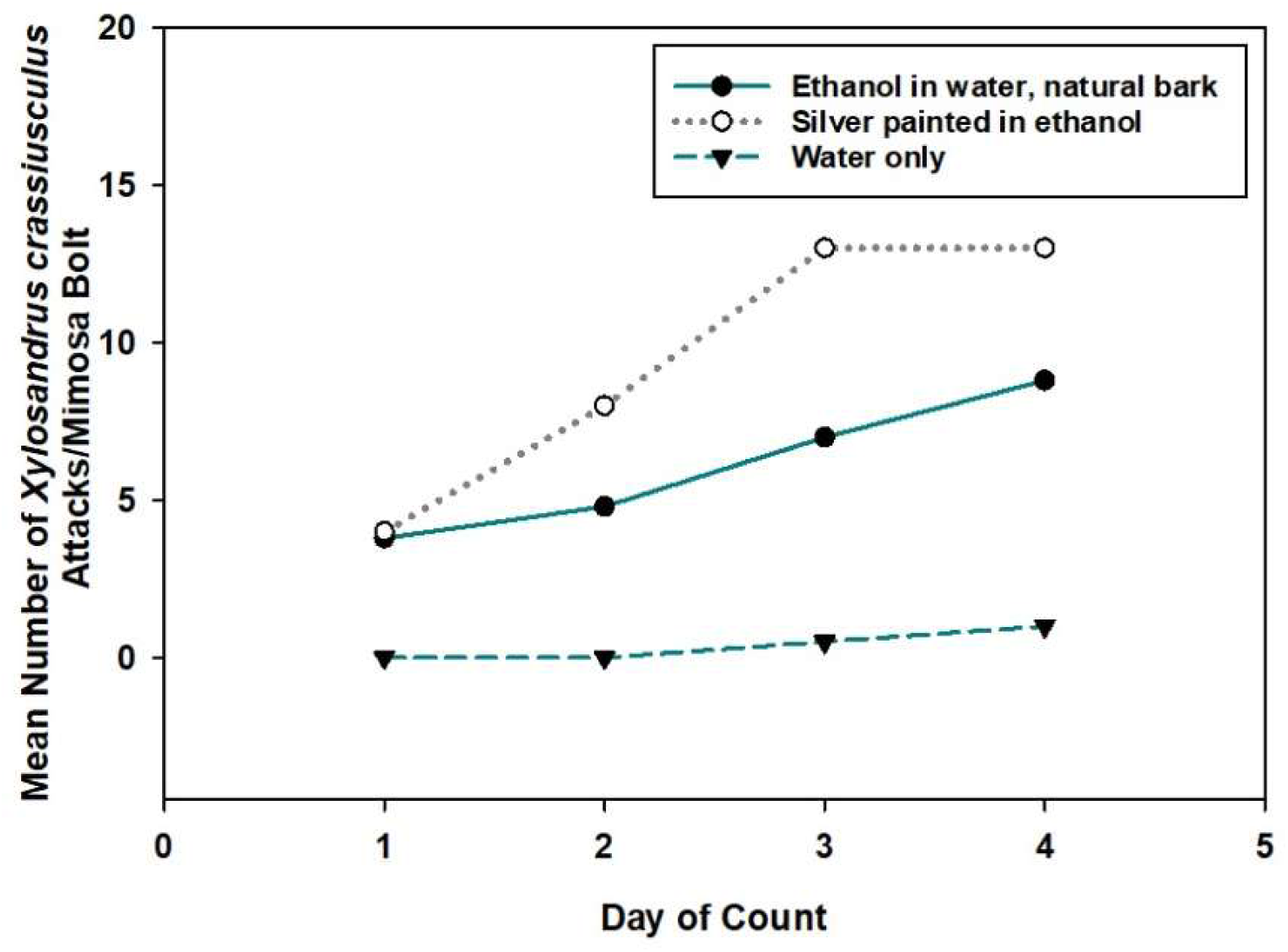
Response of mean number of *X. crassiusculus* to mimosa bolts in 3 treatments: 1. a natural (e.g., just bark) water control, 2. a natural 10% ethanol and water standard, and 3. bolts painted metallic silver standard that matched UV mulch in color and an ethanol lure.

**Figure 21:**
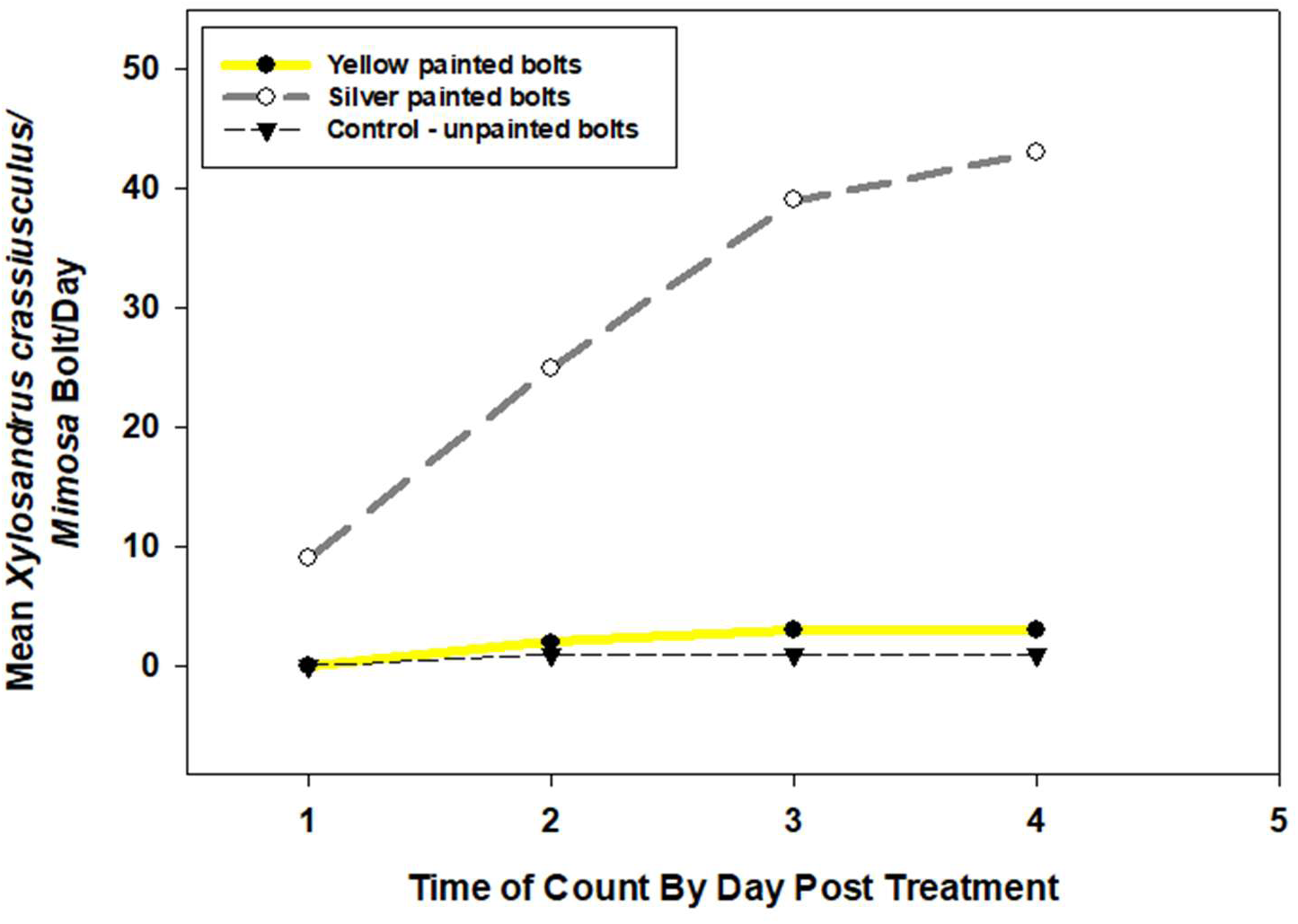
Response of mean number of *X. crassiusculus* to mimosa bolts in 3 treatments: 1. a natural (e.g., just bark) ethanol and water standard (10% solution), 2. yellow painted bolts in an ethanol and water standard, and 3. bolts painted metallic silver that matched UV mulch in color and an ethanol lure.

**Figure 22:**
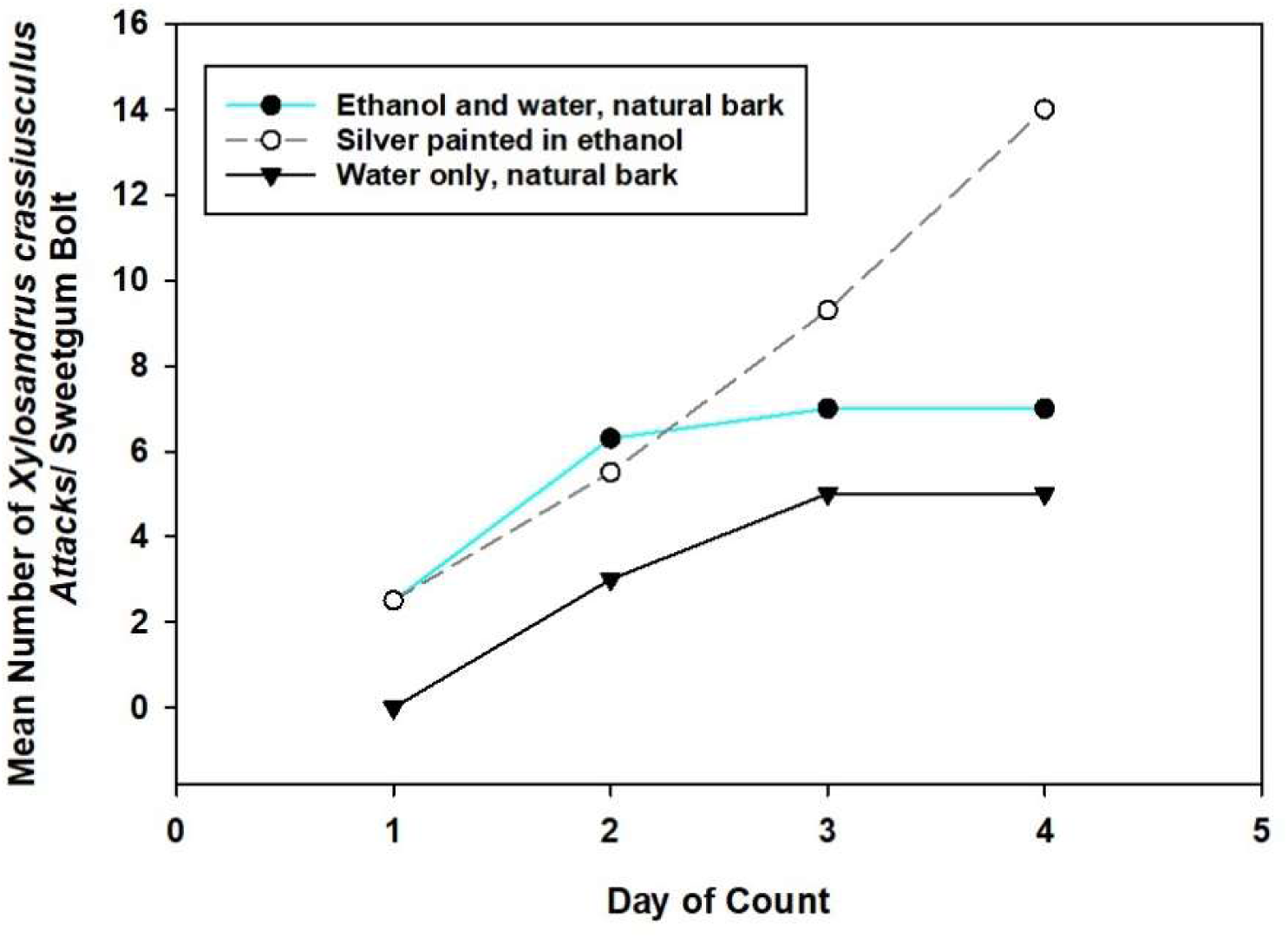
Response of mean number of *X. crassiusculus* to sweetgum bolts in 3 treatments: 1. a natural (e.g., just bark) 10% ethanol and water solution, 2. bolts painted metallic silver that matched UV mulch in color in an 10% ethanol/water standard, and 3. Yellow-painted bolts and an ethanol lure.

**Figure 23:**
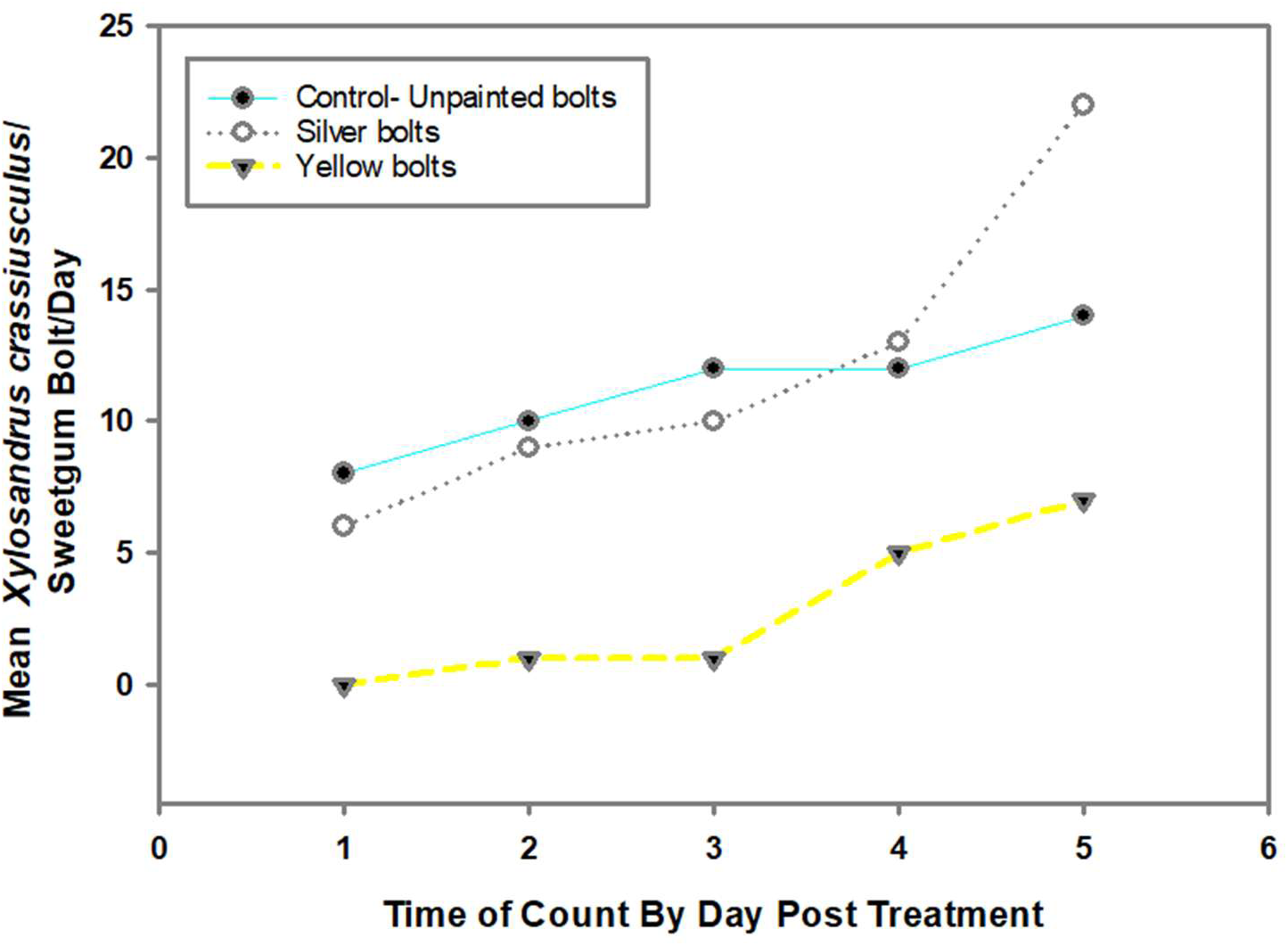
Response of mean number of *X. crassiusculus* on sweetgum bolts in 3 treatments: 1. a control natural (e.g., just bark) 10% ethanol-water standard, 2. Bolts painted metallic silver that matched UV mulch in color and an ethanol lure, and 3. yellow bolts in an 10% ethanol-water.

**Figure 24:**
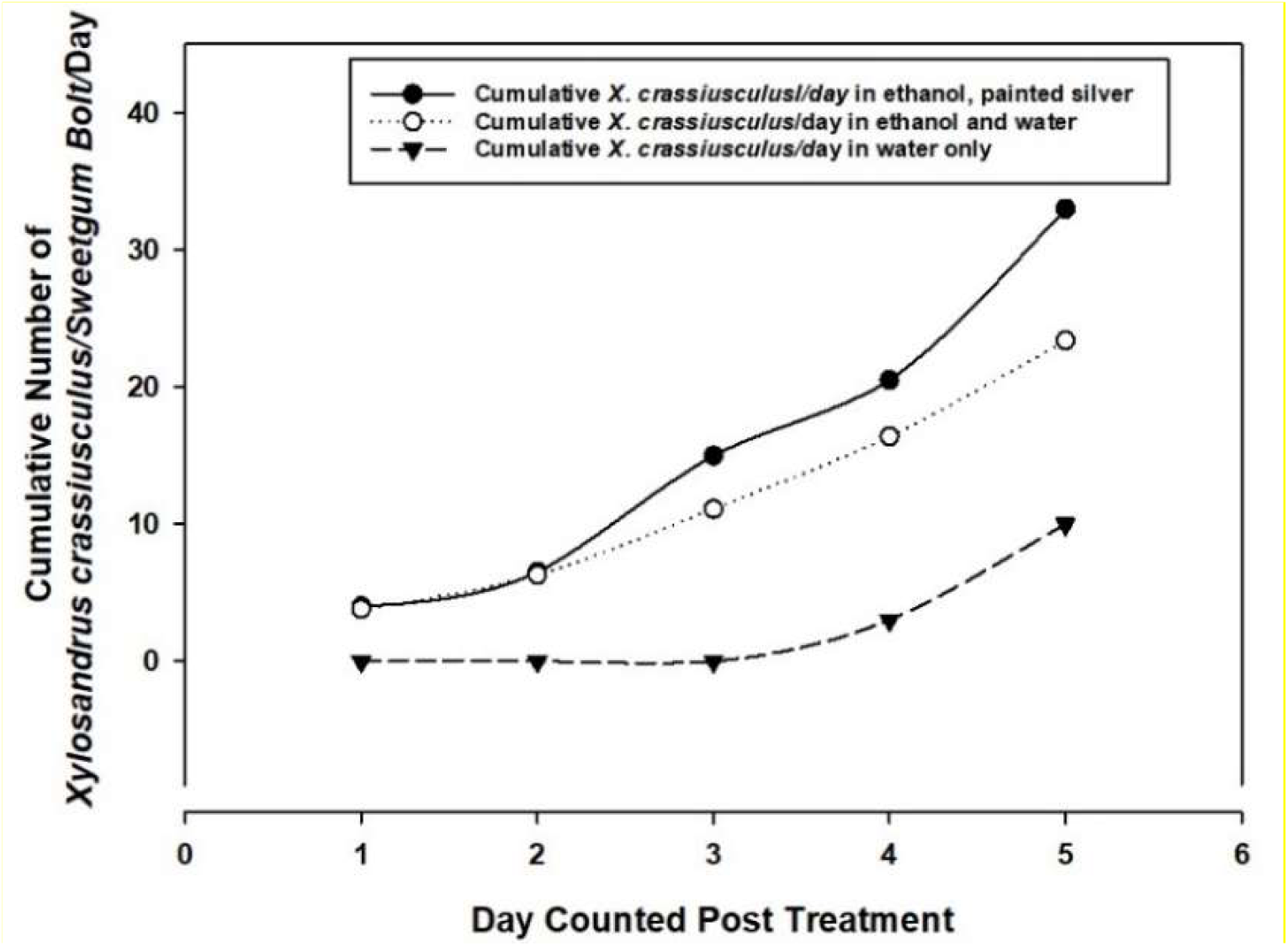
Response of mean cumulative numbers of *X. crassiusculus* to sweetgum bolts in 3 treatments. 1. bolts painted metallic silver that matched UV mulch in color in a 10% ethanol/water standard, 2. a natural (e.g., just bark) and an ethanol lure, 3. bolts in water alone.

**2.14 Results:** In this experiment the bolt treatments were formed with spray paint and the paint as a physical barrier had the potential to reduce the number of attacking beetles. However, the total number of attacks among treatments and reduction in attacks/treatment were consistent across all color experiments either positive (silver) or negative (yellow) or by both evaluation tools, e.g., UVMT (as above) and tree bolts (Fig. 20 - 24). The results are straight forward and are self-explanatory (Fig. 20-24). Basically, the numbers are higher overall, the results are the same for both host plant species and consistent in the comparisons of the colors and UVM. UVM attracts while yellow repels! Thus, “It appeared that a trap was born and an IPM tool was discovered”! The next section follows from this result and identifies another novel but somewhat similar vision-based manipulative approach and questions.

#### 2.15: Effects of specific color patterns via use of camouflage materials on ambrosia beetle response to a UVMT

The consistent positive results above with mimosa and sweetgum trees painted silver and the lower beetle response to yellow bolts elicited the following questions: 1. What is the underlying explanation for the heretofore, unrealized behavior by ambrosia and bark beetles (see below) to the visual spectrum from UV mulch?, 2. Could this observed behavior of enhanced attraction to UV mulch and/or reduced visual response to yellows be exploited alone or together enough to protect host trees (again, push-pull)?, and 3. If so, are there other available patterns and tools that could be discovered and exploited to test and increase the effects such as provided by the color yellow that would broaden the usefulness of potential IPM tools? These ideas were partially raised by the use and opposite effects from UV blocker material previously described above on color blind Cervidae and ambrosia beetles.

As an initial experiment related to the above questions, the comparison of the visual cues available and possibly used as well as avoided by ambrosia beetles during their searching flights through the forested landscape for host tree odors was considered as a potential starting point. From that idea and others (see below), the potential value of camouflage (camo) used by many other organisms as well as by human game hunters, was considered as another potential tool and pathway to more information and perhaps answers toward an explanation for the efficacy of the new discovery. Camouflage cloth and clothes are readily available as are mostly any other products from shirts and hats to tarps, blankets, etc., all often available in many different patterns. Hunters of wild animals such as wild turkey, deer and other Cervidae, etc., wear camouflage materials to cover themselves and to blend into the various landscapes they hunt in at different locations and times during the year. Of course, most everyone should be aware that many insects use camouflage as protection against predation.

These features surrounding the use of camouflage raise questions about how an ambrosia beetle that spends the great majority of its life inside a tree trunk and a very small percentage of that life spent visually navigating between host trees, might evolve a behavioral approach “using” and “avoiding” (my key words) the habitat factors presented in navigating hardwood/conifer forests. Because yellow-green is a major part of the visual spectrum offered by leaf-covered trees in a forest (Enders 1993), getting through the trees is an obvious barrier to a moving beetle searching for an odor emanating from a dark-colored tree trunk. This raises the question: do the beetles pick the best path through the forest vegetation by avoiding the yellow color of hardwood leaves (demonstrated above, but less than 100% effective) which has been shown to be non-preferred at minimum. And in fact, because of avoiding yellow-green leaves, substitute the light spectra available in the latter parts of a day as the main visual factor (as either an opening or an attraction?) and operate the navigation factor in the hunt for the kairomones emanating from target host trees?

To develop potential information on this subject matter, different types of camouflage with patterns that might blend in with a hardwood forest whose colors were yellow, green, brown or similar, and also ones with patterns similar to hardwood tree leaves (either green or brown) were obtained. See supplemental material pictures for the camouflage patterns used. Camouflage is available in countless patterns from all kinds of stores. Some of these are UFPro.com. www.fabric.com, The Party Store, JoAnne™ Fabric and Craft, Amazon, Ineedfabric.com, FieldsFabric.com, WayFair and many stores on ETSY.com, etc. DPM which is “disruptive fabric material” is a common name for various available patterns.

The following efforts use the above tools in a mix of comparative tests to determine the best combinations of “tools” (now termed as a result of the above previous results) along with how they operate! Using camouflage materials as the treatments and the UVMT, the experiments and methods were repeated in large measure as they were conducted using the 3 hues of yellow as above. With these treatments, a smaller piece of UVM (10 x 20cm) covered with Tangle-Trap™ was only placed in the center of the camouflage and UVM treatments that, unlike in the yellow coroplast traps, was coated over the entire treatment sections. As usual, the UVM side was used as the control in comparison with the camouflage treatments placed on the other 1-2 or 3 sides of each UVMT. An ethanol lure was added onto the top of the traps or as otherwise indicated. Four different combinations of camouflages, yellow paint, and UV mulch were used as treatments. Different numbers of traps were used in the individual experiments, beetles were counted, evaluated as *X. crassiusculus* or total number of ambrosia beetles (*X. crassiusculus* is the unidentified species total) and at each count the beetles were removed and the Tangle trap cleaned and replaced as needed.

##### 2.15.1: A treatment in yellow, 2 in camouflages and the UVMT as control and a way to manipulate and evaluate the results of the materials

An experiment with treatments as follows: a control UVMT with the ethanol lure, a side with DPE green-yellow beaded-type camouflage, a side with Mossy Oak camo (autumn leaves and limbs) and the fourth side in safety yellow. Four such traps were used and counted 2 days apart over 4 dates.

**2:15.1 Results:** The experiment lasted 8 days and the results are represented in 2 figures. The control – the UVMT + lure side - had a higher overall average capture rate of *X. crassiusculus* (Fig. 25) and even higher numbers with the total counts (Fig. 26). The numbers of beetles captured on the other three sides were both similar and much lower (by order of magnitude for the *X. crassiusculus* plus the unidentified total). These figures indicate that both camo types elicited the same response that the yellow treatments did from multiple species of ambrosia beetles similar as found within the figures above with both colored treatments on traps and bolts from different host plants.

**Figure 25:**
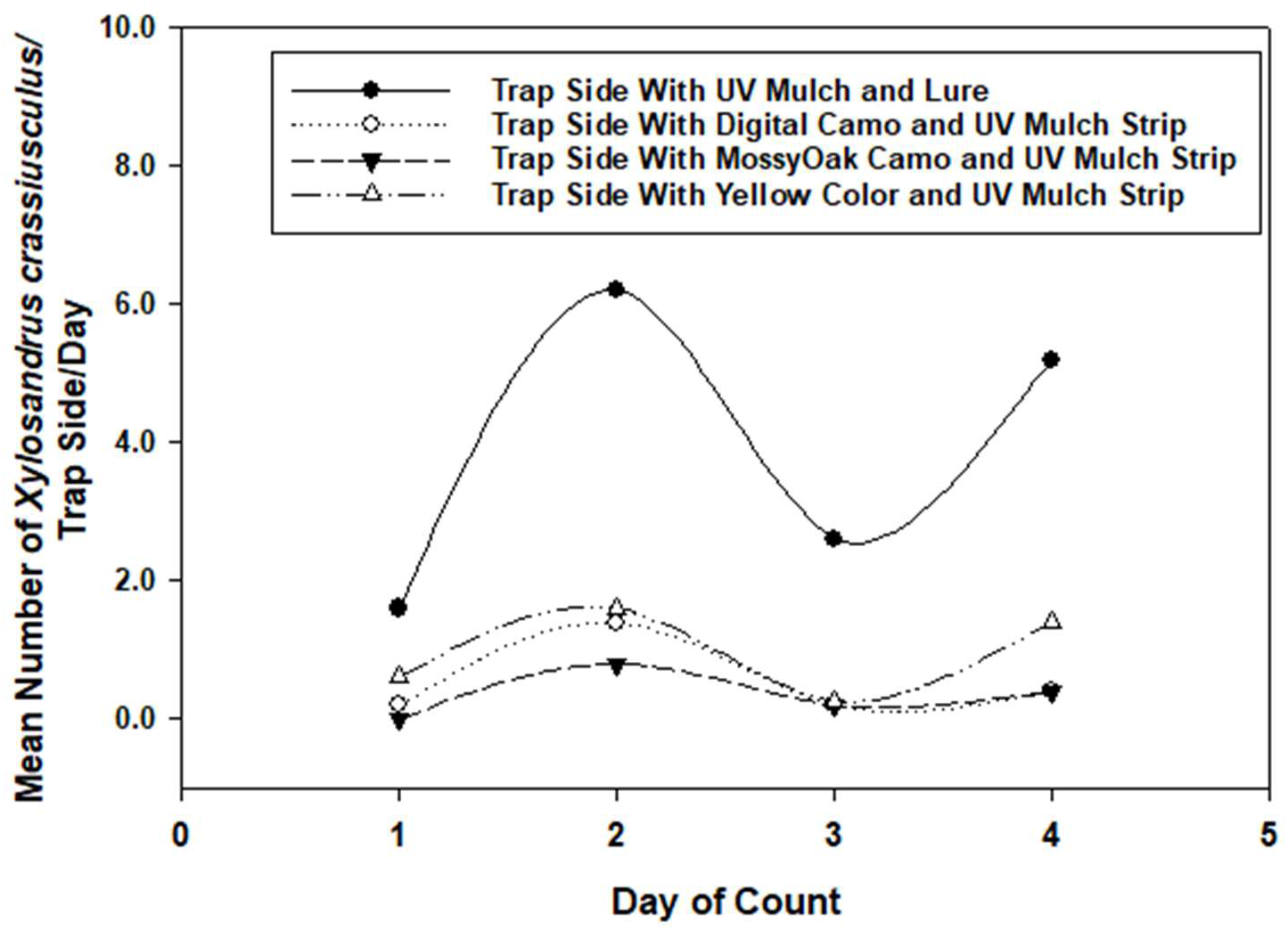
Response of mean number of *X. crassiusculus* to a square UV mulch trap containing: 1. 1 yellow – bright safety yellow – side, 2. a digital camouflage covering, 3. a mossy oak camouflage covering, and 4. a UV mulch side and an ethanol lure. Each side also contained a small piece (7.5 x 31cm) of UV mulch with Tangle trap.

**Figure 26:**
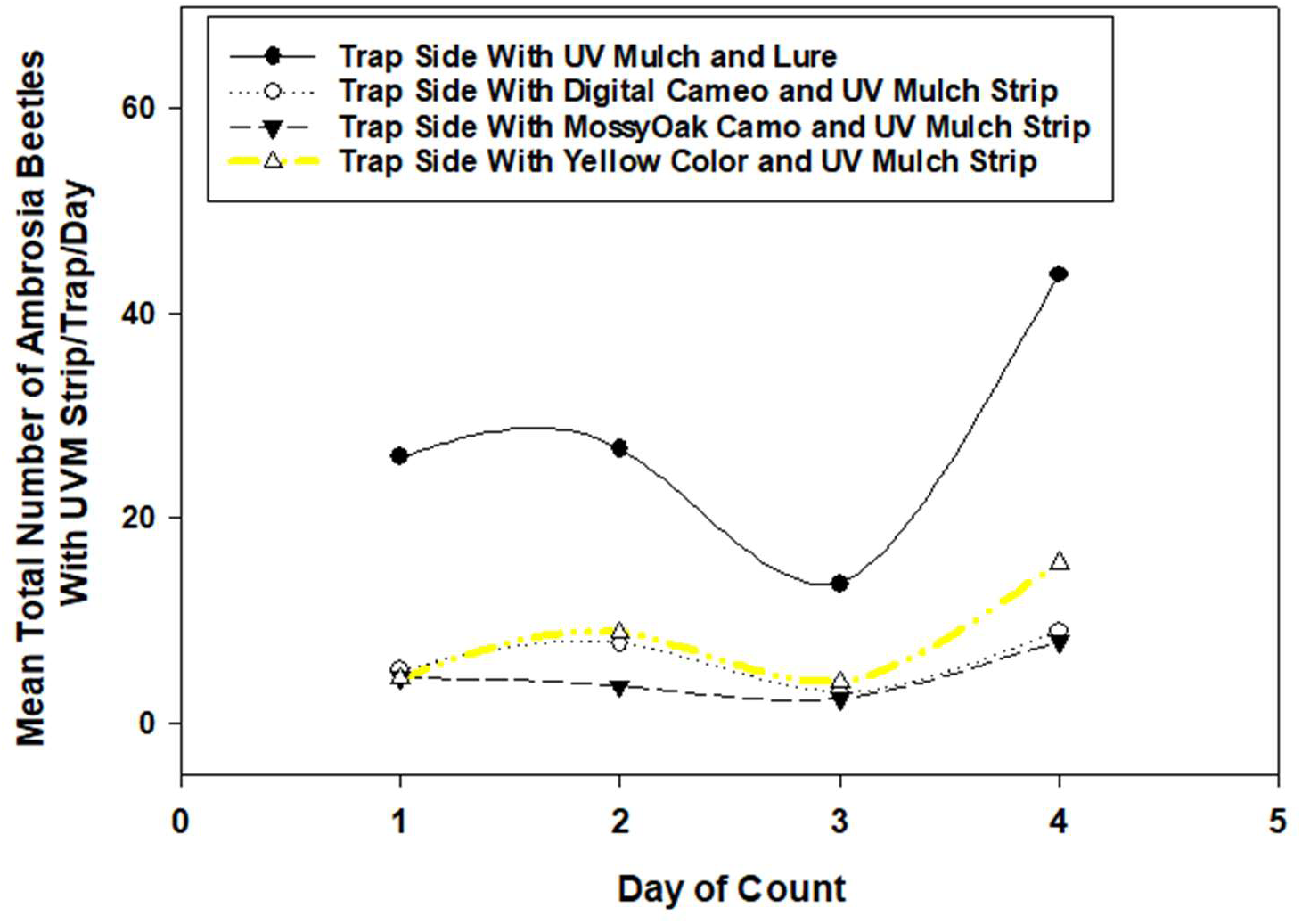
Response of mean total number of ambrosia beetles to a square UV mulch trap containing: 1. a UV mulch side with an ethanol lure, 2. a digital camouflage side, 3. a mossy oak camouflage covering, and 4. a bright safety yellow side. Each side also contained a small piece of UV mulch with Tangle trap. Clearly, the small patches of UV mulch in the center of the trap sides did not affect the treatment and results as expected.

##### 2.15.2: DPE camouflage plus lure, absorbent, reflectent and UV blocking sunscreen

This experiment compared the results of DPE camouflage plus lure, and the 3 suncreen treatments: absorbent, reflectent and the UV Blocker. Five traps were used and counted 2 days apart over 6 counts. As before, the captures were collected as *X. crassiusculus* and total (*X. crassiusculus* plus unidentified species of ambrosia beetles). The lure was placed on the DPE camo to determine if the combination would increase the capture rate of the camo.

**2.15.2 Results:** No effect occurred. The experiment lasted 12 days and is represented by Fig. 27. The DPE camo plus a lure had no effect on the camo capture rate, it remained low as expected. The reflectent treatment had a higher overall average capture rate of *X. crassiusculus* and even higher numbers with the total counts (data not shown). From the number of beetles captured on the other two sides, the adsorbent was the lowest overall, and the UV Blocker was overall close to the reflectent as expected since it has no negative effect on ambrosia beetle vision behavior. These outcomes indicate that the treatments when compared as above, regardless of minor changes to the treatments, e.g., addition of the lure to the camouflage evokes consistent beetle response). Moreover, the treatments actually worked better in this experiment, likely due to weather rather than like the results shown in the previous experiments with those treatments (e.g., the reflectent sunscreen). Again, the unidentified beetle species behaved as expected mimicking the *X. crassiusculus* behavior here and above.

**Figure 27:**
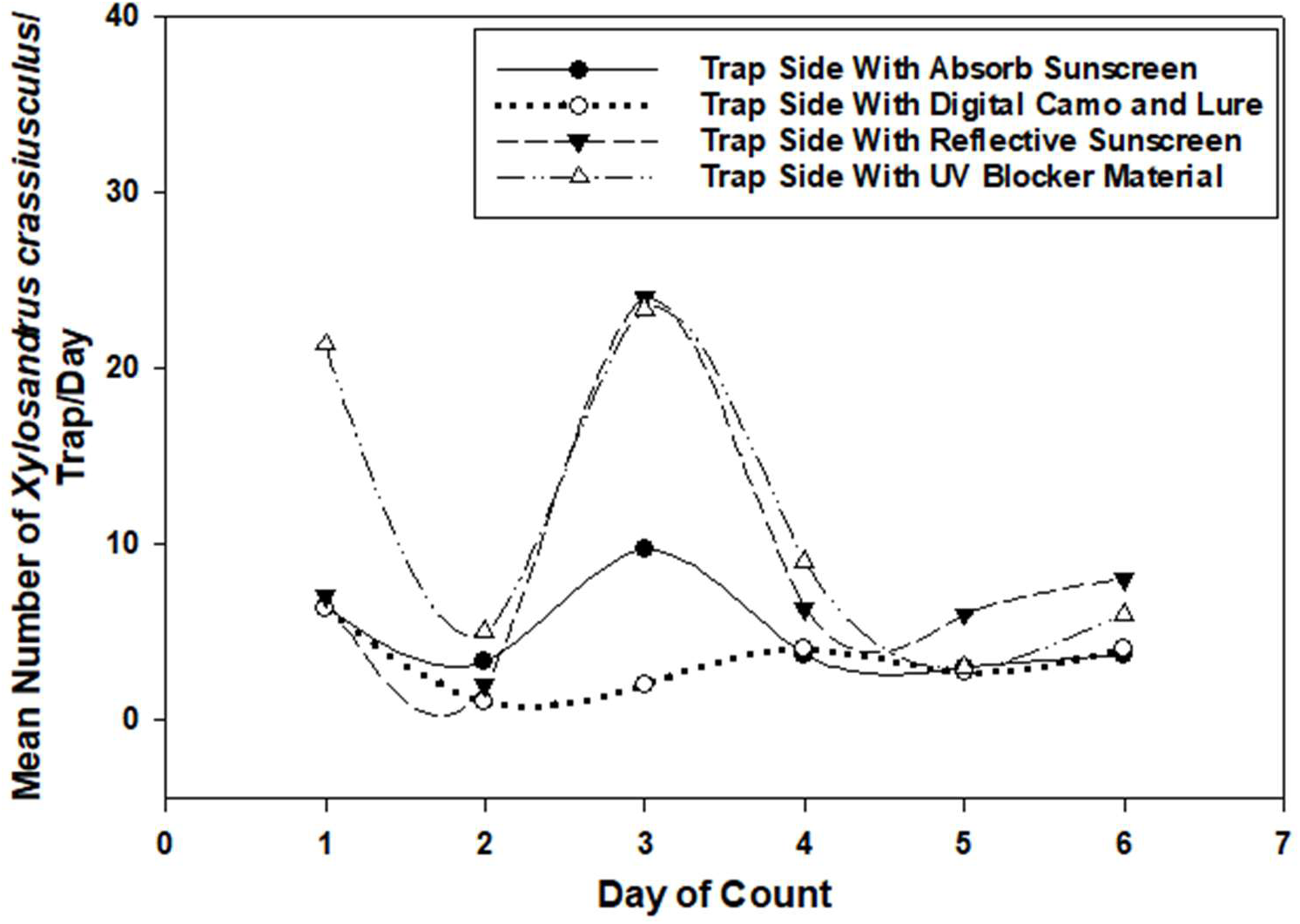
Response of mean number of *X. crassiusculus* to a square UV mulch trap containing: 1. a UV mulch side with an absorbent sunscreen, 2. a digital camouflage, 3. A reflective sunscreen, and 4. UV blocker material. Each side also contained a small piece of UV mulch with Tangle trap as above and an ethanol lure on top.

**2.15.3:** This experiment compared the treatments together of DPE camouflage plus absorbent, the UVM with absorbent, the UVM side with the lure, and safety yellow color. Four traps were used and counted 2 days apart over 6 counts. As before, the captures were collected as *X. crassiusculus* and total (*X. crassiusculus* + unidentified ambrosia beetles).

**2.15.3 Results:** Numbers of ambrosia beetles were relatively low (Fig. 28) but the results were the same as in previous experiments with these treatments as expected for both *X. crassiusculus* and the total number of ambrosia beetles. UVMT plus the lure captured more than the other treatments for both groups: the *X. crassiusculus* and the total of adding the rest of the unknown species collected (Fig. 29).

**Figure 28:**
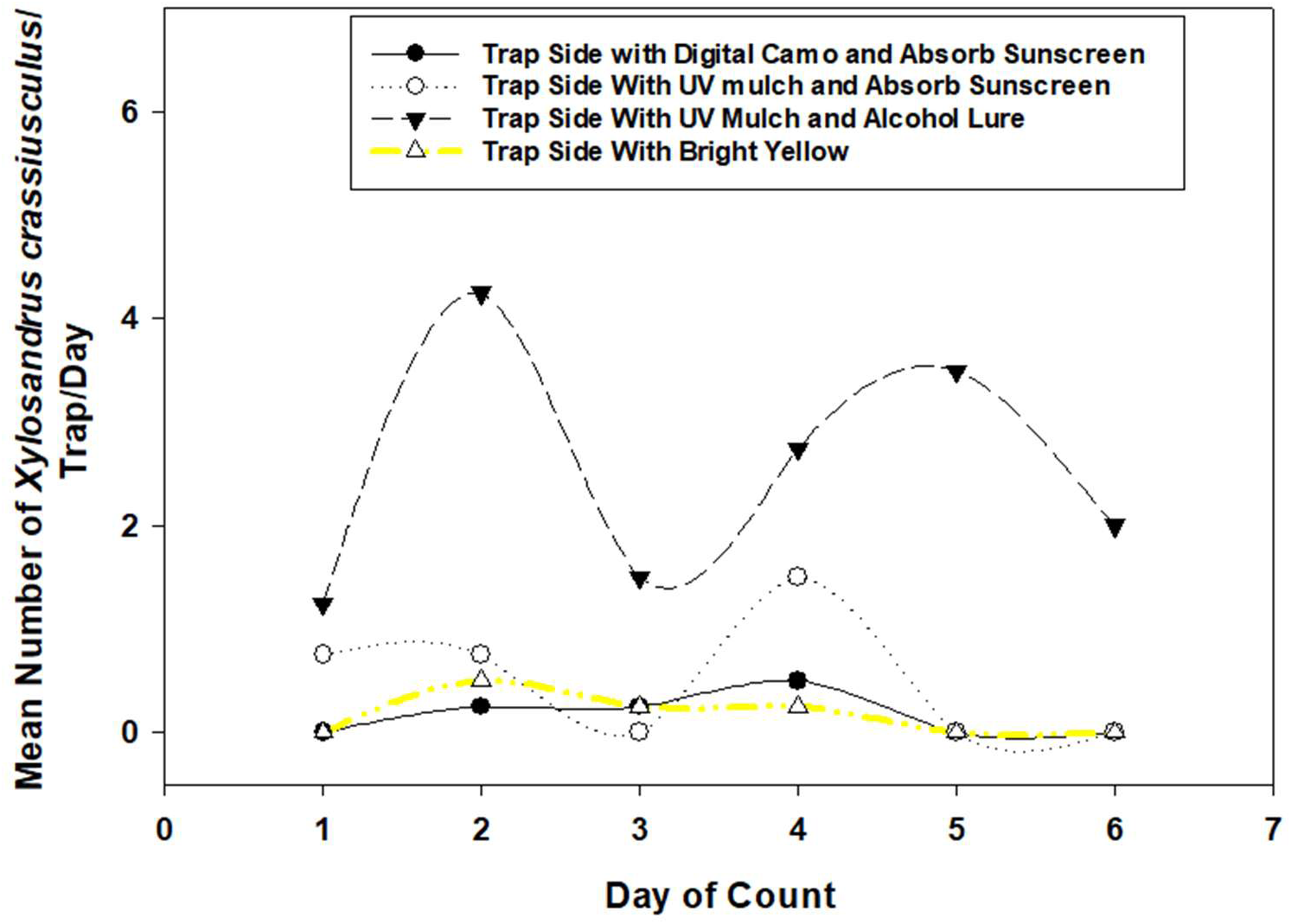
Response of mean number of *X. crassiusculus* to a square UV mulch trap containing: 1. a digital camouflage covered with absorbent sunscreen, 2. a UV mulch side with absorbent sunscreen, 3. a UV mulch side with an ethanol lure, and 4. a side covered with bright yellow. Each side also contained a small piece of UV mulch with Tangle trap as above.

**Figure 29:**
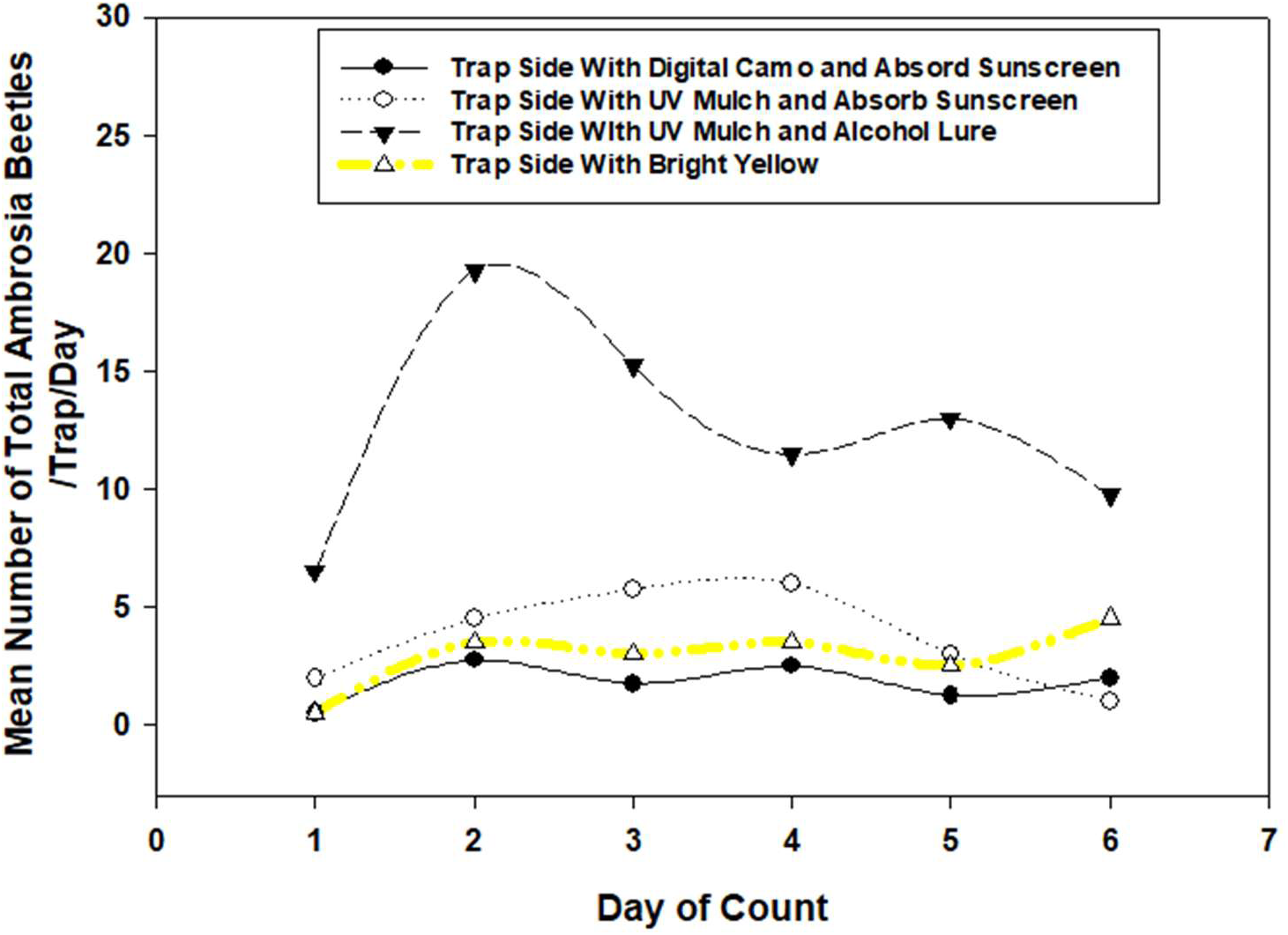
Response of mean number of *X. crassiusculus* plus unknown species (total) to a square UV mulch trap containing: 1. a digital camouflage covering with an ethanol lure, 2. a UV mulch side with absorbent sunscreen, 3. a UV mulch side with an ethanol lure, and 4. a side with bright yellow. Each side also contained a small piece of UV mulch with Tangle trap as above.

**2.15.4:** This experiment tested the UVM the lure standard control vs DPE and MossyOak camo and yellow. Five traps were counted over 5 days at 2 days apart. Counts were made of both *X. crassiusculus* and the total as above.

**2.15.2 Results:** All results were as expected with the UVMT + lure control the highest and other three treatments the lowest as expected (Figs. 30 and 31).

**Figure 30:**
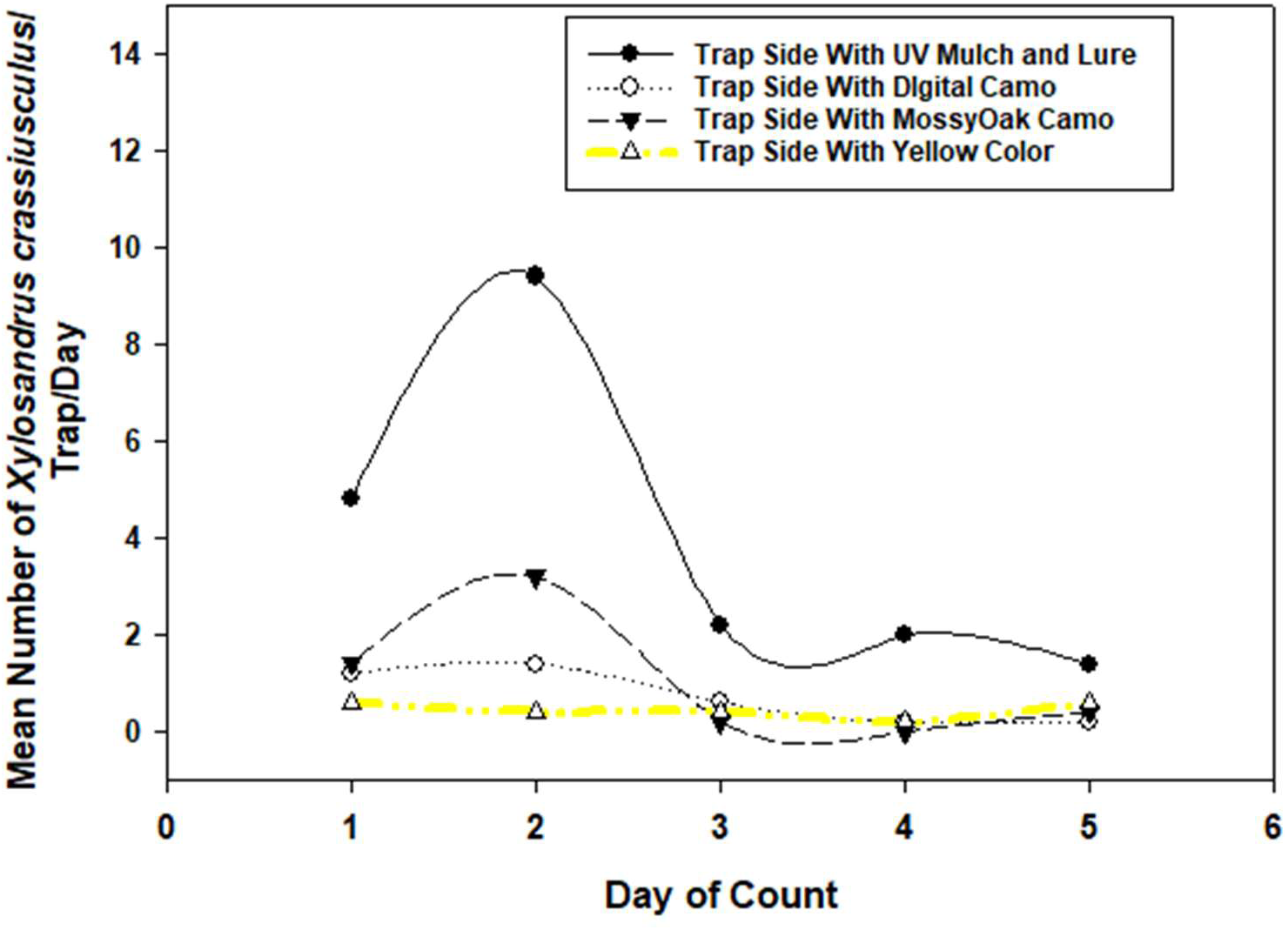
Response of mean number of *X. crassiusculus* to a square UV mulch trap containing: 1. a UV mulch side with an ethanol lure, 2. a digital camouflage covered side, 3. a mossy oak camouflage covered side, and 4. a bright safety yellow side. Each side also contained a small piece of UV mulch with Tangle trap as above.

**Figure 31:**
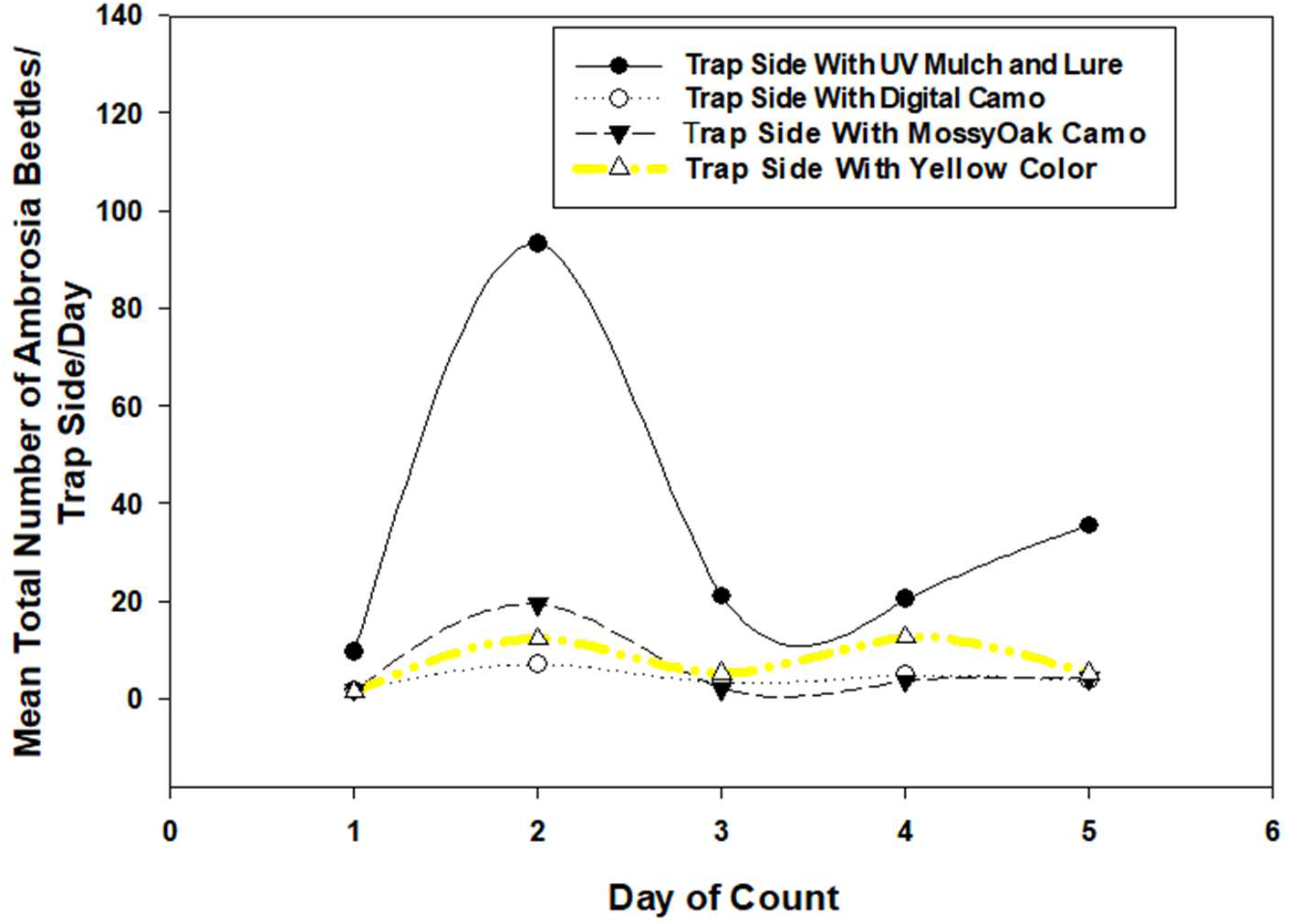
Response of mean total number of ambrosia beetles to a square UV mulch trap containing: 1. a UV mulch side with an ethanol lure, 2. a digital camouflage side, 3. a mossy oak camouflage side, and 4. a bright safety yellow side. Each side also contained a small piece of UV mulch with Tangle trap as above.

##### 2.15.5: Evaluating the previous colors and addition of testing burlap as an overall view of the above experiments

The above-described broad responses of ambrosia beetles to natural tree trunks and the fake tree trunks now termed UVM traps (UVMT) supported the testing of burlap fabric. Made primarily from jute fibers or hemp, burlap is a coarsely textured, durable, plain-woven fabric with usually a brownish color very much like certain tree trunks. Therefore, an experiment was conducted in north Florida in springtime whereby natural burlap, camouflage materials, yellow and green paints were tested on UVMT and compared for the responses of ambrosia beetles. The five treatments were arranged on 4 UVMTs/treatment by adding the treatments to the traps separated on 2 sides along with UV mulch on two separate sides. The fifth treatment was the control whereby all 4 sides were UV mulch. Four blocks of the 5 treatments were set up in a forested area with each block of traps separated from each other by 250m or more. Traps within each block were placed in the ground on the usual tomato stake ∼30m apart. Ambrosia beetles captured were mostly *X. crassiusculus,* with some unidentified species. An ethanol lure was placed on the top of the trap and a 21 x 25cm piece of clear plastic was stapled to each of the 4 sides on the UVMTs and covered with Tangle-Trap™. The traps were counted and cleaned once per week for 8 weeks. At each count the 5 traps/block were turned in one cardinal direction but not rearranged. Since burlap material was very attractive to the beetles, another treatment, burlap matching paint, was added in the sixth week and then counted on the seventh and eighth weeks. The data from the traps were processed to compare the numbers of beetles in the 4 treatments to UV mulch on each trap as well as to the UVMT - control trap. The results are provided (Fig. 32) to demonstrate the relative response of ambrosia beetles under the 2 methods. The data were handled separately for each of the eight dates as follows. The number of beetles counted on each of the 2 sides per 2 treatments (couplets) of each trap were added together into one count per trap/treatment. Those numbers for each day and treatment were added to produce a mean number per treatment per trap per day by couplets of each of the 5 treatments of each type: a UV mulch plus yellow, green, camouflage, burlap or 2 UV mulch (control). The mean number of beetles per the two treatments were added together and then the percentage of the numbers of the two treatment couplets were used in the figure to display the relationships.

**Figure 32:**
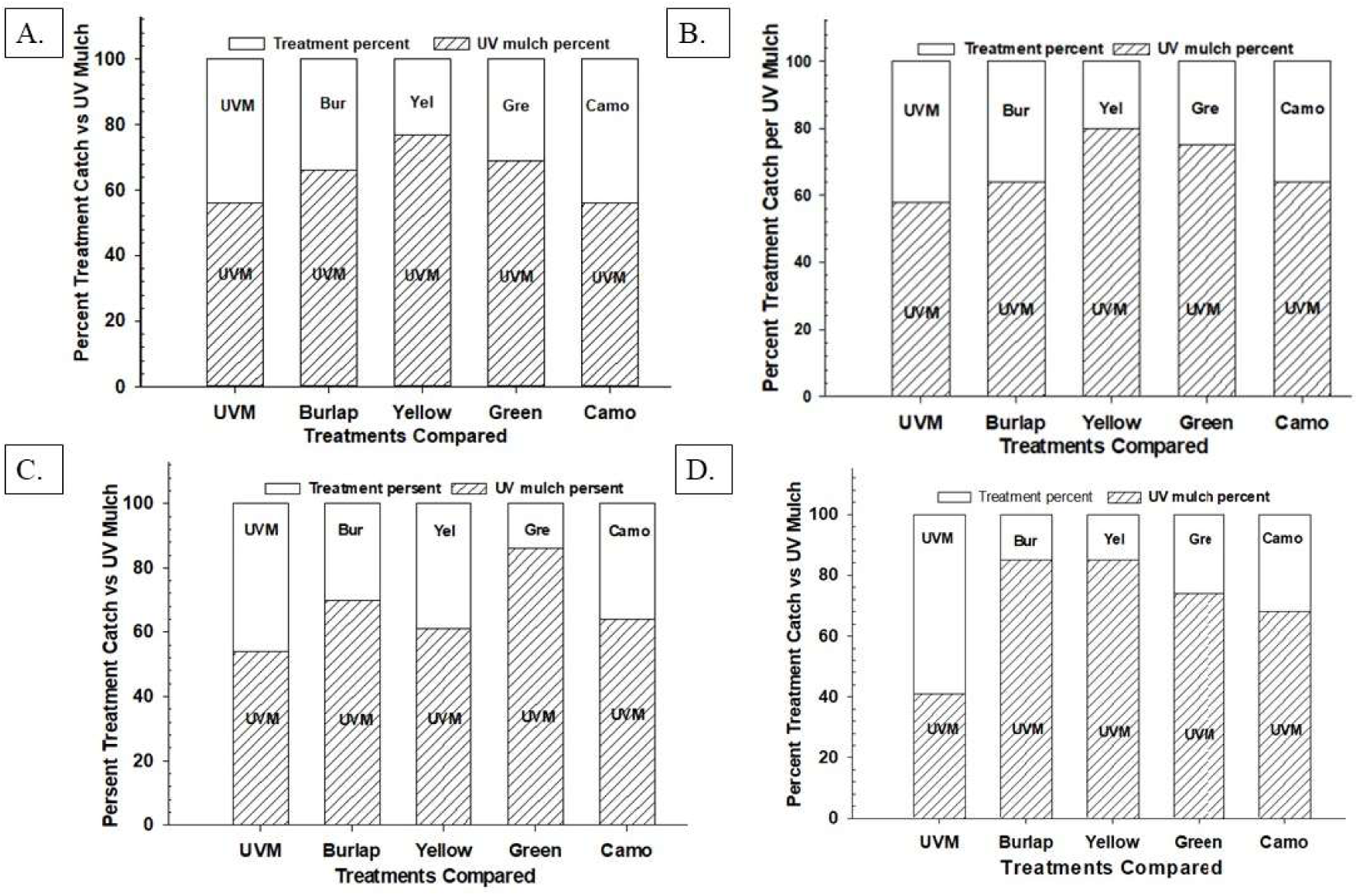
Field comparison of treatments selected by *X crassiusculus* using the UVMT. The UVMT treatments were set up so that 2 opposite sides of each trap contained UV mulch and the other two sides contained only one of the 5 treatments compared. The control traps had all 4 sides covered with only UV mulch. A lure containing ethanol (AgBio, Inc.) was placed on the top of each trap. The burlap paint was only tested during the last two weeks as shown in the figures. This is the first and only testing of the color green, burlap material and burlap-like paint in this text. Count data from each week was added together from each trap by date, block, and trap type and then the mean number of beetles captured per date, per block per trap by treatments were calculated. The mean total numbers of *X crassiusculus* from each or the treatment pairs were turned into percentages of the totals which appear on the figures to show the levels of comparisons along the X side for each (A., B., C., D) of the first 4 weeks. Rather than 4 more figures, the 5-8 weeks are written here to reduce the sizes and the “only” UV mulch is not provided since all the rest have it to compare. Here are the numbers of the UV mulch left for each in percent: Burlap: 76, 46, 56, 30. Yellow: 75, 80, 70, 70. Green: 75, 84, 2, 58. and Camo: 80, 82, 70, 80. The opposite of course, is the difference of the UV mulch vs the treatment.

**2.15.5 Results:** This experiment was like some of the previous experiments discussed above. Those experiments were set up differently in terms of the arrangement of the treatments as well as without the addition of burlap in a similar role as the yellow, green and camouflage. It is also important to remember that the previous attractant/repellent treatments were tested with and without lures on both traps and fresh bolts of trees. The use of the ethanol lure here on the top of the UVMT attracts the beetles in numbers much higher than to trees or traps without lures and UV mulch. The figure relates the results using the relative percentage caught by each treatment to give a reasonable mathematical result of the treatment effects on the beetles (Fig. 32). The results shown for yellow and green paint, and camouflage indicate that ambrosia beetles, here specifically *X. crassiusculus* and some unidentified species, were not attractive/repellent, as the UV mulch percentage numbers were much higher. However, the burlap treatment unexpectantly increased the number of beetles collected over and above the UVMT control several times. The response to the color of burlap and a burlap matching paint (Fig. 32), both of which are very similar to a tree trunk, likely present a mechanistic comparison between ambrosia beetle’s responses to the light reflecting UV mulch and a dull colored ‘fake’ tree trunk (see discussion).

#### 2.16: Another approach: efficacy of UVMT on behavioral response of conifer-feeding insect species and general lures

**2.16.1:** Millar et al. (2008, 2013, 2015) have reported information on the relationships of trap type, lure type and other factors involved in the monitoring of ambrosia and bark beetles and their related behaviors. Up to this point everything reported herein – *X. glabratus, X. crassiusculus* and unidentified species - has dealt with the use of UVMT for only ambrosia beetle species that use hardwood species as host plants. In this section, for the first time, the efficacy of the UVMT in capture and monitoring of similar bark and related beetle species as well as other common insects found in both hardwood and conifer host trees that were reported primarily in Millar et al. (2013) were tested. The same lures were tested with the traps which included ethanol, ∞-pinene, ipsenol and ipsdienol. To accomplish the experiments, a large black vane trap made of 4mm coroplast and a smaller vane trap built by the author and completely covered by metallic silver paint. In Aug.-Sept. 4 in Monticello, FL traps of each type were placed 30m apart in an area that was being slowly harvested leaving downed, injured and normal hardwood trees in the area. The 8 traps were counted once a week for 8 weeks to gather the results. The species collected were identified to genus by comparison to Millar’s previous work as well as the authors.

**2.16.1 Results:** Twenty-one species were collected over the 8 weeks for a total of 670 insects. Four hundred insects (68%) were caught in the black vane traps, and 270 (32%) insects were caught in the metallic silver trap. Ambrosia beetles, bark beetles, cerambycids, buprestids, ostomids, wood wasps, platypodids, *Dendroctonus* spp., *Hylobius pales,* Cucujidae, Meloidae, and some other common species that generally respond to cut or downed hardwoods as well as the lures used in the study.

**2.16.2 and 3:** As a follow up to experiment 1 above, the ambrosia-bark beetle response behavior to ethanol, ∞-pinene, ipsenol and ipsdienol, a second experiment was conducted as follows. Eight traps made up of two replications, each of a Lindgren trap in natural black funnels and 2 other similar traps with funnels painted metallic silver (all with 8 funnels/trap) were used. Four more traps – large black vane traps were used with 2 in the natural black coroplast and 2 others with the natural coroplast painted metallic silver (the equivalent of UV mulch). The traps were placed along the edge at random 30m apart in 2 blocks of 4 of a recently harvested area of ∼5ha of forest in Quincy, FL. The lures were the same in all traps as described for experiment 1 above. The traps remained in the initial area and counted once a week for 6 weeks. Traps were randomized into new positions relative to the previous week after each count to reduce potential location bias. A second experiment here (3 in below results) targeting *I. grandicollis* was also conducted for 6 weeks as in experiment 1.

**2.16.2 and 3 Results:** Experiment 2 and 3: Very few *X. crassiusculus* or *I. grandicollis* were captured in the area but the number of *Ips calligraphus* was adequate to provide good results. There were no significant differences in the numbers by trap types – treatment effect (ANOVA procedure, F = 1.59, Pr > F = 0.20). However, the numerical differences were as follows: (mean ± SEM) black Lindgren trap 15.4 ± 3.4, silver Lindgren trap 20.4 ± 4.2, black vane trap 25.5 ± 4.9 and silver vane trap 25.5 ± 7.8. Experiment 3: same traps and handling, the same pheromones as experiment 1 but targeting *I. grandicollis*. For *I. grandicollis* the trap counts were also not statistically different. Numerically, the mean ± SEM/trap over all dates, was 23.3 ± 4.2 for the black Lindgren trap and 39.9 ± 10.7 for the silver. The numbers for the black and silver vane traps were statistically the same at 30.4 ± 5.9 and 33.8 ± 12.8, respectively. As indicated in the results of both experiments, the behavioral attraction responses to commonly used UV mulch, i.e., silver traps, by ambrosia and bark beetles appear to be very similar. A novel finding that should certainly make the reader stop and consider the implications for beetle behaviors when outside the tree (see discussion).

#### Part 2.16.4

##### Results: Field comparison of treatments selected by *X. crassiusculus* using UVMT

Four separate blocks of hardwood forests in Jefferson County, Florida were set up using 5-6 different treatments in each block with traps placed 30m apart. As described above, UVMTs were made of 21cm square boxes 0.9m long covered with UV mulch and attached to a tomato stake placed in the ground. The treatments were set up so that 2 opposite sides of each trap contained UV mulch and the other two sides contained only one of the 5 treatments compared. The control traps had all 4 sides covered with only UV mulch. Treatment traps were covered with burlap material, yellow paint, green paint, camouflage material, or matched burlap-colored paint. All 4 sides of the traps were covered with a 21 x 27cm piece of overhead transparency plastic and lightly covered with Tangle trap. A lure containing ethanol (AgBio, Inc.) was placed on the top of each trap. The experiment was initiated in the first week of March and traps were counted once every 7 days for 8 weeks. Counts were made of *X. crassiusculus* and any unknown ambrosia beetles (very few) that were trapped. The traps were cleaned as well as possible and were turned in one cardinal direction (1/4) after each count. The burlap paint was only tested during the last two weeks as shown in the figures. This is the first and only testing of green, burlap material and burlap-like paint in this text. Count data from each week was added together from each trap by date, block, and trap type and then the mean number of beetles captured per date, per block per trap by treatments were calculated. The mean total numbers of *X crassiusculus* from each or the treatment pairs were turned into percentages of the totals which appear on the figures to show the level of comparisons.

### Part 3: Understanding the interactions between ambrosia beetles and their potential host plants

**3.1:** In this final section, the many years of observations of ambrosia beetles, in particular *X. crassiusculus,* will be summarized along with other results with the purpose of raising some questions and potential results not in literature. There has been much research that in various ways have used the term “stress” together with the interactions between ambrosia beetles and their host plants. The next data is portrayed in Fig. 33 to show the distributions of *X. crassiusculus* that is common in early spring in plant nurseries in north Florida. The figure caption indicates the details of the trees’ developments. In this commonly observed situation of Bradford pear on the “contagious” side, the attack pattern has fewer trees that were attacked, but most of the individual trees had high numbers of attacks (Fig. 33). Whereas, the Drake elm tree attack pattern had a lot more, and in fact most of the trees attacked, but at low numbers of beetles/tree. The Bradford pear trees were usually killed, whereas the Drake elms more often were not. Note: such trees were only attacked by the earliest and first generation emerging/arriving ambrosia beetles in spring and were not attacked later in the year by proceeding generations.

**Figure 33:**
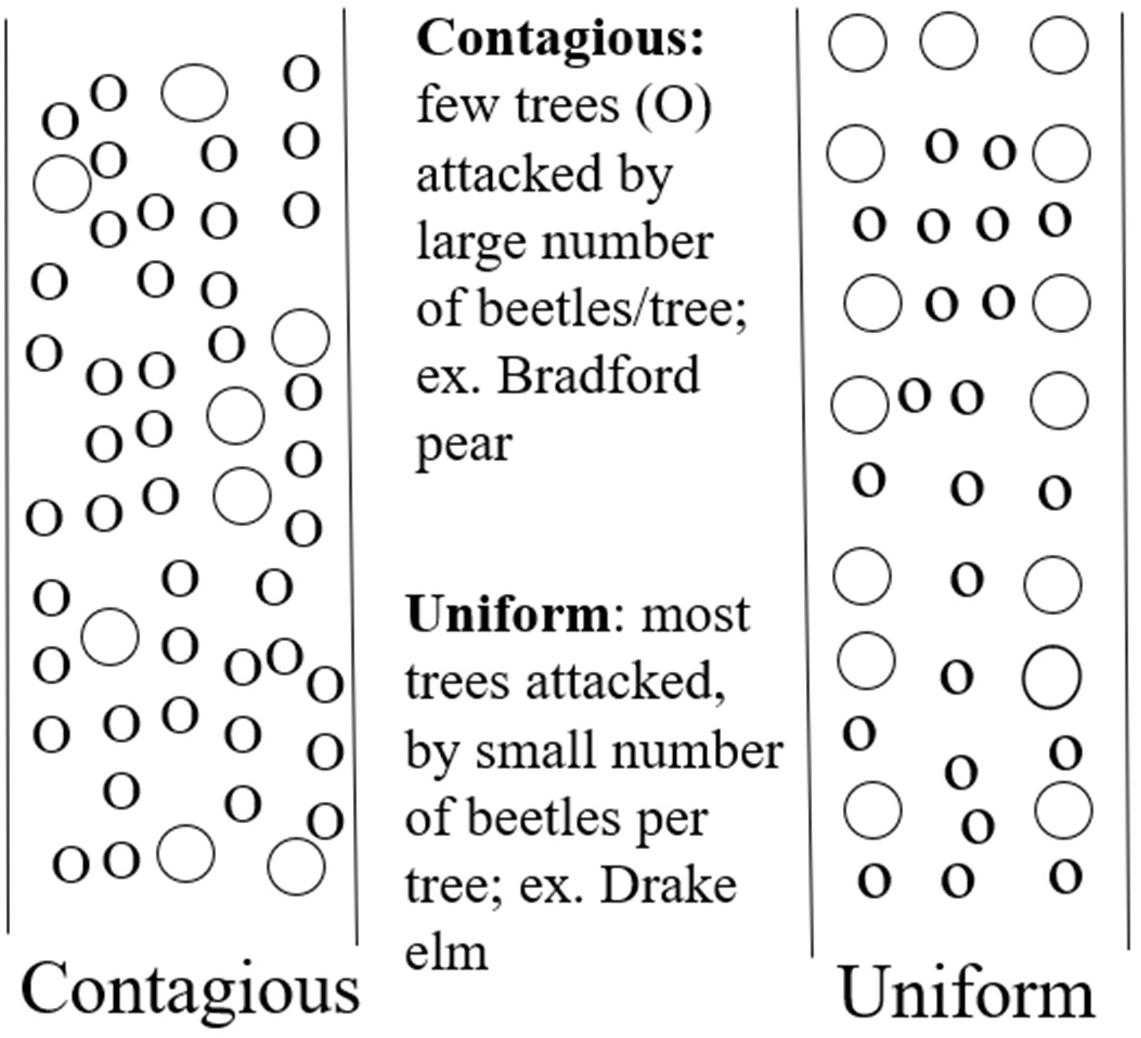
The small circles here indicate trees that are not attacked while the large circles indicate an attacked tree in 2 different “patterns”. This is a commonly observed behavior of ambrosia beetles within host trees in north Florida in early spring. The “trees” represented in each side were located together for a year or more within 25m, are cloned material of that species with identical age, water irrigation, fertilizer, etc., along with being exposed to the same weather. Bradford pears were in 19l (5 gal) pots and Drake elms were in 95l (25 gal) pots, tree size/within species is similar based on circumference at DBH. The figure compares two types of observed attack patterns. In this observed situation of Bradford pear on the “contagious” side, the attack pattern has few trees that were attacked but most of those trees had high numbers of attacks. Whereas, the Drake elm tree attack pattern had a lot more, if not most trees attacked, but at low numbers of beetles/tree. The Bradford pear trees were usually killed, whereas the Drake elms more often were not. Note: such trees were only attacked by the earliest and first arriving generation of ambrosia beetles in spring and were not attacked later in the year by proceeding generations.

The following Fig. 34 is provided as an explanation of the timing of changes in the host plant and beetles, in this case *X. crassiusculus* and *X. glabratus,* but redbay, elm and dogwood are also trees observed in a similar response that result in attacks by beetles. The model was drawn because of observed occurrences by the author and shows how plant phenology and physiology (equal safety from beetle attacks?) change over time “naturally” which results in susceptibility and nonsusceptibility to ambrosia beetles, e.g., *Xylosandrus* spp. and *Xyleborus* spp. It must also be understood that the insects are invasive species, non-native to the U.S. as well as plant species who are non-native to north Florida and the southeastern U.S. Moreover, *X. crassiusculus* attack trees which often produce ethanol while *X. glabratus* responds to specific kairomones produced by redbay as well as a few other related species.

**Figure 34:**
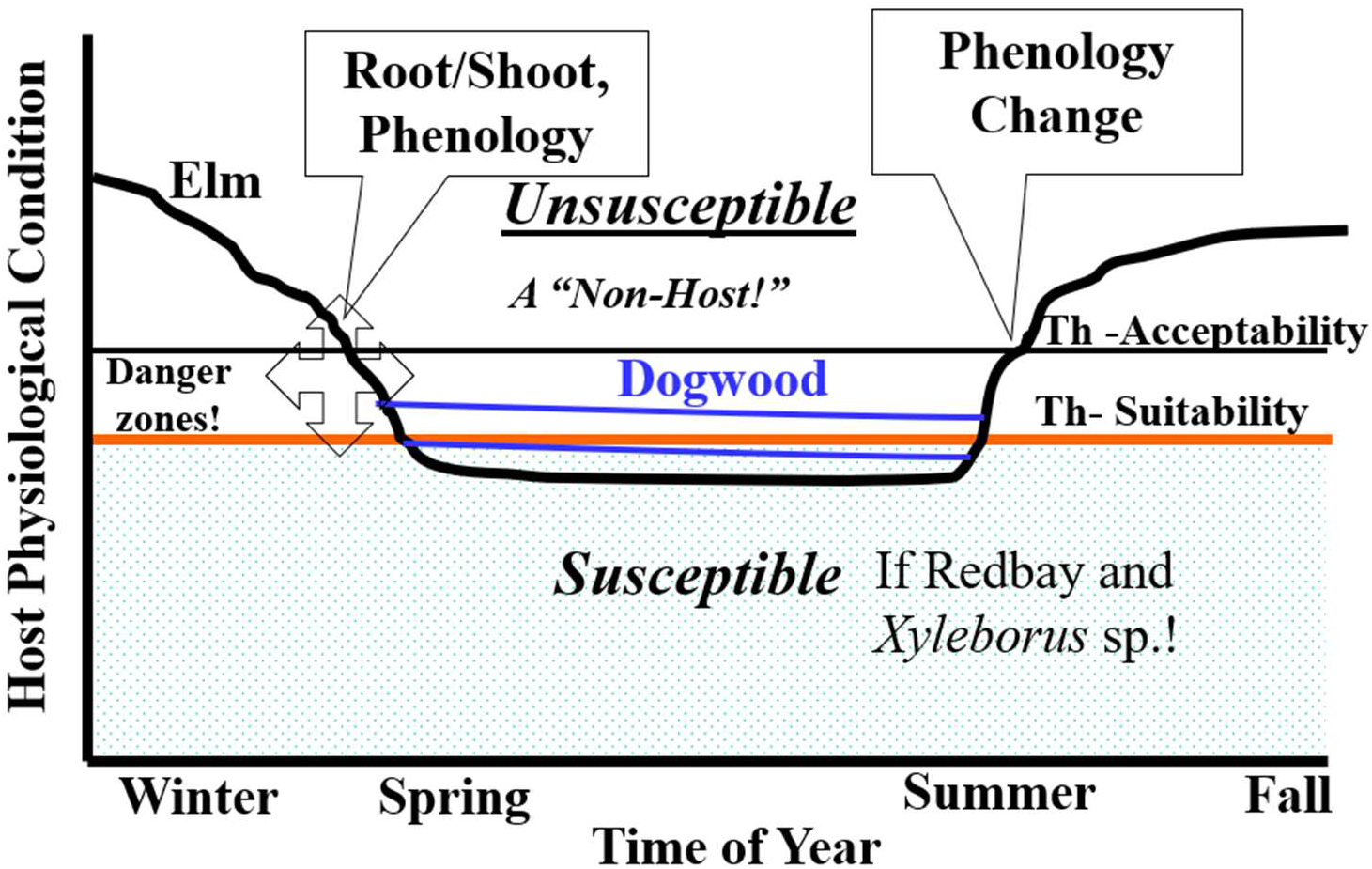
This figure is provided as an explanation of the timing of changes in the host plant and beetle’s (*X. glabratus* and *X. crassiusculus*) behavior in response that result in attacks. The model shows how plant phenology and physiology (= safety from beetle attacks?) change over time “naturally” which results in susceptibility and nonsusceptibility to ambrosia beetles, Examples: *Xylosandrus* sp., *Xyleborus* sp. In north Florida. See methods section for the in-depth explanation.

To understand what is happening herein, as is common, one starts at the bottom left of Fig. 34. The x axis is the time of year starting in winter to start the tree and beetle’s year together. The y axis indicates changing physiological conditions. The “susceptible” area (bottom in gray) indicates that redbay trees could possibly be attacked at any time but are prohibited until the weather warms up enough to activate beetles that overwintered inside of trees. This is a simple phenological/physiological change in a tree that will occur in response to higher temperatures that could be of short or long length in occurrence in any one year. See the example of 10 years from *X. crassiusculus* back in Fig. 1.

Step 2 in Fig. 34 is to look at the black line labeled “Elm” which also could be a number of other tree species. e.g., Bradford pear, mimosa, sweetgum, dogwood, etc., that have been observed by the author over the years and elute ethanol which is attractive to many ambrosia beetle species. This line moves out of the “Unsusceptible label” with “A Non-Host” suggesting that a plant species may not be a host at all, or at variable times of the year the beetles and the trees have not changed yet from the overwintering factors (low temperatures and change in phenology/physiology, etc.). Also, the Elm black line is buttressed at each end to indicate the beginning and end of the susceptibility of the trees (usually early spring) to the beetles, again phenology/physiological change. Step 3 is the parallel lines labeled on the left indicate the “Danger zone” and on the right by “Acceptability” top line and “Suitability” on the bottom line are 2 different interactions points between the trees and the beetles. The interaction area is very important to both the tree and the beetle’s interactions. Step 4, the “Dogwood” label and blue line are a special case. Throughout the year, but especially during the spring, trees such as dogwoods are often moved around from place to place in the ornamental nursery industries. Movements are often from northern latitudes to southern latitudes, e.g., Tennessee to north Florida which winds up pairing plants of the same species that have different weather exposures, and difficult physiology and phenology as a result. As a result, either group of trees may be attacked by beetles that, had they been kept in their area of origin, would not have happened. Dogwood trees that were handled this way were often attacked by beetles, but the tree physiology was such that the attacking beetles drowned in excessive tree sap. This was unlike the same trees that originated in north Florida that often were not attacked or were attacked at a different time or under a different growth situation (see Fig 34). This dogwood situation is common and is a good example of the interaction between different trees and the attacking beetles that occur in the 2 ways: acceptability (sometimes dogwood – Yes) and suitability (sometimes dogwood – No). Thus, there are physiological mechanisms changing - not necessarily stress!

#### 3.2: Induction of tree attacks by ambrosia beetles in several ways toward “inducing” ambrosia beetle infestations in various trees

In an earlier lucky day using the herbicide Roundup against small mimosa trees, it was found by accident (C. Riddle, personal communication) that topping the trees around 45-50cm above ground and then covering them with Roundup using a ∼10% solution in water induced attacks by *X. crassiusculus.* As a result of that finding, other chemicals known to affect the physiology of plants were tested as follows. Prep® brand of ethephon used for cotton and tobacco, 55.4% A.I., Bayer CropScience Ply Ltd, 390-393 Tooronga Rd, East Hawthorn, Victoria 3123, AU. It is an ethylene generator and regulates plant growth which accelerates boll opening and preconditioning before defoliation of cotton. For an experiment, the suggested label rate of 2ml/L water equal to 0.54% AI of the chemical was used. The second chemical was cacodylic acid (dimethylarsinic acid). Stream Chemicals, Inc. 7 Mulliken Way, Dester Industrial Pk, Newburyport, MA, 01950-4098. This chemical in powder form was used in a 10.8 % water solution. On mimosa a 1.5gms /15 ml water, and also 10gms in ethanol as a 6% solution. Since the 1940’s (Mayo, 2024) turpentine has been collected from conifers and this arsenic-containing chemical has long been used in southern pines to cause trees to store their chemicals in the roots for a time and later after harvesting the timber are extracted and processed for an array of useful chemicals (see Bouwmeester 2019 and https://arboris-us.com/evergreen-solutions/pine-chemistry/).

Treatments for the tests were conducted on live mimosa trees, which is an invasive tree species naturally attacked by the invasive *X. crassiusculus.* The control treatment for comparison was made up of fresh pieces of mimosa placed in 10% ethanol as described above in several experiments. The objective was to determine when and if post treatment the ambrosia beetles attack the trees. Small mimosa trees similar to the sizes (less than 10cm truck diameter at ∼<50cm height) discussed in previous experiments above were used. All live trees were clipped (topped) at the same height above ground and then treated with the liquid chemicals by using a brush on the top to transfer the liquids over the cut portions. Four trees were used for each of the treatments along with 4 topped water-treated controls. Prior to delivery of the liquids, a 17cm piece of clear plastic was wrapped around the tree trunks at the halfway point above ground on the trees and treated with Tangle trap. All trees were visited each day post treatment for 2-3 weeks. The ambrosia beetles on the trees were counted, removed along with any other insects (Cleridae) as well as trash, etc. This experiment was tried several times but only the first experiment provided good results. Finding enough similar sized trees was one problem and lack of enough beetles in the test areas was another.

**3.2 Results:** The results from the first experiment went very well (Fig. 35) as the treated trees were attacked over different times (both beginnings and ends per chemical) after the application of the treatments. The results clearly demonstrated that live trees, at least mimosa, can be induced to become attractive to *X. crassiusculus*. All 3 test chemicals resulted in attacks by *X. crassiusculus,* but the responses were different when the attacks occurred post treatment, the number of beetles that attacked a treatment and how often the attacks occurred. Prep™-treated trees came under attack within 2-4 days with an early and later (11-12 days post) peak, numbers from cacodylic acid were very low on day 10 only and the highest numbers attacked the Roundup-treated trees over the last 10 test days. Also, several very small Cleridae individuals were found on the traps with the ambrosia beetles.

**Figure 35:**
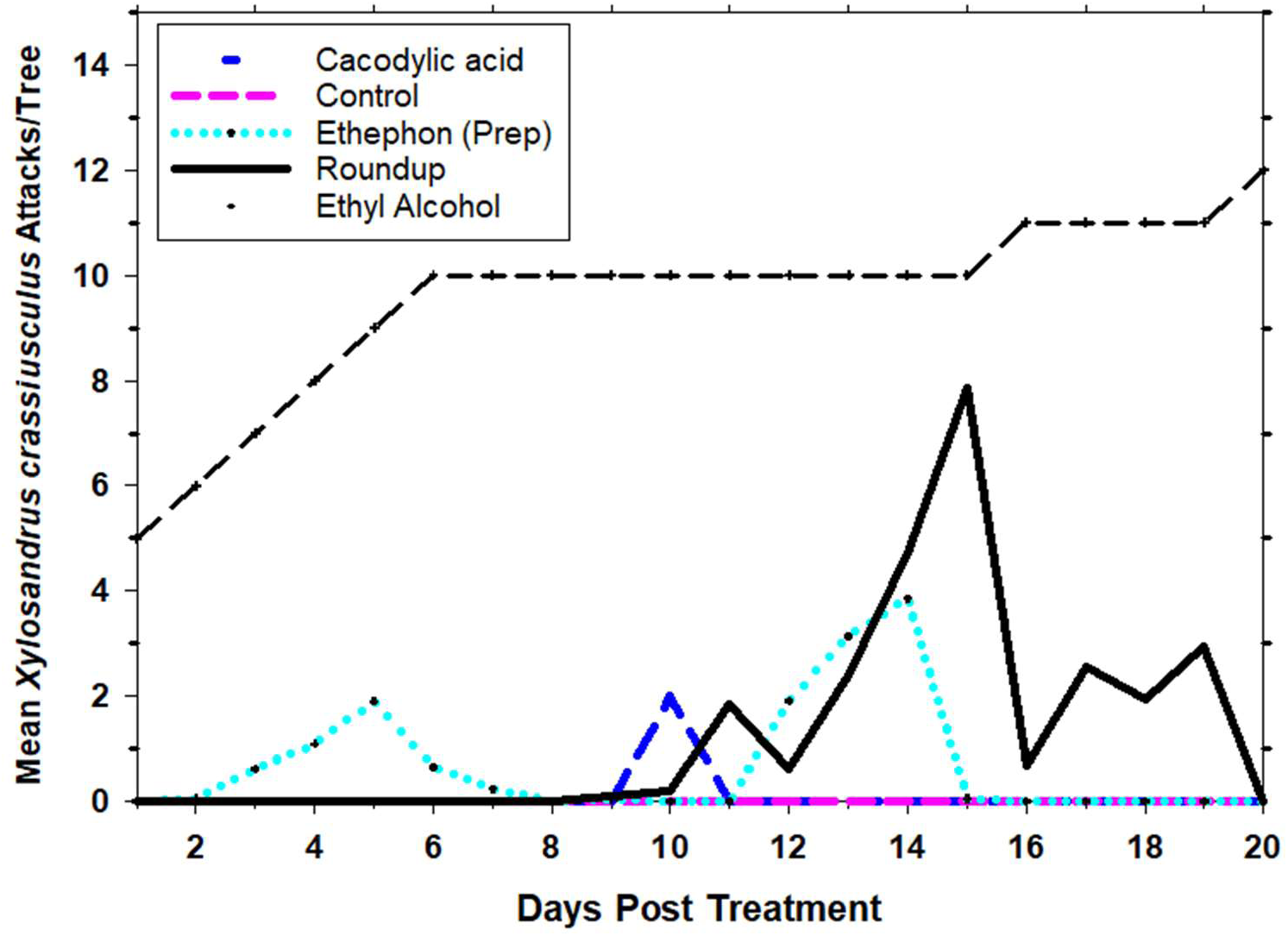
Mean number of attacking *X. crassiusculus* beetles/day over 20 days posttreatment. Use of various chemicals on “topped” cut trunks of mimosa trees that induced attacks by *X. crassiusculus*: chemicals and attack patterns. Chemicals of ∼3 ml were applied to the cut area on the trunk tip once using a small paint brush. Controls were topped but received no chemicals and the ethyl alcohol treatment (top black line) was a 10% water solution to 46cm long mimosa bolts in small cups placed in the area against wooden stakes.

From the results provided above concerning the behavior and its manipulation of ambrosia beetles (Fig. 33, 34 and 35), other experiments were attempted to further these ideas (Fig. 36). Some data developed over time are arranged and provided in this figure to group the main factors together as they were developed. Starting with the “Common” host plants box 1 (Fig. 36), with a list of known host plants on the left and the number of *X. crassiusculus* attacks per bolt as a measure of the relative preference on other potential hosts (Fig. 36). These values are the sum of the attacks on 6 bolts per tree type held in a 10% ethanol and water liquid as described above. The variation in numbers is surprising going from a high of 113 to the lowest of only 23 in mimosa which is an often-attacked invasive tree species as described previously in both live trees and cut bolts. These results paired with the results shown in Fig. 35 suggest a large number of trees may be attacked by this beetle and/or can be manipulated to do so. In box 2, a similar study using mimosa bolts, 3 plant essential oils vs a control using ethanol as above indicated that peppermint, cinnamon and wintergreen oil attracted 2-3x as many attacks by *X. crassiusculus* than ethanol.

**Figure 36:**
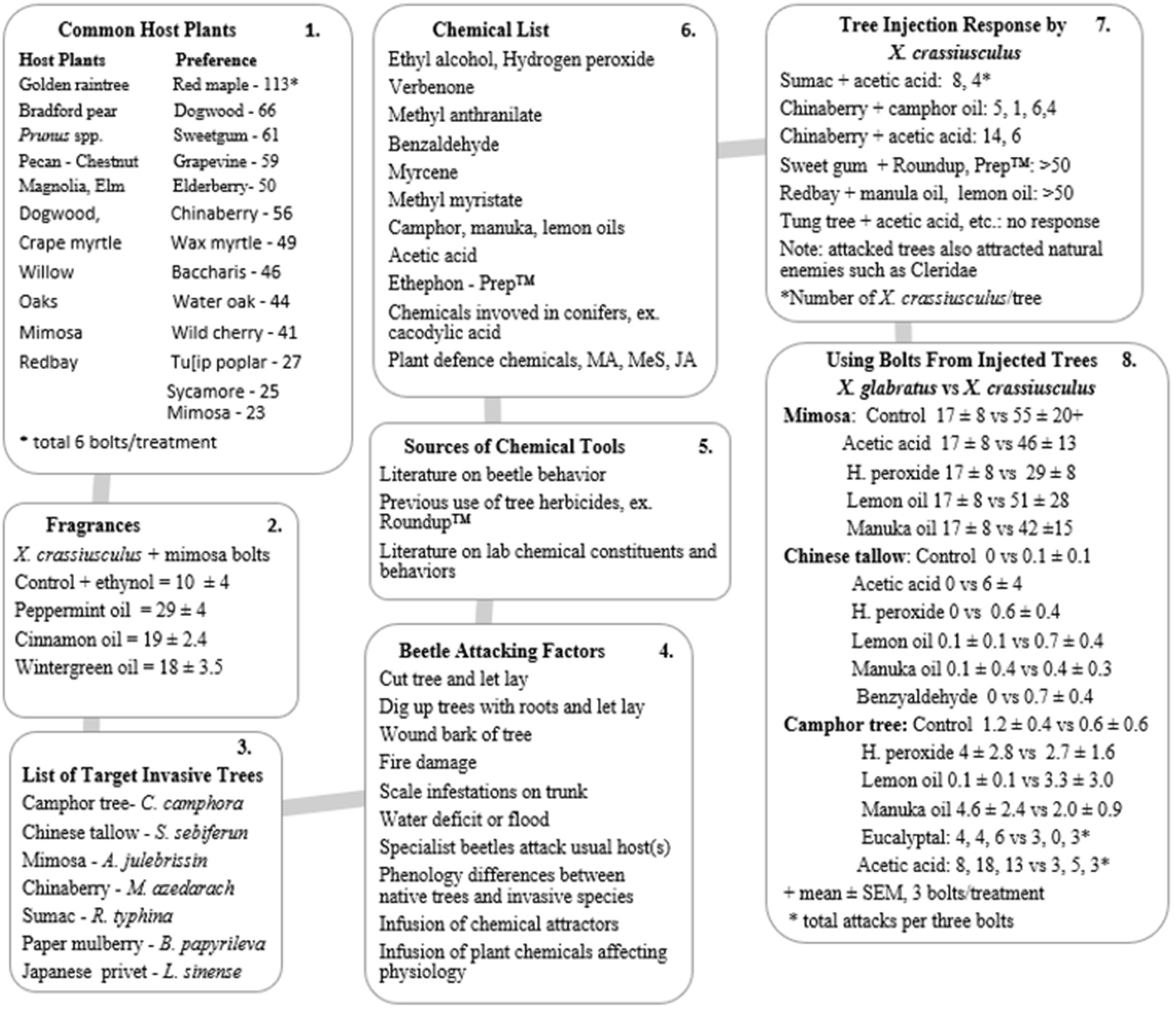
This figure provides a summary of the approach (developed over some years) toward chemicals and other tools used, efficacy, and results of an attempt to develop what I call “Hexapodacides”, e.g., manipulating and using invasive ambrosia beetle species to suppress and kill invasive trees potentially like herbicides.

As a result of these findings the name “Hexapodicides” was coined from ideas developed in Part 3 above towards the potential efficacy of using such a technique on invasive trees like the target trees listed (Fig. 36, box 3). The trees in the list are well known invasive, nuisance species, common in Florida and other places in the Southeast. To execute some experiments, a list of factors related to ambrosia beetle response to host plants were developed (Fig, 36, box 4). Note that there are some new methods listed that the author has observed as well as tested and discovered. These are methods that can be used to induce attacks by various species of ambrosia beetles. Literature was searched for more chemicals for potential use (Fig. 36, box 5 and 6), experiments were executed and the results are presented (Fig. 36, box 7 and 8). Two methods were used and the first was the injection of various chemicals into living trees using a drill bit to make 3-4 holes in the bottom 31cm of the tree trunks. The holes were made using a 1-2cm sized drill bit, held at a 45-degree angle into the wood toward the bottom of the tree trunk. Hole length was dependent on the size of the tree trunk but usually less than ∼10cm. Chemicals were placed into holes at ∼2-3ml and then the holes were plugged immediately using modeling clay. Once set up in the field, the trees were monitored over 5-10 days or until beetle attacks were obvious due to fresh frass on the trees. Method 2 was similar to treatments of the live target trees, similar experiments, e.g., using the same delivery approach and chemicals, were treated to tree bolts (Fig 36, box 8) by cutting the tree bolts as above and injecting them similarly as the live trees. It was used to follow up on the “preference” results data from Fig 36, box 1. The injected bolts were then set up in water-filled cups to evaluate the response of ambrosia beetles as described previously. The response of *X. glabratus* and *X. crassiusculus* are reported (Fig. 36, box 8).

## Discussion

### Part 1

This paper carries a lot of subject matter across a number of years gathered by the author. Climate change is related to a number of outcomes unexpected or in the process of changing much faster than observed some years before. As time passes in the future new insects such as ambrosia beetle species will find there way to Florida and other areas of the Southeastern U.S. In preparation for such, several sets of data gathered years well before climate change appeared to most foresters and entomologists in the represented areas. The data on *Xylosandrus crassiusculus* populations in different habitats at different settings along with 10 years of spring emergence by *X. crassiusculus* are offered as a longterm example of these original observations over 1995-2005 (Fig. 1). During the above observations in Fig. 1, efforts to find effective insecticides were also commonly tried. In Figs. 2-5 are the results of combining the repellent DEET along with and separate to insecticides alone over eleven test days using fresh mimosa bolts. The 4 experiments were conducted in 2 different locations around 50 miles apart. It was found that *X. crassiusculus*, the major ambrossia beetle present at the time, responded to the insecticides with and without 40% DEET alone or mixed. The treatments did not last much more that 5 days and were best once in each of the locations.

### Part 2

Prior to the discussion of part 2 and the new UV mulch related experiments, it is important to provide and recognize some of the previous work available in the literature relative to invention and use of a high number of currently available/used ambrosia beetle insect traps. Four publications are suggested due to their survey and detection of bark and woodboring beetles (Allison and Redak 2017), a more recent evaluation of the factors affecting catches of bark and woodboring beetles that includes a number of colored pictures of the traps (Tobin et al. 2024, Dodds et al. 2024) and a fourth very recent publication (Miller 2025) that evaluated colors on funnel traps in forest understories. With these 4 publications you can quickly get an idea of how their content may or may not relate to the UV mulch results that follow. There are also a large number of other publications that cover similar ideas related to beetle traps. I suggest that the other previous and current publications are evaluated as the UV mulch experiments and implications are discussed.

The main area of importance described is the finding of the behavioral relationship between ambrosia beetles and bark beetles to the well known UV mulch materials used to repel vectoring pests such as whiteflies and thrips from vegetables, etc. Note that there is only a little “color” reflected and there is low energy returns from green and yellow wavelengths which most people have assumed relate to forest plant leaves where ambrosia beetles occur and move among trees. Moreover, most entomologists early on consider the colors related to the insects they work on for a number of reasons, one being “could it be used against the target insect as an attraction”, especially for a pest insect. The original idea here was to find a repellent for ambrosia beetles and that led to testing UV mulch. The results of testing UV mulch and a similar metallic silver paint indicated attraction of UV mulch by the insects *X. crassiusculus* and *X. glabratus* and also many others. Fig 6, 7 and 8 provide the reflection spectrum of this well known ground-covering farming tool. Once these results presented by UV mulch and metallic paint were realized, other simple experiments were run to better determine how the behavior of the insects might be in North Florida and further exploited. The two major ambrosia beetle species available in the above paragraph were available and used in several small experiments with only statictical comparisons presented. Parts 2.1, 2.2 and 2.3 sum up the results of the 3 experiments along with some extra accompaning information provided (supplemental material Fig. S). In terms of beetle numbers, in these experiments, the redbay beetles had much higher populations at the time. When any traps were tested ∞-copaene and/or ethanol were placed on wooden stakes at 1.2m high and 1m away from targeted redbay trees.

The UV mulch and the silver metallic paint are very similar and potential use of these colors as a tool was of interest. With **Parts 2.2 and 2.3** as presented, the idea was to test redbay beetles for their response to “Fake” trees which were wrapped with UV mulch. The fake host trees worked by attracting both of the beetles – redbay and *X. crassiusculus -* along with other unknown species including some occasional Platypodinae. The followup tests using fake trees at three different distances from the target redbay trees indicated possible manipulation of the fake trees with UV mulch in combination with an attracting lure. Given the obvious potential effect of UV mulch/metallic silver paint as a tool, other experiments with redbay and redbay beetles along with *X. crassiusculus* and other unknown occurring ambrosia beetles were conducted.

### Part 2.4

This section is part of the evaluation to determine whether the UV mulch was visually attractive enough to attract beetles from host trees, in this case redbay was normally attacked. No lures were used. There were 3 sections that provided similar results on the level of tree deaths from ambrosia beetles. Fake trees, when without lures, attracted mean numbers of *X. glabratus*/week anywhere from a minimum of 22 in week 2 to 170 in week 3. The overall mean ± STDERR was 24 ± 9.3 percent of the total numbers of *X. glabratus* caught/week on any fake tree or natural tree traps. This result indicates the strong attraction of UV mulch to invasive ambrosia beetles; at relatively high numbers and unexpected. Moreover, the trap numbers on non-hosts in this experiment were like numbers on the main host redbay, again indicating the unexpected attraction of ambrosia beetles to UV mulch even from small size traps and ones without emitting normal odor attractants. The results were gathered somewhat differently but indicated the same results as expected. With the three experimental results, the potential impact of the UV mulch for protection of host trees appeared very high, supported the finished experiments, and made it well worth continuing to study its uses!

### Part 2.5

This experiment was conducted to determine how the redbay beetles respond to the trunk size of live redbays that were covered by UV mulch. The results of the indicated experiment are provided in Fig. 9 and 10. Interest in such an experiment emerged from the idea that UV mulch in some way might be turned into a tool (note, use of the “fake tree” in same way as mentioned above) and that optimizing such a tool might be important toward optimizing the size of the tool based on the behavior of the ambrosia beetles. The results of these experiments using redbay beetles indicated that with UV mulch as the “visual lure”, there was some size differents in the response. However, the beetles responded in high numbers overall despite the various trunk sizes.

### Part 2.6

From part 2.5 results on the surprizing attraction of ambrosia beetles to UV mulch, it was decided to further evaluate redbay beetles’ response to different sizes and shapes, etc., of fake trees which at this point could legitimately be termed “traps”. Four sizes of mailing tubes and building form tubes were compared that provided fake small to large trees and response to the trees via redbay beetles and others (Fig. 11, 12) (supplemental material F S). Further details are in the methods sections and pictures are placed as part of the supplemental information.

Combination of larger sized test traps indicated that cardboard traps covered in UV mulch were as effective as any tested. From these positive results other materials were considered to finish a final and cheaper insect trap that could be used in various new ways for ambrosia beetles, maybe other insects as well, and cheaper in costs. The details are provided and the final product turned out to be a (2 ft or 3ft) 62 or 91cm (preferred) long corrugated cardboard box attached to a tomato stake keeping it off the ground about 0.5m but placed within the ground ∼15cm for use in the field. The 4 equal sides of the trap were covered with UV mulch, lures such as ethanol were placed at the top of the trap or on individual sides (see cardinal direction ahead) and another piece of UV mulch of 20-30cm was added below the lure and covered with stickem to deteremine efficacy.

### Part 2.7, 2.8

These experiments were conducted to further test and enhance the UV mulch traps from several ideas but nothiing was changed. The square shaped trap was deemed the best.

### Part 2.9.1, 2, 3, 4

These experiments were to examine the apparent potential of cardinal directions which the square 20cm sized UV mulch trap provided. This feature was considered relative to the shape of the tubes that were decided as less effective. Moreover, the square traps present 4, well defined sections, which could possibly affect the beetle’s behavior on each side, thus enabling the ability to learn more from each trap. Furthermore, the sunlight and other reflective or absorbent factors as well as added chemicals (lures, etc.) (supplemental material F S) might be affected. For these reasons the square trap was renamed the UV mulch trap (UVMT). These results relate to why and how beetles might respond to UV mulch whereas many other leaf-feeding insect pests do not! In this case, aluminum foil reflects similarily to UV mulch without polarized light. Interstingly, Tabanidae are also an example of how other insects respond to light wheras in their case, polarized light is the driver (Horvarth et al. 1998). Note that the UVMT with 4 sides that can point in one of the cardinal directions, can be changed several ways: affixing treatments such that they are in different relative cardinal positions and each other on each individual trap giving at least 16 different combinations of relative treatments to move around and compare.

Along with the sections are the details involved related to using the new trap and its potential components that might be added such as colors, compounds, light spectra, materials, etc. This would optimize the data that can be quickly gathered and used (see Materials and methods (supplemental F S)). Also, previously the term “Fake tree” has been used herein, and coming up with that term here, will soon take a new meaning!

### Part 2.10

In this series of tests the 5 UVMTs had 4 sides including a control side with the lure and 3 other sides, one for UV Blocker, one for an absorbent chemical (Sport Sunscreen 50) and one for a sunscreen mineral reflectance chemical (Bare Republic Vanilla Coco). The methods and materials have the background of using such methods. Following the application of the treatments over each appropriate side of each trap, an additional piece of UV mulch in size indicated previously was placed over the middle third of the trap for each side and covered with Tangle trap stickem. The results in Fig. 13 and 14 demonstrate the response of the beetles, primarily *X. crassiusculus*, to be exactly as expected on the first day in order of high to low as reflectance 1, the control 2, UV Blocker equal to control, and then the lower amount of absorbent. The second day had very bad weather and in following days the reflectance was not as consistant, absorbent remained low and the other 2 more or less equal. Due to the problem with the reflectance malfunction, the data was changed to provide Fig 14 without the reflectance but its easier to see how the other three treatment appeared. This indicated very well that the searching beetles landing on the UVMT responded constantly over the 13 days of data as expected.

### Part 2.11

The potential for changes by beetles to the UVMT when outdoor lights were provided was tested. The square design of the UVMT along with the spread over cardinal directions enabled an experiment comparing the effects of with or without LED outdoor lights (Fig.15). No differences were found using *X. crassiusculus*, unlike the results reported by Gorzlanck et al. (2014).

### Part 2.12, 13

After developing the UVMT and testing some of its features as “tools”, new experiments related to other insect behaviors were conducted. At the beginning of part 2 above, comments about the occurrence and use of colors involved in insect behavior was a common activity of entomologists, especially early in the arrival of a pest species. In these experiments, yellow was a main target to begin, and as with other insect pests mentioned above, and also about the use of UV mulch against thrips, whiteflies and psyllids. Moreover, different yellow hues have been found to be in fact the best for use in a trap for certain pests including psyllids (Hall et al. 2010). As with previous data, colors have been evaluated early on under a number of cituations. Wearle et al. (2014) used 13 colors to compare in Tennessee and Mississippi the response of ambrosia beetles. Beetle capture using opaque (60) and red (54) were significantly higher than the beetles from yellow and white. It was decided that black traps were acceptible. In this part yellow hues were compared and tested using the UVMT as above. However, similar experiments will follow the trap data by comparing the colors and beetle responses on/to fresh wood.

From this “Part” on, the tests will be mainly based on 1. *X. crassiusculus* alone, and 2. other species that are termed “totals” and are the unknown species also counted but not identified. At this point, I refer to Conover et. al. (2026) as the first printed work with these ambrosia beetles and the tools mentioned here. It includes a list of approximatley 20 identified anbrosia beetles he collected in avocados in south Florida. Note that the publication used the tube traps I used before I changed over to the square box and the UVMT. Using the UVMT, light, medium light and bright light (safety yellow) colors along with a control UV mulch side were tested. The details are included in the figure sections. The data (Fig. 16, 17, 18 and 19) are 2 test approaches divided into 2 sections as explained, 16 = *X. crassiusculus*, 17 = totals (unknowns added into the first figure) and 18 and 19 are handled in the same manner. The explanations of the setups of the UVMTs used in the testing are also provided in the materials and methods. First, the capture rate of the ambrosia beetles in comparison to any use of yellow was clear and similar across the results. The UVMT with or without lures and both types, e.g., *X. crassiusculus* and “totals” attracted much higher numbers. Yellows were very low in four evauations, and overall results from the UVMT with 4 individual sides differed depending upon where and if a lure was used or not in that individual side. Moreover, with Fig. 18 and 19, the UVMT had 2 sides with no lures or yellow and those two sides started out the highest capture rate on the first day, then changed to the second side for 4 days rated highest. Additionally, the mean number of *X. crassiusculus* were kept lowest on the yellow side and it was the “total” traps that had a few different species that affected the patterns. Despite that factor, what resulted in this test initiated a whole new idea and the usefulness of theses experimental, practical results were began! At the same time, the conclusion presented, that the 4 sided UVMT provided a new and useful tool, was very much solidified.

### Part 2.14

This section continues the evaluations of the behavior of *X. crassiusculus* and relationships of the insects to real trees via fresh bolts. Much of the background is provided in the methods and materials section under 2.14. Fresh cut trees of mimosa (2 experiments), and sweetgum (3 experiments) are used within the next 5 figures – Fig. 20, 21, 22, 23 and 24. As above where the UVMT were used to compare responses of various combinations on the UVMT, these test trees received water and regular bark (control), ethanol (lure), yellow paint, and/or silver paint in a number of combinations. Also, as previously, the behavior in response to the factors tested are used in the forests with real host plants used by wild and free ambrosia beetles outside the tree looking for available host trees. There are very few differences among the results in the 5 figures. Regardless of the 2 types of host plant bolts, the controls and the yellows always had the lowest number and anything with ethanol and silver together or alone were highest. These results followed very well expectations that were provided from using UVMT and similar treatment combinations previously.

### Part 2.15

Given the positive results from the previous 2.14 part, the next part was developed to compare and contrast the behavior of enhanced attractions via ambrosia beetles to UV mulch and/or reduced response to yellows exploited alone or together enough to protect host trees, e.g., a push-pull? Also, if there are other patterns or tools such as yellow appears to be, that would add other related tools. Most of these related ideas are provided in the materials and methods section and a number of similar supporting factors are addressed similar to the evaluation of yellow. They are provided with the background data in a few paragraghs similar to yellow, which of course is tied to ambrosia beetle’s vision behavior. From there, camouflage in detail is presented, tested and compared to yellow in much the same way, thus the reason it was chosen. Here I will report the results from 7 figures related to the background information used by game hunters such as for turkey, deer, military or other such situations. Similarly, ambrosia beetles were used outdoors the way the experiments were conducted. Seven figures are provided (Fig, 25-31) to cover this unusual approach to insect pests’ behaviors. DPE and Mossyoak camouflage (camo) are 2 types of “camo” used for the experiments. While UVMT traps were used and compared for the information provided using the fresh wooden bolts, these 7 figures of data were very similar in the field development, handling, executing and presenting as well as were the results; e.g., the formations of the results in color and camo figures match very closely. The factors used such as the color yellow, sunscreen materials, absortion materials and UV Blocker material were also used similarly within the UVMT. Also, the handling of the figures used were *X. crassiusculus* means and totals (*X. crassiusculus* plus unknowns) since they were the highest number of ambrosia beetles available. As was stated within this paragraph, the results from the 7 figures using similar materials as previously provided along with the new camo results were very similar. The results prove that camouflage is responded to by ambrosia beetles with similar responses as provided from the color yellow. Thus, there were very low numbers of response to both factors. Much of the material is provided in the methods and materials section. Therefore, the figures can be read but here there will not be an indepth overlapping discussion of the data presented that is very similar to that of the yellow tests.

### Part 2.16.1, 2, and 3

In this next to the last of Part 2, the recognition of the related and mostly conifer-feeding bark beetle species will be minorly addressed with more data presented in the methods and materials. Most importantly, there are ample publications that are available for comparisons when this research was ongoing (Miller et al. 2008, 2013, 2015, 2025) and there are many others. Three experiments were conducted in north Florida with the commonly used forest entomology traps and lures, and pest species comparing the results of both types of forest insects. Note that the lures – ethanol, ∞-pinene, ipsenol and ipsdienol - were used along with Lindgren traps with 6 sections, black vane traps, and smaller vane traps made by the author covered with metallic silver paint. The details for the 3 experiments are listed in the methods, materials and results sections. As a first overview of part 2.16.1, twenty-one species were collected over the 8 weeks and a total of 670 insects. Sixty eight percent were caught in the black vane traps and 32% in the silver trap. The list of the collection is in the methods section and represent both types of ambrosia and bark beetles as well as a wide array of common forest insects.

For Part 2.16.2 the traps differed somewhat as stated in the methods. However, the trap numbers collected were not significantly different. On the other hand, the numerical difference between type of traps were somewhat different: black Lindgren lower than silver Lindgren and black vane and silver vane equal but larger than the first two. Part 2.16.3 was similar to 2 but collected a larger number of targeted *Ips grandicollis* as expected. The black Lindgren trap caught a mean of 23 and the silver trap 40. The black and silver vane traps were more or less equal at 30 and 34.

### Part 2.17

The next section of UVMT evaluations is summerized in Fig. 32 in the figures section. The letters from A-H (Fig. 32) are the data gathered and separated across week one to week 8. The experiment was initiated in the first week of March and traps were counted once every 7 days for 8 weeks. Counts were made of mostly *X. crassiusculus* and any unknown ambrosia beetles (very few) that were trapped. The traps were cleaned as well as possible and were turned 1 quarter (1/4) for one cardinal direction after each count. Remember that the square sides of the UVMT traps are 22cm wide with 2 sides for each of the two treatment types with the lures on the top and more open. This is the first and only testing of green, burlap material and burlap-matching paint in this text. Other factors are provided in the materials. The results are presented as shown to suggest the potential use of the information as an IPM tool to suppress and control the damage done by ambrosia beetles in crops and other worthwhile factors that receive significant losses, usually death. Also, Conover et al. (2026) presented a related successful trial with UVMTs that were tubes, although they were named otherwise in that paper that was evaluated on avocados.

At this point, the data will be evaluated using Fig. 32 details. Again, remember that the WVMTs provide 4 different sides to compare and in this case the number 1 “control” holds only UVMT on all 4 sides along with a lure, ethanol, on top and Tangle trap material on each side. Number 2, the WVMT has 2 UVMT sides and 2 of one of the other five treatments which are safety yellow paint, green paint, camouflage material, burlap cloth or burlap paint (latter only tested over the last 2 weeks). With this proven effective use of the UVMT and the setup in comparison of treatments, the effectiveness of each of the treatments can easily be determined. The colors match the treatments as do the relative sizes of the treatments versus the control, each other or any specific combination of interest, simply by comparing the number of insects attracted to each treatment. Lastly, the burlap cloth as well as a matched burlap color paint are quite different than UV mulch and represent two different ways that ambrosia beetles would visually respond to the 3 materials. UV mulch is highly reflective whereas the other two darker brown materials show little reflection and are close in color to a tree trunk. Therefore, these 3 colors indicate the two mechanisms by which ambrosia beetles would move through their environments: UV mulch provides the reflected appearance of an “open space” while the burlap mimics the tree trunks sought producing odors. Based on the provided information on the behavior of ambrosia beetles in positive response to UV mulch and burlap and repellency of yellow, green and camo, the correct answer as to what is going here is as follows. These small insects that spend a very small part of their lifetime outside of a tree trunk do the following. They leave the home tree, use their vision to avoid yellow and green foliage and anything related, e.g. camo added in this case, search to find the odor provided by the tree(s), follow it as a usual ambrosia beetle does to the emitting tree(s), land on the tree, smell and taste it. If the tree does not meet the need, they repeat until they find one that does or die.

Other potential factors: I want to recognize the contribution to Tobon et al. (2024) on testing trap designs and lures for similar ambrosia beetles. The results here in this section are predominantly about *X. crassiusculus.* However, on something as big as the change in understanding of major behaviors of an entire group of insects for the first time and the many species that were part of this research, it is highly unlikely that most don’t act the same as shown. Second, the colors that were expected to be responded to in a special way did not and now it is known why. Third, the combination of attraction to UV mulch and response positive to reflection and negative to absorbents, backup the “open space” attraction, identifies burlap as operating as an attractive tree trunk and identifies yellow, green and camo as unattractive. It turns out that the burlap presented as a tree trunk or “dead and dying tree” fits to a number of insect species’ behaviors including brown marmorated stink bugs (*Halyomorpha halys* Stal) which now have effective traps developed for the overwintering populations that is the subject of my next publication (see Mizell 2026).

Using IPM tools that may manipulate and control the populations of ambrosia beetles along with the UVMT in combination with yellow, green. camo or other available and similar-acting visual factors appear possible. The presented behavior and actions of the ambrosia beetles might be used to manipulate and manage these pests within a number of species and crops. As presented in Fig. 32, the number of insects were highly reduced when yellow or camo was used on either the UVMT or the fresh bolts of host trees. However, working the IPM tools together would likely be a more effective approach. Using the tools along with push-pull effects on such pests as pecan, many types of ornamental trees, avocados and such in spring would likely be much more effective and cheaper and is discussed in the next section.

### Part 3.1, 2, 3 and 4

The methods, materials and the final figures (Fig. 33, 34, 35 and 36) contain details of some observed interactions by the author between ambrosia beetles and their potential host plants. Most of the information is unusual and would likely be of best value if the sections were approached together. Thus, I am minimizing the comments in this discussion section on the last 4 parts since the other sections have most of what is needed. The observations of common behaviors by ambrosia beetles, primarily *X. crassiusculus* (Fig. 33), which were mainly the attackers of regular host plants in ornamental nurseries, occured repeatedly over many years as reported. Less assume here that there are some species that are more commonly and continuously attacked, others that show up as newly attacked trees have just arrived in Florida 14 or more days before via a truck from Tennessee or other colder more northern areas. Also, the trees involved all came from the very same background but some are attacked and others are not. Thus, the phenology of these plants have not changed soon (still in dormacy) enough or long enough to repel the attacking beetles. So one might question the suggestion that “plant stress” is the correct term for the problem. Of course. physiological change also plays a part in these insect-plant interactions.

For Part 3.2, Fig. 34 is presented as an effort to explain how the attacks by ambrosia beetles might be explained within the change in the occurance of tree phenology and physiology. The methods section along with Fig. 34 will provide an attempt to explain what could be occurring when beetles are searching for food.

For Part 3.3, Fig. 35 is presented as an additional report of how interactions among ambrosia beetles and trees interact when trees are chemically affected. Again the information is in the methods and figures section that explains what happened in this attacking response by ambrosia beetles. Here the use of herbicides initiated the method to induce ambrosia beetles to attack living but trimmed mimosa trees in north Florida. One of these chemicals, Prep™, is used in cotton defoliation in the fall. The results of these findings are attacked trees, and as long as you can find growing trees, new attacked trees can be made and used for whatever is of interest: beetle predators were also on the attacked trees for example. Also, some may remember that chemcials such as dimethylarsinic acid that are used on pine trees, wind up as sources of many useful chemicals when extracted from the roots of the trees (Mayo 2024). Perhaps mimosa or other tree species attack by ambrosia beetles are responding to important and useful chemicals also.

For Part 3.4 and Fig. 36, which is an unusal figure in shape and content, add more data to the use and occurrence of chemicals for induction of ambrosia beetles into trees for scientific as well as other reasons. First, there is an available paper from Florida (Wheeler et al. 2021) working with Brazilianpepper trees that also found similar results. Also, being that the target tree is full of milky sap, it is surprising that ambrosia beetles use the thick, sticky, milky liquid. Second, as the last subject in this paper, I gave it the name of what is being tried, i.e., “Hexapodacides” or “six legged killers”. That is - trying to manipulate “invasive” ambrosia beetles for the “greater good” and the induction to remove more invasive trees. The methods section and Fig. 36 hold the raw data that was developed in trying to test and use “induction’ to manipulate the beetles and kill invasive trees. The material and methods as well as the hard data along with tools are presented for further uses as the reader desires.

### A list of materials and some ideas that are presented and might be used to better maange ambrosia and bark beetles

Ambrosia beetles are important pests attacking many types of valuable trees. Recent results from scientific experimentation and field work by entomologists have led to new understanding and approaches that may become new and unique tools to manage these pests. The hope here is to make it come true! Not only do we need the potential tools to ask questions and to set up the new experiments, but we also need to evaluate the cost vs savings of the new tools, the difficulty of using them, time to maintain, ease of hand, speed and efficacy, etc. For those people that own the crops, etc., we expect to work with them up front.

## The Potential Tools

### Attractants

Liquids: ethanol, camphor (+30%), ∞-copaene only for redbay beetle, possibly conopothorin, increases for certain species along w/ethanol

UV Mulch (6’6” roll), metallic silver paint, aluminum foil (24” x 2000’, $68.00) along with ‘fake’ trees and various new combinations of equipment mentioned here

Colors: repel = yellow, green, camouflage. Attract = burlap (paint, materials in light brown).

Same as above and/or part of the mix as all below.

**Repellents:** (reduces beetle response)

Verbenone-10%, methyl salicylate-5%, piprotone, myrcene and comphene (half the effect of verbenone)? According to Ranger et al. (2013), a push-pull approach is likely needed

**Reflectance of light:** (increases beetle attraction)

Metallic-based sunscreens, added to traps (Bare Republic, SPF 50 mineral, non-nano zinc oxide)

**Absorbents of light**: (reduces beetle response, opposite of reflectants), chemical-based, regular sunscreens, SPF 50 was tested

**Insecticides:** 1. Legal chemicals that fit in with the types of materials that are still legal to use and commonly available. 2. The second kind, entomopathogenic and mycoparasite fungi as examples of several commercial strains, recognizes some work by Castrillo al. (2013) on *X. crassiusculus* using biocontrol agents. The cases show that *Beauveria bassinia* strains along with *Trichoderma harzianum* were virulent strains to the beetles and so were the *Matarhizium brunneom* materials. The scientists suggest that the fungi could kill the adult females of *X. crassiusculus* or could also be controlled by growth suppression or placing fungal symbionts in galleries.

### Traps

UV Mulch Trap (UVMT) with Tanglefoot (stickem) = lures or the similar but different UVMT that catches live insects, recently invented, tested by Russ Mizell; pictures in the supplemental area. These traps can be used very well as tools in the field to learn about beetle behavior. They are the basis for attempting these ideas to manage the beetles in a different manner. There are many other traps, available but not necessarily as good or better. Also, lures are needed to use with most of these traps although UVMT does attract some ambrosia beetles alone without an added lure or special trees. Also, remember that aluminum foil reflects colors same as UV mulch.

### Main Goals and Targets: Test and Use the Listed Materials to Protect Different Types of Trees Attacked by Ambrosia Beetles

“Individual” trees such as a few new ones planted in yard(s).

Groups of trees of different types are or will be planted in multiple areas using a list of trees that may be attacked by beetles early in the year. An example would be a newly planted portion of a pecan or fruit orchard or similar.

Large areas with trees of different sizes that are at risk from beetles at different times of the year, e.g., ornamental nurseries, new areas receiving newly obtained trees at risk, high value yards with developing buildings, etc.

Orchard crops with trees of various sizes that have product(s) of high value, are exposed to large numbers of ambrosia beetles, and often ones that are known to be especially at risk most of the year, e.g., avocados in south Florida. Most of the damage to crop trees is due to the attacks by beetles that infect the trees with fungi that the beetles consume. On the other hand, these beetles can also serve as vectors of pathogens that are passed along to other insects and increase the losses. To this point, avocado growers have lost $54,000,000 (T. Luginski, WUSF, 21 May 2024) in south FL from ambrosia beetles. There is strong need to introduce some effective tools in the near future for avocados as well as other crops that lose money from ambrosia beetles and others.

### Factors Related to Common Crops and Handling: Two examples

Avocados: Modern and high-density plantings: Row spacing: 4.5-6m (15-20 feet) is common in modern commercial groves. This spacing allows for easier harvesting, more efficient use of resources, and better management of tree size. It requires more intensive pruning and canopy management to prevent trees from outgrowing their space. Traditional and widely spaced plantings: Row spacing: 6-12 meters (20-40 feet) was more traditional. Benefits: Provides ample room for large, naturally growing trees and maximum light utilization. Management: Less intensive management is often required, as trees have more space to grow without overcrowding. Avocado spacing varies: 15 feet between Hass trees is manageable, 15 feet between rows with 12 feet in rows, and 18 feet apart in some. It is also important in having horticultural extension people help develop what is necessary to understand the needs. Peaches: General spacing guidelines: Dwarf trees: Space these 8 to 10 feet apart. Semi-dwarf trees: Allow 15 to 20 feet between trees. Standard trees: Provide 20 to 25 feet of space between trees for optimal growth. The mature canopy size of the tree is the most significant factor in determining spacing requirements with pruning, sunlight and air.

### Behaviors of the Insects That Will Need to Be Controlled

The target ambrosia beetles are females that grow up inside the host tree from eggs, feed until mature, mate, and then are prepared to find and attack a new host tree to continue the cycle. Note: we would be dealing with more than a few ambrosia beetle species here and there. It depends on the location being addressed. Conover et al, (2026) identified more than 15 different species after the collection with the original testing of the UV mulch traps in avocados in south Florida. There also was quite an amount of testing during the avocado work on an original push-pull test that can be found in (Conover et al. 2026). New ways need to be developed to use our IPM information against beetle behavior, e.g., to stop their “natural” tree searching behavior, and either use tools that make them stay away from the valued products (my new ones), make them kill themselves by getting them to capture themselves in baited traps, etc., or they are eradicated (maybe just a few could work) as usual with chemicals. This latter step could be used at the beginning and/or the end of the potential beetle manipulations.

Some of the new tools will be easy to develop, test and use. However, the major information about the larger areas will need interactions from other scientists and growers on how an individual avocado grove or fruit orchard could be managed effectively. A lot of details would need to be understood and effectively used here to be successful. Also, likely, there will be keen interest in the costs, which should be much lower than what is suggested under the different approaches listed above.

There are also two important terms to consider: 1. phenology, which means the times when the insects are operating through their natural life cycle wherever they occur, and 2. physiology, which is the term for the similar development of plants. Knowing the occurrences of these special times are key to understanding and utilizing the tools within the growth and behavior patterns of the target ambrosia beetles. See Fig. 1 with *X. crassiusculus* data over 10 years to better understand this and its importance. In north Florida, this type of data has been developed since I arrived in Monticello, FL in 1982 and covers insect behavior on crops like pecan, peaches, cold hearty citrus, tupelo honey, ornamental nursery production, etc.

Finally, the natural behaviors of certain insects can be quite different. One way to approach the problem is to use the three feeding types of ambrosia beetles and other insects. “Specialists” such as the redbay beetle (*X. glabratus*) are more closely related to insects that generally use individual or closely related host plant species and are the first attackers of such trees. “Generalists” (example, *X. crassiusculus*) and “opportunists” attack tree species later after the specialist beetles go to work (or occur due to other non-beetle tree problems) and where a few related species might use the trees. Also, “specialists” will usually be responding to different chemicals (lures) from different host trees they prefer. In the generalist species situation, tree resistance will be compromised in some way and produce general chemical attractants such as ethanol common across many species. Opportunists, mostly “decomposers”, will attack following up on the available trees changed by the two other damaging feeder types. These types of behaviors point to likely factors that could be used for research of the known species and their interactions as explained in above figures.

To handle ambrosia beetle problems, it will take some knowledgeable people with wider areas of expertise in each type of crop to develop and set up the required IPM tools discussed here. From a wider scale of thinking, one term is “landscape ecology” (now also farmscapes) which is what is present and how it works in what are known about the areas of interest as the “corridors” (where living things use to move about, also, pathways or field crops), “barriers” (unopen areas that inhibit organisms from movement, also, forest edges or interiors) and “matrices” (areas that are acceptable or not acceptable) and important for use by specific insect pests, also vegetation density or species. Many people don’t think of landscapes as structures. However, there are changes of distributions and abundances by life stage and time of year. “Trap crops” are the use and manipulation of plants along with their predators and parasites to control target pests. A publication for you that is related to stink bugs is available (Mizell et al. 2008). There is much involved around these subjects and the related crops, but these new tools can be effective if developed and used correctly (see the figures).

## Acknowledgments

This research was partially supported by a USDA-NIFA, Critical Issues grant with project number FLA-NFC-005067 in 2010 to the author, Peter Andersen and Henry Fadamiro. I thank Charles Riddle for outstanding technical help with early parts of the research and for discovery of the treatment effects on live trees with Roundup herbicide in 2005. I thank Thom Paris for help with the collection and evaluation of the spectra of various trap components.

