## Supplemental for "Discovery and implications of a novel, visual-attracting trap for wood-boring beetles (Curculionidae: Scolyidini): beetle response behaviors and underlying mechanisms"

**Supplemental Material:**

**List of Tools That Facilitate Working With Ambrosia and Bark Beetles:**

**Purpose: to fill in the experiences that many people gained in the past (but do not now) by growing up on a farm. (see pictures in supplemental material, not in any specific order)**

- A 4-wheel drive vehicle with an electric hoist, a 25-50' metal cable with a connection hook on the front if you are working in areas that can get wet quickly leaving large mud puddles and other hazardous problems that could lead to a stuck vehicle or a road closed by limbs or trees.
- A well equiped tool box with some of the listed objects below as well as simple tools such as screw drivers (both kinds), sharpeners and other accoutrements that are needed with other tools (e.g., chain saws, etc.), regular and pistol-grip staplers and staples in different sizes, hammers and assorted nails, screws, etc., and rolls of several colors of marking tape.
- A good camera to document everything important! Many phones have good ones now and carring a phone in the woods is a safety tool, but they are also often lost there. So add a fanny pack to keep it handy and safe.
- A good chain saw with enough power to cut the largest trees that you will work with along with a course in chain saw use safety, plus the required (by your institution) safety equipment. There is no more dangerous tool to operate than a chain saw!
- A very good health and safety kit in case something goes wrong in the forest, etc.!
- A sharp axe, large hatchet and/or folding hand saw for use in lieu of a chainsaw.
- Handsaw to cut smaller pieces of host plants.
- Drawknife – multiple uses in removing bark samples, wounding trees to increase attraction and smoothing of bark to make fitting of traps or other components tighter against the bole. See picture below in the supplemental material.
- Wounded trees – bark shaved in various locations and sizes on tree boles with drawknife.
- Infusion of tree bolts or injection of live trees with numerous other chemicals. See this publication part 3.
- Baited trees with attached lures with or without accompaning wounding, attractants/repellants.
- Visual cues added to bolts or limbs to compare treatments or other factors such as trap evaluation.
- Different parts of host trees either fresh or treated over longer aged periods after removal from the tree for evaluation of chemical composition over time, etc.
- Tree bolts or other parts infested with one or more species to determine potential attractions/repellency infestations from other species or congeners. Example: hardwoods and conifers will have different chemical profiles by species and over time (decomposition).
- Comparison of well known traps in different colors and at different heights with and without lures.

- A battery-powered drill with high power and a auger(s) bit for drilling holes in trees or soil to make or mount traps or for gathering wood samples, e.g., tomato stakes, see pictures for 2 examples. Also, in the supplemental materal picture there is a handmade metal digger that can be of whatever size fits your needs.
- DEET spray (or other brands and repellants) in no less than 40% concentration to avoid ticks, redbugs, ants, etc. Note: DEET also repels ambrosia beetles.
- Knives: a machete for making a path through heavy vegetation, and other smaller knives to use for addressing numerous problems. Snake leggins may also be of use in this chore.
- Shovels, small and regular sizes, for hole digging and other needs.
- "Shelf" shaped traps: see patterns for cutout of coroplast that have parts that catch either attacking or emerging beetles if coupled with a mesh-killing material such as insecticide-treated mesh, PermaNet® 2.0, (Vestergaard, Vestergaard Sàrl, Place Saint Francois 1, CH - 1003 Lausanne, Switzerland). See pictures.
- Camoflouge materials in different patterns (see this publication part 2) to determine insect visual response behavior.
- Sunscreen and a good hat to avoid or delay the inevitable multiple trips and needed skin treatments from a dermatologist.
- UV mulch materials now that they have been discovered as potential traps for ambrosia and bark beetles and as scientic tools.
- Liquid materials that affect sunlight or parts of the spectrum such as suncreens and U-V-Shield, U-V-Killer, etc.
- Stickem material for use on traps: there are a number of types and brands but the one that stays soft and is easiest to use, especially in cold weather is the Tangle-Trap™ sticky coating 'paste formula'. The Ortho Group, P.O Box 190, Marysville, OH, 43040.
- A regular staple gun and a pistol grip staple gun. Half-inch regular staples work best and cover needs for most jobs. The pistol grip is very handy in attaching things together.
- A number of bags and or cages that are 100% capable of containing your collected target species (or new species it has attracted) so that old or new pestiferous insects are not spread to uninfested areas unknowingly.
- Measuring tape(s): they come in different sizes, get what you need and make sure the color is easy to see under any circumstances if you drop or misplace.
- A collection of coroplast material with 2, 4, 10 mm thickness for making most anything needing to be flexible, waterproof, and easy to form with a razor knife, e.g., the "shelf" and other needs mentioned in the pictures section to place on tree trunks, and other places to monitor attacking or emerging insects.

**List of Ways To Induce Attack of Host Trees By Ambrosia Beetles**

- Phenological/phyiological manipulations: keep target trees cool or near freezing in the winter period to delay phenological related physiological changes. Then after the seasonal time passes for normal leaf development of the species used, take plant(s) out of the cooler and plant as normal; monitor for attacks.

- Use bark scales by hand infesting the trees so they can attack the bark of the target trees.
- Cutting off the trunk of a living tree, it appears to work best usually for smaller trees under ~10cm in diameter about 40-60cm above ground. Cut the trunk of the tree off neatly at ~45cm above ground and apply a chemcial lightly to the remaining cut trunk top that make the living trees attractive to ambrosia beetles. Chemicals that work for *Xylosandrus crassiusculus* and other species include Roundup herbicide, cacodylic acid, ethephon (Prep™ and other similar products used to remove leaves from cotton once it has matured), and ethyl alcohol. There are a number of other chemicals that will work in trees that are not "decapitated" so to speak. (see part 3 of this publication).
- Remove bark and phloem by shaving live tree trunks with a drawknife.
- Water deficiency or extended flooding will change the plants physiology enough to make it attractive to ambrosia beetles.
- Uproot an entire tree (roots included) and let it sit on top of the ground, many species attack these trees in a relatively short time period. Quick way to discover and start a lab colony.
- A naturally-attacked tree by ambrosia beetles of one "specialist" species usually attracts others in the area (see part 3 discussion). An example is redbay tree attacked by *X. glabratus* that prefers to attack redbay trees and uses kairomones to find the hosts also attracts *X. crassiusculus* and many other species that respond mainly to ethanol from suitable host trees.
- Effective chemicals that can change tree physiology and elution of attractants/repellants, etc., see Part 3.
- Ageing host tree bolts to change the elution contents and concentrations.

**Pictures and Uses of Some Handy Equipment:**

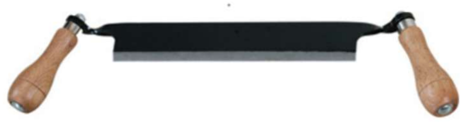

Fig. S1: Draw knife or draw shave knive useful for shaving bark finely off of trees and other jobs such as Mizell and Nebeker (1981) for determining the distribution of the pupae of *Thanasimus dubious* in pine boles

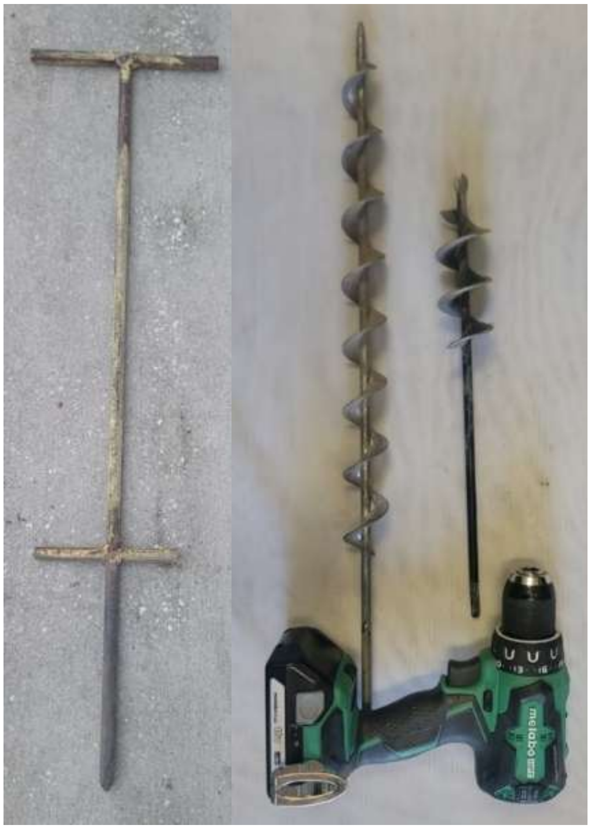

Fig. S2: An old fashion handmade metal foot-driven hole maker and of a strong battery-driven drill. Use with different kinds of bits to drill holes in the soil easier that closely fit whatever you are placing in the drilled hole that needs to have strong bracing against the wind.

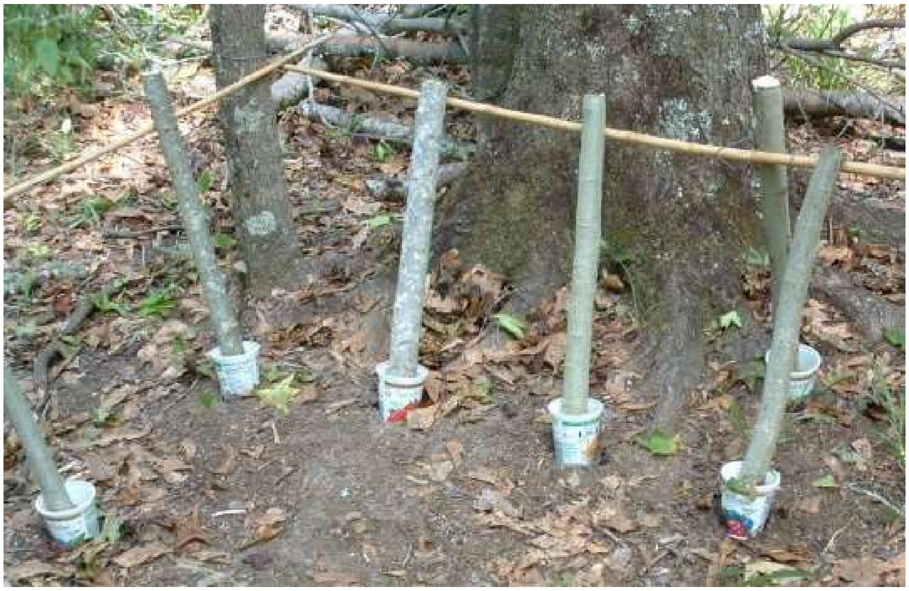

Fig. S3: Method that was used on the enclosed insecticide and other experiments with ~10% ethanol-water infusion of tree bolts provided in the plastic cups.

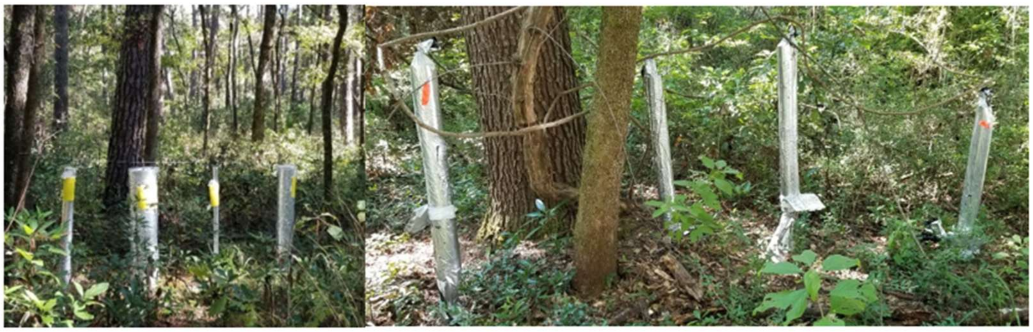

Fig. S4: What the earliest UVM Tube traps looked like in the woods after initial discovery and size tests.

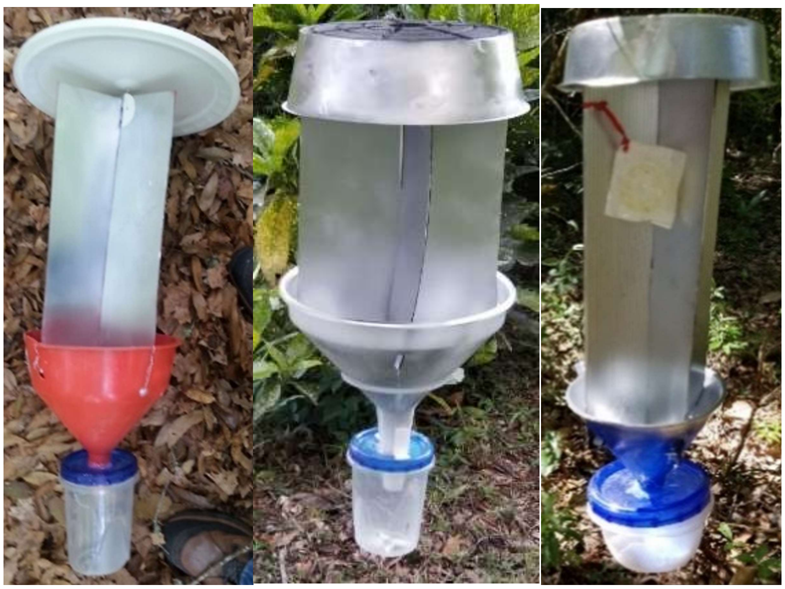

Fig. S5: Three vane traps I made and tested with UV mulch or metallic silver paint (the subsitute for UV mulch) for the capture of the very small *X. glabratus*. Left red trap has 21cm vanes over a 25cm funnel with 31cm wide lid. Middle trap vanes are 35 x 31cm over a 31cm “beer’ funnel and a lid made from a plant water holder also 31cm. The right trap vanes are 38 x 21cm over an 21cm funnel and 31cm plant holder lid.

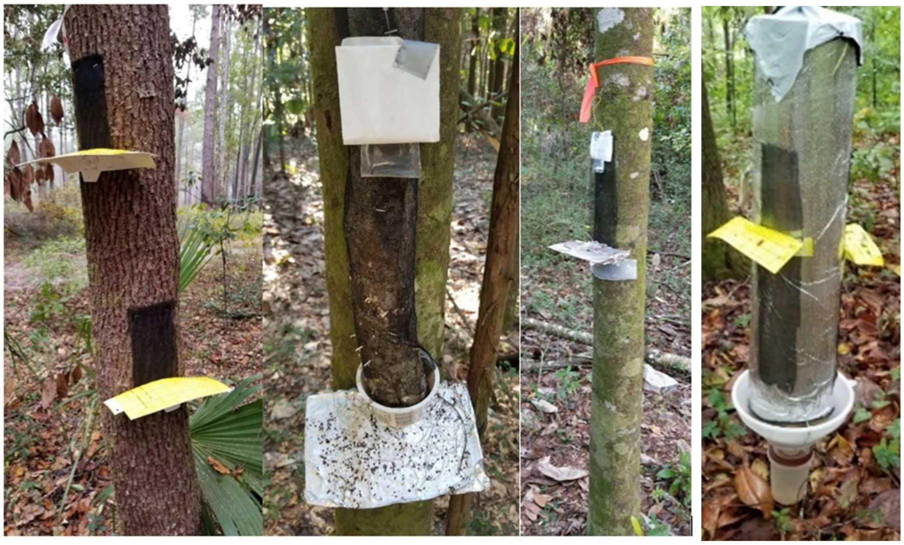

Fig. S6: This demonstrates some potential uses of Permanet™ insecticidal cloth with “shelf” traps on infested trees and a tube trap. Note: the lure can be used to increase the number of trapped ambrosia beetles since when they crawl out of the trees, they are looking for odors. Providing a line then may increase the capture on the closest trap? The yellow traps on attacked trees can be used since they come through an insecticide and have sticky material and squares to make count easier. Using a draw knife to smoothly shave the bark will help. Color may get in way on other uses of traps since some are repellent (see Part 1). The righthand trap is a Quik Tube turned into a multi-purpose trap.

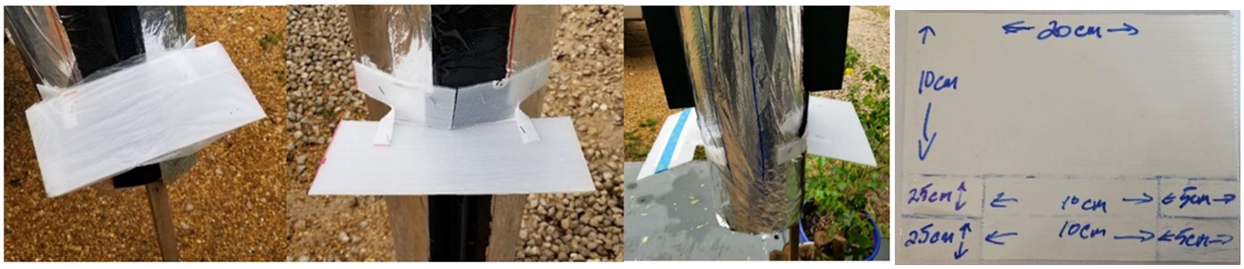


Fig. S7: Add clear overhead plastic paper with Tangle-Trap™ on it under the corners using aluminum foil and paper clips or the readily available commercial cardboard traps with Tangle-Trap™ that come in white or yellow as above. It works different ways as in the picture. Its easily mounted on most anything that will allow stapling. Meansurements for the shelf trap presented above right.

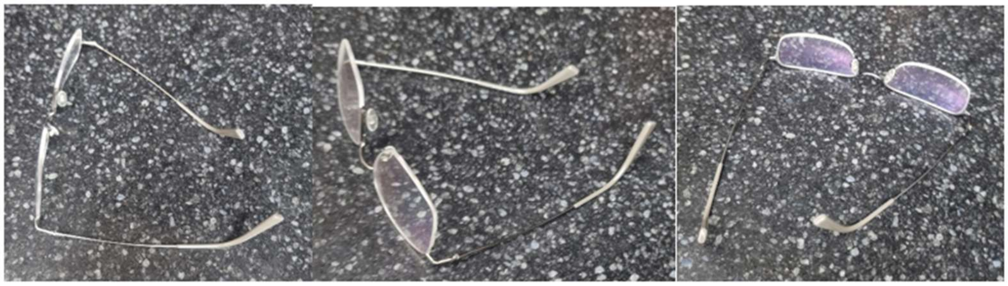

Fig. 8: Demonstration of a common example of the reflecting light changing the color on metal like UV mulch when at different angles.

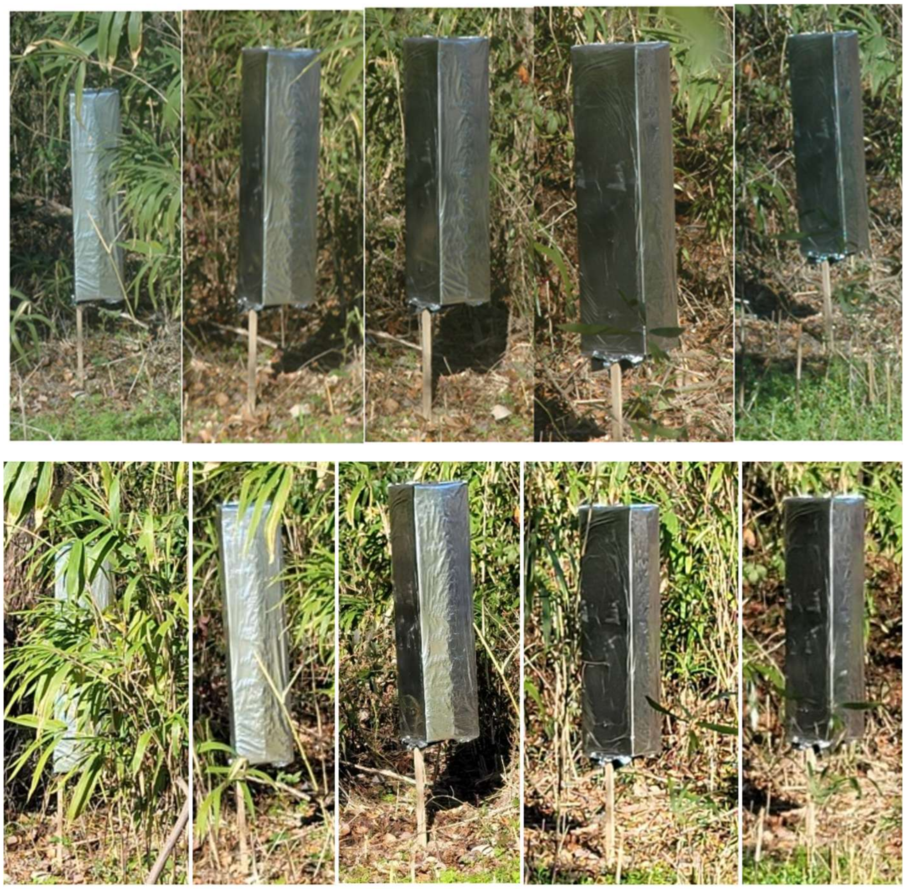

Fig. S9: Two different periods of sunshine indicating a change in reflectance of sunlight at different angles by UV mulch box-shaped traps. The angle of vision changes from left to right within the pictures and the sun is east behind the camera holder's movement. Top picture taken during a cloudy period, and bottom 2. a period with brighter sunlight. Both pictures were taken of the same traps at 10-11:00 am in an early summer day in 2023 in Monticello, FL.

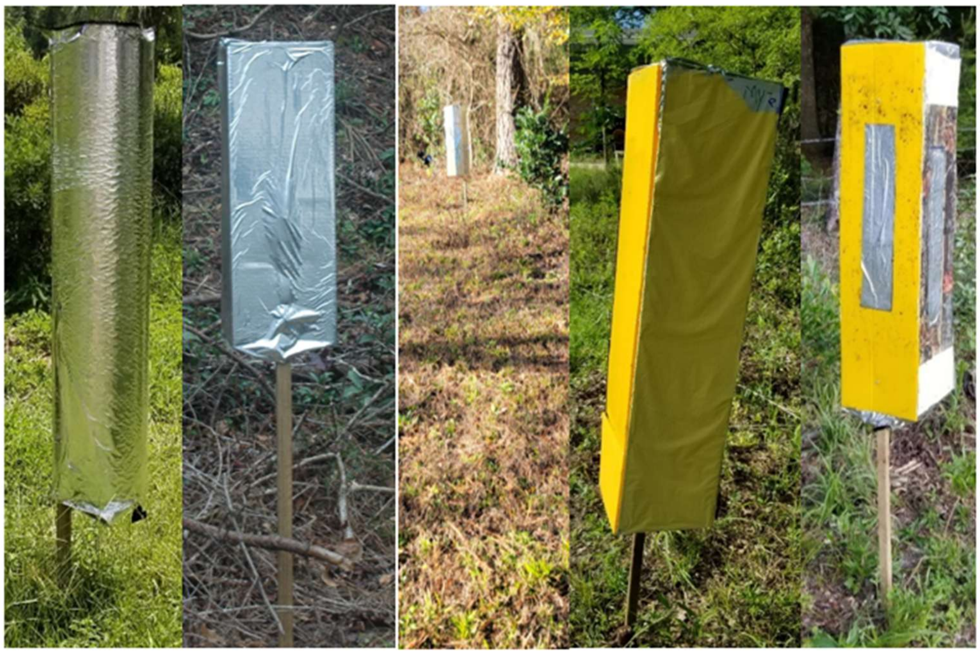

Fig. S10: a. The original Quik-tube trap covered in UV mulch,  b. the box trap of UVM, c. UVMT in cardboard shape reflection in early morning from the North, d. pure yellow, and e. yellow on one side with camo on another side.

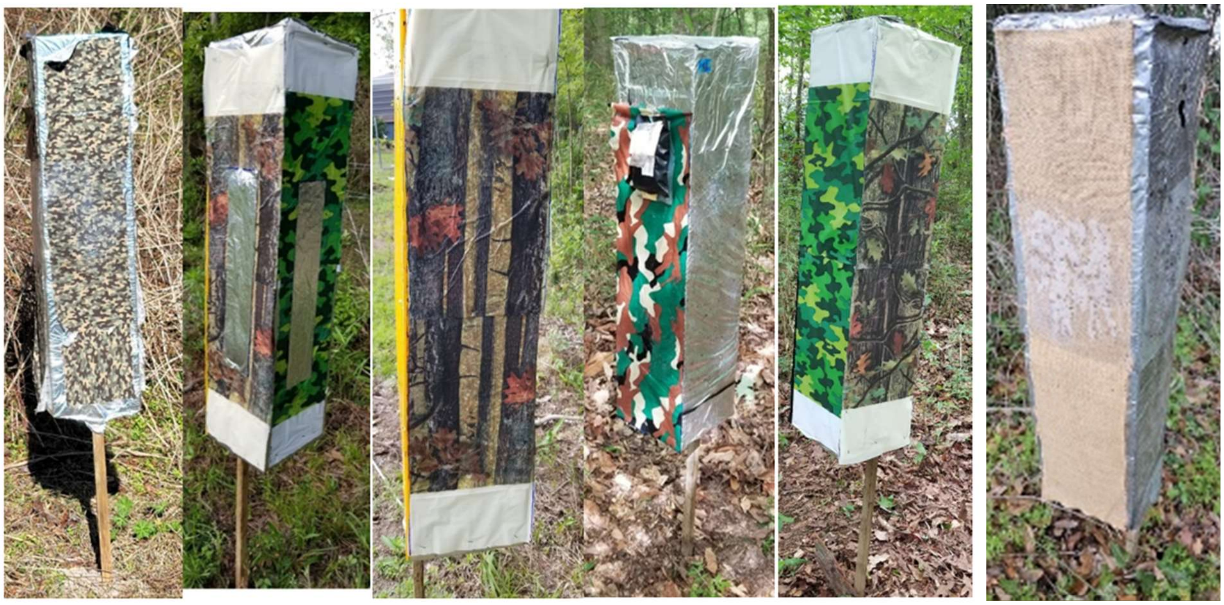

Fig. S11: Four types of camouflage on the UVM traps: a. (and c.) Marpat green digital camo from USADiscountFabrics via EPSY, b. Mossy Oak Break-up camo “Infinity” or “Country”, d. Oojami pack from Oojami store via Amazon, model number SYNCHKG063280, and e. also from Amazon, Inc. named camouflage party supplies. INeedFabric.com also has a large number of many different camo patterns. Note: the sources of camouflage changes over time and titles and numbers are not always provided by the producers. The fact is that most any pattern that resembles natural plants or backgrounds such as the above that resemble the plant leaves in colors, etc., are only responded in very low numbers to by the beetles on the UVMT.

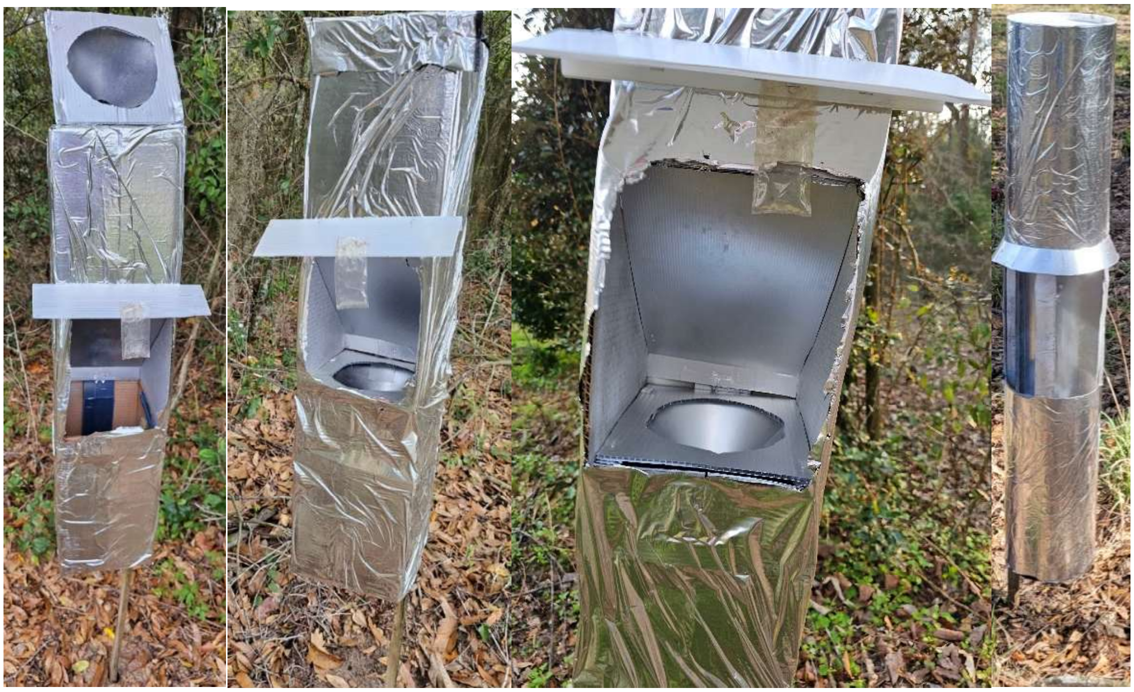

Fig. S12: Three pictures of the UV mulch trap built to enable capture of live ambrosia beetles along with the fourth picture showing the use of a tube trap in the same manner. The lure is placed close to the opening, where there is a small coroplast piece added to reduce the entry of rain water. Coroplast is also used throughout to mount a funnel inside with a catch vial at the bottom. The funnel is mounted under the piece of coroplast on top using other pieces of coroplast to place supports on the 2 sides of the wall as well as pieces under the hole to enable adding the funnel so that the pieces can be removed to count the insects captured inside.
